# Morphogen and juxtacrine signalling dynamically integrate to specify cell fates with single-cell resolution

**DOI:** 10.1101/2025.09.15.676088

**Authors:** Alicia Donoghue, Lewis S Mosby, Inês Lago-Baldaia, Tamara Hodgetts, Evelina Ursu, Zeynep Erten, Zena Hadjivasiliou, Vilaiwan M Fernandes

## Abstract

Morphogen gradients guide tissue patterning but do not act in isolation. How they integrate with other signalling modalities, like juxtacrine signalling, and how these integrations influence pattern resolution and robustness remain unclear. We address this in the Drosophila lamina, where columns of precursors are patterned with single-cell resolution into motion-processing neurons, dependent on a photoreceptor-derived Hedgehog gradient and glial-orchestrated differentiation. Combining experiments and theory, we show that glial-induced ERK activity drives *Delta* expression in lamina precursors, generating a graded Notch activity pattern. Notch restricts Hedgehog morphogen relay and enhances positional information. Glia act as timekeepers, scheduling ERK-driven differentiation after Hedgehog and Notch patterns are established. Thus, Notch and ERK dynamically integrate with Hedgehog to encode positional information, enabling reproducible cell fate patterning with single-cell resolution.

## Introduction

Developing tissues produce complex patterns of cell identities with remarkable precision and robustness, even in the face of fluctuating environments and genetic variation. Central to tissue patterning are morphogens, secreted molecules that form concentration gradients and direct cell fate by eliciting distinct transcriptional responses in a concentration-dependent manner (*1–3*). However, morphogens rarely act in isolation. Rather, cells are exposed to multiple signals simultaneously, which they integrate to inform cell fate decisions. Indeed, integrating opposing morphogen gradients enhances patterning precision, robustness and scaling (*4–8*). Despite this, how morphogens dynamically integrate with other signalling modalities, such as juxtacrine or other non-morphogen signals, and how such integrations impact pattern resolution and robustness remain largely unexplored.

To address these questions, we focused on the developing lamina, the first optic neuropil in the Drosophila visual system to receive photoreceptor input (reviewed in (*9*)). The lamina mirrors the spatial organisation of the compound eye, containing ∼750-800 modular units, or cartridges, each responsible for processing visual input for a single ‘pixel’ in the visual field (*10*); thereby contributing critically to motion detection (*11, 12*). The lamina is built progressively, starting with the assembly of lamina pre-cartridges, called columns, each consisting of six post-mitotic lamina precursor cells (Fig. 1A). Strikingly, each ∼4μm-wide precursor adopts one of five neuronal fates, L1-L5, or undergoes apoptosis based on its position within the column. Thus, cell fates are specified with single-cell resolution, repeatedly over hundreds of columns.

**Fig. 1:**
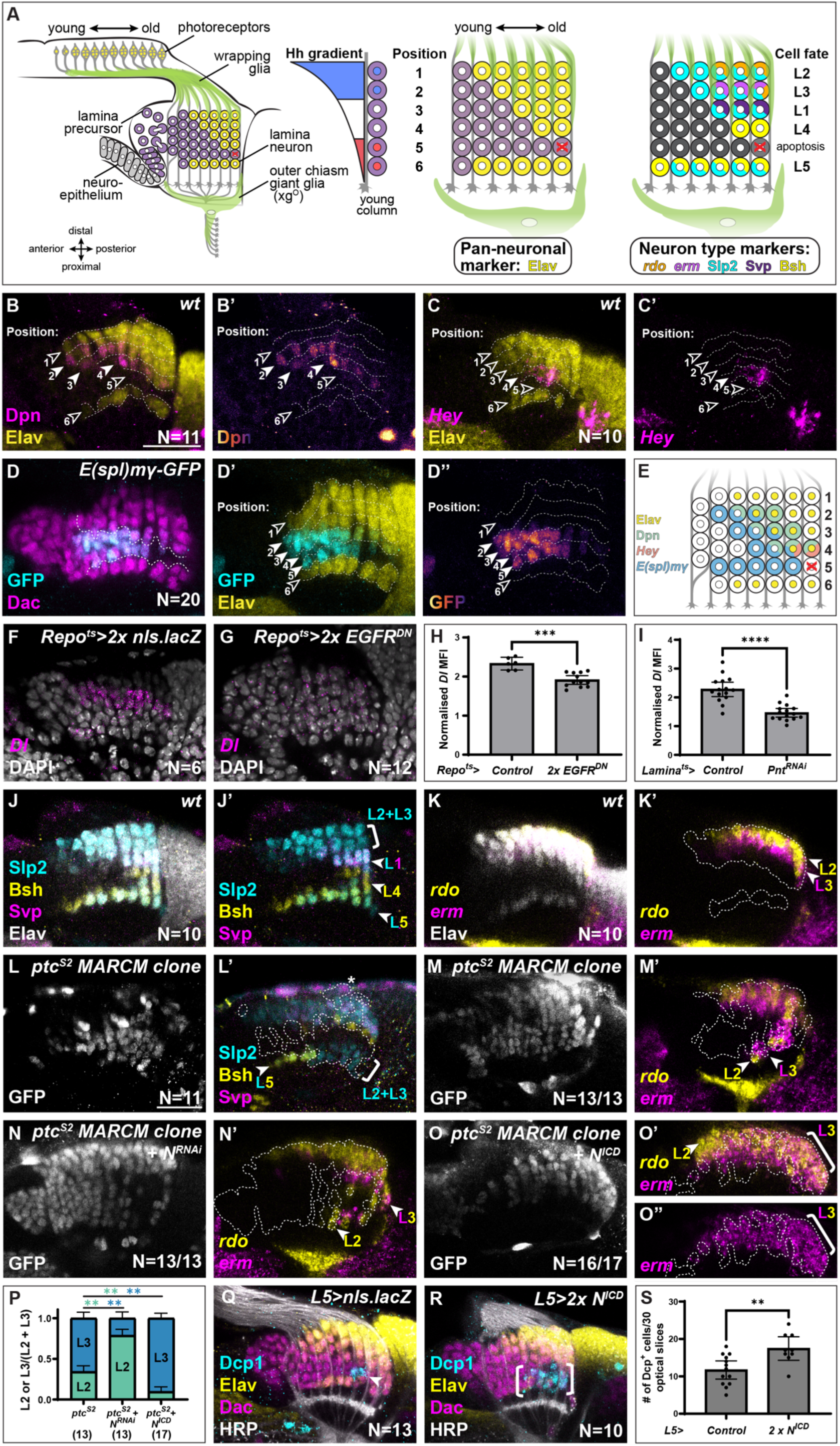
Glia pattern Dl-Notch signalling activity in lamina columns, with Notch instructing cell fate decisions in high and low Hh-responsive domains. **(A)** Schematic of the developing Drosophila eye disc and lamina, showing the organization of lamina precursor cells in columns of different ages, wrapping glia, outer chiasm giant glia (xg^O^) and differentiating lamina neurons. A photoreceptor-derived Hh morphogen gradient is polarised from high to low concentration along the length of young lamina columns, such that three Hh concentration-responsive domains are established. Wrapping glia and xg^O^ induce lamina neuron differentiation, marked by Embryonic lethal abnormal vision (Elav), a pan-neuronal marker. Each precursor will adopt a unique fate based on position in the column: L1-L5 neurons and an apoptotic fate. L1-L5 neurons can be distinguished by a combination of markers: *reduced ocelli (rdo)*, *earmuff (erm)*, Sloppy paired 2 (Slp2), Seven-up (Svp) and Brain-specific homeobox (Bsh), whose expression patterns are indicated. **(B-E)** Notch pathway reporter expression in the developing lamina with neurons marked by Elav (yellow): **(B)** Deadpan (Dpn; magenta and pseudo-coloured in B’), **(C)** *Hairy/E(spl)-related with YRPW motif (Hey*; magenta), **(D)** *E(spl)mγ-GFP* (GFP; cyan in D and pseudo-coloured in D’’) with Dachshund (Dac; magenta in D) to mark all lamina cells. Dotted line in D marks the boundary between undifferentiated precursors and Elav+ differentiating neurons. Cell positions are indicated with rows separated by dashed lines. Solid and empty arrowheads mark rows with or without detectable reporter expression, respectively. **(E)** Schematic summarising the expression patterns of Notch pathway reporters Dpn, *Hey* and *E(spl)mγ* for each position in the lamina. **(F, G)** *Delta* (*Dl)* expression (magenta, as assessed by HCR) with all nuclei labelled by DAPI (white) in **(F)** a control lamina and when **(G)** EGFR activity is blocked in all glia. **(H, I)** Quantifications of *Dl* mean fluorescence intensity (MFI) in the lamina normalised to the lamina plexus when **(H)** EGFR is blocked in the glia (F, G) and **(I)** when *pnt* is knocked down in the lamina (Fig. S1B,C). Mann-Whitney U-test, p***<0.0005, p****<0.0001. **(J)** A wild-type lamina labelled with Slp2 (cyan), Bsh (yellow), Svp (magenta) and Elav (white). L2/L3 neurons are Slp2+ only; L1 neurons are Slp2+, Svp+ only; L4 neurons are Bsh+ only and L5 neurons are Slp2+, Bsh+ only. **(K)** A wild-type lamina labelled with *rdo* (yellow), *erm* (magenta) and Elav (white): L2 neurons are *rdo+erm-,* whereas L3 neurons are *erm+* (irrespective of *rdo* expression). **(L, M)** GFP-labelled *ptc^S2^* MARCM clones (white; dashed outline) labelled with **(L)** Slp2 (cyan), Bsh (yellow) and Svp (magenta), and **(M)** *rdo* (yellow) and *erm* (magenta). The asterisk indicates a Svp-positive surface glial cell within the clone. Both L2 (*rdo+erm-*) and L3 (*erm+*) neuron types were recovered within the clone. **(N, O)** GFP-positive *ptc^S2^* MARCM clone (white; dashed outline) expressing **(N)** Notch^RNAi^ or **(O)** N^ICD^, labelled with *rdo* (yellow) and *erm* (magenta). **(P)** Quantifications of the proportion of L2 and L3 neurons out of L2+L3 neurons from (M-O) shown together. One-way ANOVA with Dunn’s multiple comparisons test, p**<0.01 **(Q, R)** Optic lobes stained for cleaved Death caspase-1 (Dcp1; cyan), Elav (yellow), Dac (magenta) and Horseradish Peroxidase (HRP; to mark axons in white) in **(Q)** a control and **(R)** when two copies of N^ICD^ are expressed in L5s. **(S)** Quantification of Dcp1+Dac+ cells in a 30µm section of the central lamina from (Q-R) (See Materials and Methods in Supplementary Materials). Mann-Whitney U-test, p**<0.01. Error bars represent 95% confidence intervals. Scale bars = 20 µm.

Hedgehog (Hh), secreted by photoreceptors in the developing eye disc, drives the assembly of lamina columns (*13, 14*). It also forms a morphogen gradient along the length of young lamina columns, which contributes to specifying neuronal identity by establishing three concentration-responsive output domains (Fig. 1A, (*15*)): high Hh signalling specifies L2 and L3 neuronal types (positions-1 and -2), intermediate levels specify L1 and L4 (positions-3 and -4), and low levels are associated with apoptosis (position-5) and L5 identity (position-6). Supporting this, we previously showed that dampening Hh signalling to different degrees favoured either L1/L4 identities or L5 identity, irrespective of cell position within the column (*15*). Whereas activating Hh signalling to its maximal physiological level specified a mix of L2 and L3 neuronal identities, rather than either identity uniquely, at the expense of other cell identities (*15*). These findings demonstrate that Hh acts in a concentration-dependent manner to subdivide columns into three domains, but that its gradient alone cannot account for the full range of lamina cell fates, and suggest the existence of additional signals that act alongside Hh to direct cell fate decisions with single-cell resolution.

Neuronal differentiation initiates and proceeds in a stereotyped sequence, driven by two glial populations: wrapping glia and outer chiasm giant glia (xg^O^). In response to Epidermal Growth Factor (EGF) from photoreceptors, wrapping glia secrete insulin-like peptides, while xg^O^ secrete the EGF Spitz and Collagen type IV alpha 1, respectively (*16, 17*). In turn, these peptides activate ERK signalling non-cell-autonomously in lamina precursors, inducing their differentiation into neurons (Fig. 1A) (*16, 17*). Wrapping glia ensheathe photoreceptor axons and progressively infiltrate the lamina from the distal surface, thereby driving a distal-to-proximal sequence of neuronal differentiation within each column (from cell position-1 through -4; Fig. 1A) (*16*). In contrast, xg^O^ are positioned beneath the lamina, do not infiltrate the tissue, and induce differentiation only in the most proximal precursors (position-6) concurrently with differentiation in position-1 (*17*). Position-5 cells do not differentiate but are eliminated by apoptosis, a process shaped in part by the limited range of the signals emitted by xg^O^ (*17*). The lamina therefore provides a rich and tractable model to uncover how distinct cell types – photoreceptors, glia and lamina precursors –deploy and integrate multiple modes of signalling to generate highly precise and reproducible cell fate outcomes.

In this work, we combine targeted genetic manipulations with theoretical modelling to show that glial-induced ERK and Notch signalling dynamically integrate with the Hh morphogen to both shape morphogen gradient formation and enhance positional information, thus specifying lamina cell fates robustly with single-cell resolution.

## Results

### Wrapping glia and xg^O^ induce differential Notch activity in lamina columns

Given the well-established role of Notch signalling in binary cell fate decisions (*18*), we hypothesized that it may subdivide the three Hh-responsive domains to help specify all six lamina cell fates. To investigate this, we assessed Notch activity using three established transcriptional targets: Deadpan (Dpn), *Hairy/E(spl)-related with YRPW motif (Hey)* and *Enhancer of split-mγ (E(spl)-mγ)* (*19–21*).

Dpn is absent from positions-1 and -6, but is expressed transiently in position-2 through -4 at the onset of neuronal differentiation, with peak expression in position-4 cells and no expression in position-5 cells (Fig. 1B). We used *in situ* hybridisation chain reaction (HCR) to examine expression of *Hey*, a neuronal Notch target (*20*). *Hey* is expressed transiently upon neuronal differentiation in position-4 only, consistent with a previous report (Fig. 1C) (*22*). Using a GFP enhancer trap (*E(spl)-mγ-GFP;* (*23*)), we detected *E(spl)-mγ* expression in a subset of lamina precursors just prior to neuronal differentiation and in early neurons (Fig. 1D). Notably, *E(spl)-mγ* is absent from cells in positions-1 and -6, and from older neurons, but expressed in position-2 through -5 (Fig. 1D).

Since Notch targets are known to vary in their activation thresholds and expression dynamics (*24*), these distinct expression patterns likely reflect differences in Notch activity across positions (Fig. 1E). Taken together, they suggest that cells in positions-2 to -5 experience graded Notch activity, with position-4 cells exhibiting the highest and/or most sustained signalling. High and more sustained Notch activity in position-4 is consistent with expression of the Notch ligand *Delta (Dl)* being maintained in position-3, unlike other cell positions where it is transient (Fig. S1A) (*22*). By contrast, cells in positions-1 and -6 showed no evidence of Notch activity.

We noted that Notch targets were expressed just prior to or at the onset of neuronal differentiation, suggesting that Notch activity is linked to glial-induced ERK activity, which drives lamina neuron differentiation (*16, 17*). In other contexts, ERK signalling induces *Dl* expression via the transcriptional effector Pointed (Pnt) (*25, 26*), thus providing a plausible mechanism by which Notch activity may be coupled to glial-dependent ERK signalling in lamina columns. To test whether this mechanism operates in the lamina, we blocked EGFR activity in all glia using a dominant-negative form of the receptor *(Repo^ts^>EGFR^DN^)*, which non-autonomously prevents neuronal differentiation in the lamina (*17*). This led to a marked decrease in *Dl* levels relative to controls, as assessed by HCR (Figs. 1F-H). To confirm that this decrease was indeed due to autonomous ERK pathway activity in the lamina, we used a lamina-specific Gal4 driver (R27G05-Gal4; (*27*)) to knock down Pnt expression during lamina development. This manipulation also led to decreased *Dl* levels in the lamina relative to controls (Figs. S1B,C and 1I). These results indicate that *Dl* expression in the lamina depends on glial-induced ERK activity and Pnt-mediated transcription downstream of ERK signalling, linking glial morphogenesis and ERK activation to the onset of differential Notch activity.

### Notch activity drives binary cell fate decisions within high and low Hh-responsive domains

In both the high and low Hh-responsive domains, Notch activity appears to be present in one cell and absent in its neighbour: position-2 expresses both *E(spl)-mγ* and *Dpn* while position-1 does not; likewise, position-5 expresses *E(spl)-mγ* while position-6 does not (Fig. 1E). Thus, we hypothesized that in these Hh-responsive domains Notch refines cell fate decisions to specify two cell fates in each domain. In wild-type columns, each neuron type can be distinguished based on the combinatorial expression of well-characterised markers (Fig. 1A) (*28, 29*). Sloppy paired 2 (Slp2), Seven-up (Svp), and Brain-specific homeobox (Bsh), in combination, distinguish among L2/L3 (Slp2⁺ only), L1 (Slp2⁺, Svp⁺), L4 (Bsh⁺ only), and L5 (Slp2⁺, Bsh⁺) neurons (Figs. 1J and S1D). L2 and L3 identities are distinguished with *reduced ocelli (rdo)* and *earmuff (erm)* expression: L2 neurons express *rdo* without *erm*, while L3 neurons are defined by *erm* expression regardless of *rdo* expression (Fig. 1K). For brevity, from here onwards we refer to final neuronal identities (*e.g.,* ‘L1’, ‘L2’, etc.) rather than to marker expression profiles. Full marker combinations used to define each neuron type are summarised in Fig. 1A,J-K and detailed in Materials and Methods in the Supplementary Materials.

To test if Notch activity governs the L2 versus L3 fate decision in the high Hh domain, we generated clones mutant for *patched (ptc)*, which elevates Hh signalling cell-autonomously (*30*) using mosaic analysis with a repressible cell marker (MARCM; (*31*)). In *ptc* clones, cells predominantly adopted either L2 or L3 fates, consistent with previous work (Fig. 1L,M) (*15*). Strikingly, when Notch was knocked down by RNAi (N^RNAi^) in *ptc* mutant clones, cells adopted L2 fate preferentially over L3 fate (Fig. 1N). Conversely, increasing Notch signalling by overexpressing the Notch intracellular domain (N^ICD^) in *ptc* MARCM clones drove cells to adopt L3 fate over L2 fate (Fig. 1O; quantified in P). In all cases, neuron type-specific markers were expressed only within the normal spatio-temporal pattern of neuronal differentiation dependent on ERK activation, suggesting that ERK activity is a prerequisite for cells to adopt specific neuron fates. Thus, within the high Hh-responsive domain, varying Notch activity is sufficient to drive a binary fate decision towards L2 in the absence of Notch and towards L3 in its presence.

We then asked whether Notch plays a similar role in the low Hh-responsive domain. In wild-type columns, position-5 cells express the Notch target *E(spl)mγ* and undergo apoptosis, marked by cleaved Death caspase-1 (Dcp1), while position-6 cells lack Notch activity and differentiate as L5 neurons (Fig. 1D,J,Q). To test whether Notch activity drives apoptosis in this context, we mis-expressed the Notch intracellular domain (N^ICD^) specifically in L5 neurons (position-6) using R64B07-Gal4 (*17*). This led to a dramatic increase in Dcp1-positive cells in position-6, in addition to the normal expression pattern in position-5 (Fig. 1Q,R), and increased the overall number of Dcp1-positive cells in the lamina (Fig. 1S). Thus, in the low Hh-responsive domain, Notch activity drives apoptosis over survival, and although ERK activity enables L5 differentiation, it cannot protect against Notch-induced apoptosis.

### Lamina-wide reduction of Notch signalling skews cells towards L2 fate

Unlike cells in the high and low Hh domains, position-3 and position-4 cells in the intermediate Hh domain both exhibited Notch activity, albeit at different levels (Fig. 1B-E). We hypothesised that these differences may nonetheless instruct the decision between L1 and L4 identities. If true, then we predict that reducing Notch signalling activity throughout the lamina would result in each Hh-responsive domain containing cells of only a single fate, corresponding to the cell type that normally experiences lower Notch activity of the pair. To test this, we decreased Notch signalling throughout the lamina by knocking down either Notch or Dl using RNAi. We expected cells in positions-1 and -2 to both adopt L2 fate, cells in positions-3 and -4 to both adopt L1 fate, cells in position-5 to survive rather than die, and cells in position-6 to survive and differentiate as usual into L5s.

Consistent with our prediction, lamina-wide Notch or Dl knockdown abolished Dcp1 expression, indicating that cells normally destined to die at position-5 now survived (Figs. 2A-C and S2A,B). We then assessed L2 and L3 identities and found an expansion in the proportion of L2 neurons, along with a dramatic reduction in the proportion of L3 neurons, indicating that cells were specified with L2 identity at the expense of L3 identity (Figs. 2D-F and S2C,D). Surprisingly, this L2 expansion extended beyond positions-1 and -2 into more proximal positions (Figs. 2D-F and S2C,D), and indeed, we found a corresponding decrease in the proportion of L1 and L4 neurons, suggesting that cells in positions-3 and -4 also adopted L2 identity (Figs. 2G-M and S2E,F). Notably, L5 neurons were unaffected in position-6 (Figs. 2G,H,L,M and S2E,F).

**Fig. 2:**
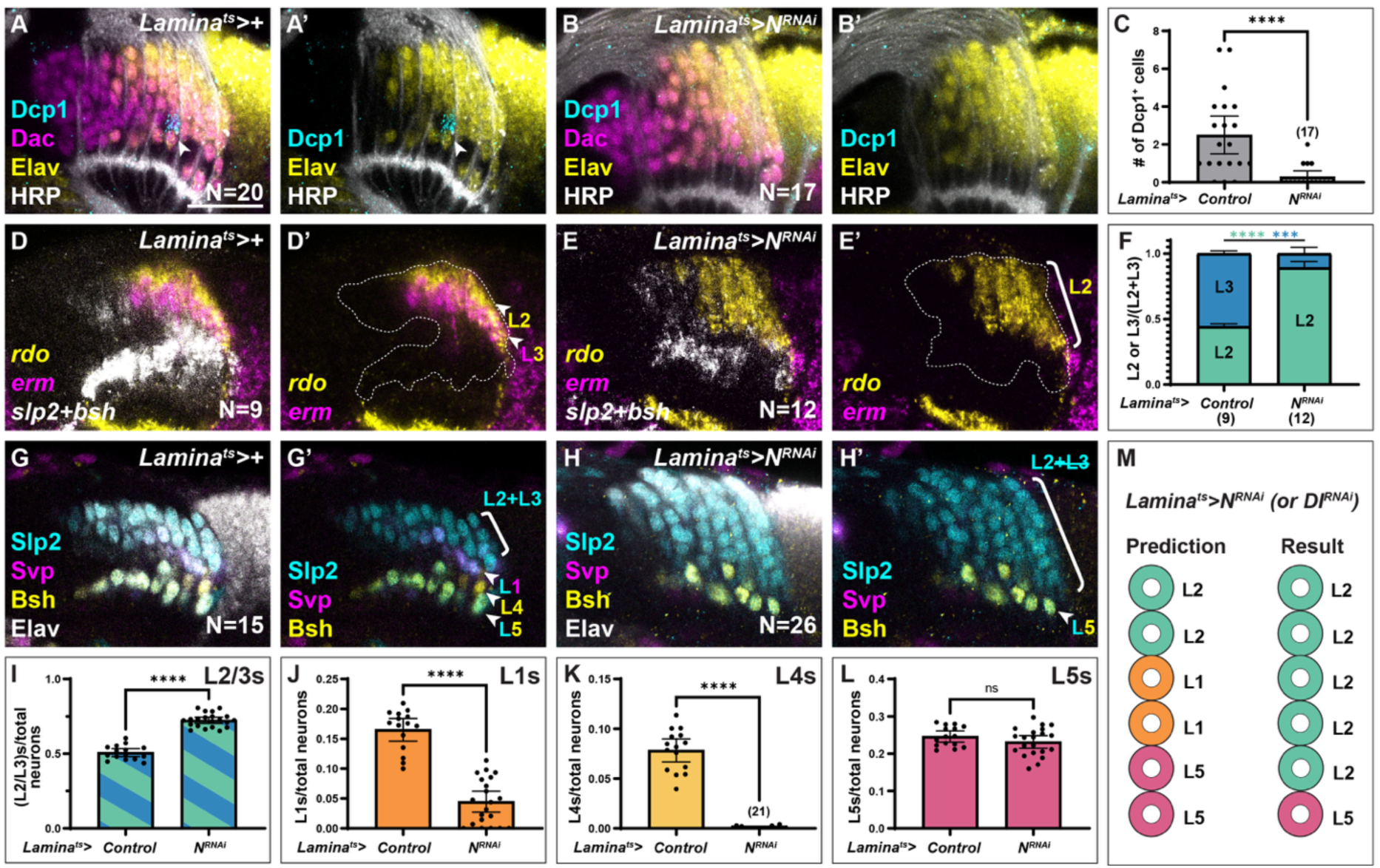
Lamina-wide reduction of Notch signalling skews cells towards L2 fate (A, B) Optic lobes labelled with Dcp1 (cyan), Dac (magenta), Elav (yellow) and HRP (white) in a **(A)** control and with **(B)** lamina-specific Notch knockdown by RNAi. **(C)** Quantifications of the number of cells co-expressing Dcp1 and Dac within lamina columns in a 30µm region of the central lamina from (A) and (B). Error bars represent 95% confidence intervals. Mann-Whitney U-test, p****<0.0001. **(D, E)** Optic lobes labelled with *rdo* (yellow) and *erm* (magenta) to distinguish L2 (*rdo+erm-*) and L3 (*erm+*) neurons and *slp2* combined with *bsh* to mark all lamina neurons (*slp2+bsh*; white) in **(D)** control and when **(E)** Notch was knocked down by RNAi in the lamina. **(F)** Quantifications of the proportion of L2 and L3 neurons out of L2+L3 neurons from (D) and (E) shown together. Mann-Whitney U-test, ***p<0.0005, p****<0.0001. **(G, H)** Optic lobes labelled with Slp2 (cyan), Svp (magenta), Bsh (yellow) and Elav (white) in a **(G)** control and when **(H)** Notch was knocked down by RNAi throughout the lamina. **(I-L)** Quantifications of the proportion of each lamina neuron type (L2/3: Slp2+only; L1: Slp2+Svp+ only; L4: Bsh+ only; L5: Slp2+Bsh+ only) out of total Elav-positive neurons in and (H): **(I)** L2 and/or L3s, **(J)** L1s, **(K)** L4s and **(L)** L5s. Error bars represent 95% confidence intervals. Mann-Whitney U-test, p****<0.0001. **(M)** Schematic of the predicted lamina column patterning outcome for lamina-wide Notch signalling disruptions versus the experimental result. Scale bar = 20 µm.

Taken together, these results demonstrate that reducing Notch signalling globally within the lamina blocks apoptosis at position-5 and leads to a widespread expansion of L2 identity at the expense of L3, L1 and L4 neuron types (Fig. 2M).

### Competence-gated Hedgehog relay contributes to shaping the morphogen gradient

The unexpected expansion of L2 neurons into positions beyond the high Hh-responsive domain (Figs. 2D-L and S2C-F) suggested that the Hh gradient itself and/or its interpretation were altered under lamina-wide Notch signalling reductions. Although photoreceptors are considered the sole source of Hh during the third larval instar and early pupal stages, this assumption has been based on immunoreactivity and enhancer-trap reporter expression (*13, 15*), both of which have limitations. In particular, Hh immunoreactivity was evaluated primarily in the eye disc, with expression in the optic lobe less thoroughly examined (*13*); whereas enhancer traps may not capture all the regulatory elements that drive expression for a given locus. To directly assess *hh* expression, we visualised *hh* transcripts in wild-type optic lobes using HCR (Fig. 3A). As expected, we detected *hh* in photoreceptors, but also observed *hh* expressed transiently in neurons in distal positions of young lamina columns, corresponding to position-1 cells (Fig. 3A,B). To test whether these *hh*-expressing neurons in the lamina differentiate into L2 neurons, which occupy position-1, we used Gal4 inserted in the endogenous *hh* locus (*hh-Gal4*) to drive expression of GFP-tagged Histone 2A variant (His2Av::eGFP). Due to its stability, His2Av::eGFP expression perdures in cells long after Gal4 ceases expression. We found that GFP-positive cells occupied the distal-most positions of lamina columns and expressed *rdo*, indicating that the transient *hh-*expressing cells ultimately differentiate into L2 neurons (Fig. 3C).

**Fig. 3:**
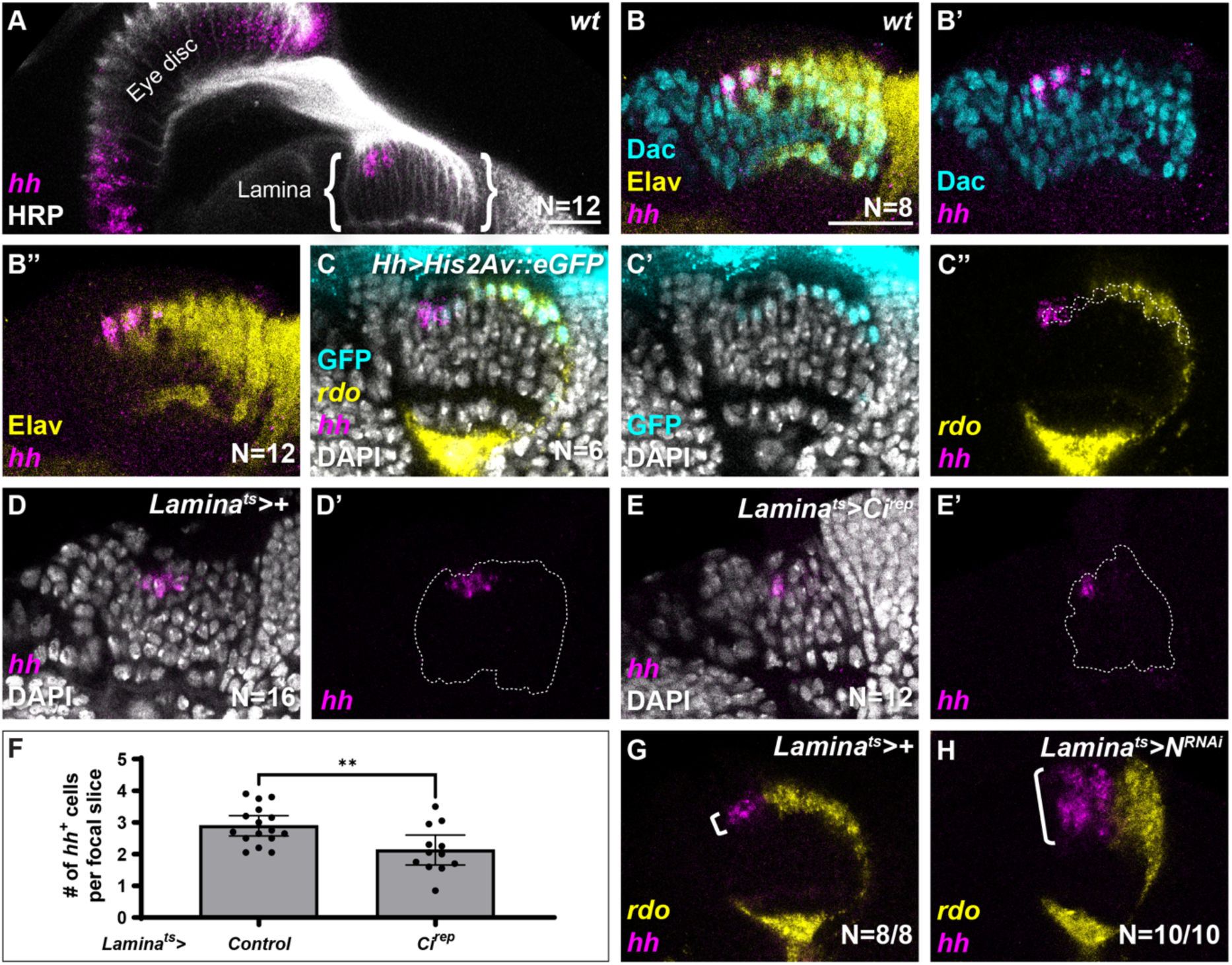
Notch limits propagation of a Hh morphogen relay by restricting *hh* induction in the lamina. **(A)** A wild-type optic lobe showing expression of *hh* (magenta) in photoreceptors in the eye disc and in distal cells of young lamina columns; (HRP; white). **(B)** A wild-type optic lobe showing expression of Dac (cyan), Elav (yellow) and *hh* (magenta). **(C)** hh-Gal4 driving His2Av::eGFP shows GFP (cyan) in the top row (position 1 cells) of the lamina. GFP overlaps with *hh* expression (magenta) and *rdo* (yellow) in older columns, DAPI marks all nuclei (white). The dashed line outlines cells positive for GFP. **(D, E)** Optic lobes labelled with *hh* (magenta) and DAPI (white) in a **(D)** control and when **(E)** Ci^rep^ is misexpressed in the lamina specifically. **(F)** Quantification of the number of *hh-*expressing cells per optical slice within the lamina for **(D)** and **(E)**. Error bars represent 95% confidence intervals. Mann-Whitney U-test, p**<0.01. **(G, H)** Optic lobes labelled with *rdo* (yellow) and *hh* (magenta) in a **(G)** control and when (G) Notch was knocked down by RNAi in the lamina. Scale bars = 20 µm.

What regulates *hh* expression in lamina columns, and what restricts expression to position-1 cells only? Morphogen source relaying is a common developmental strategy deployed to shape gradients, whereby a morphogen induces its own expression in adjacent receiving cells, expanding the effective signalling source (reviewed in (*3*)). We reasoned that the spatial restriction of *hh* expression to position-1 cells in the lamina might reflect a concentration threshold, such that these cells experience the highest concentration of Hh in the lamina derived from the primary photoreceptor source and therefore lie within a Hh concentration responsive domain which induces *hh* expression. To test this, we partially reduced Hh signalling in the lamina by expressing a repressive form of its transcriptional effector, Cubitus interruptus (*lamina>Ci^rep^;* (*15*)) (Fig. 3D,E). Compared to controls, this manipulation decreased the number of *hh* expressing cells in the lamina (Fig. 3D-F), indicating that a minimum threshold of Hh signalling is required for *hh* induction, and that only the most distal lamina cells in position-1 receive sufficient photoreceptor-derived Hh to activate *hh* expression.

An inherent risk of this form of morphogen relay is that the system would be vulnerable to runaway expansion of the *hh* source – as each newly differentiating neuron transiently expresses *hh*, it could increase the local Hh concentration enough to bring the next cell above the threshold required for *hh* expression. To prevent such uncontrolled propagation, additional mechanisms must exist to constrain the competence of lamina cells to respond to Hh with *hh* expression. The transient *hh*-expressing cells ultimately differentiate into L2 neurons (Fig. 3C), a fate governed by high Hh signalling and <u>no</u> Notch signalling (Fig. 1N). We therefore asked whether Notch signalling constrains the competence of a cell to respond to Hh signalling by expressing *hh*. To test this, we examined *hh* expression in the context of lamina-wide Notch knockdowns and observed a dramatic expansion of *hh*-expressing cells within the lamina (Fig. 3G,H).

Together, these findings demonstrate that a relay of *hh* expression between photoreceptors and the lamina expands the effective morphogen source, and that Notch signalling restricts competence for relay of the Hh morphogen in the lamina. This signalling logic could explain the expanded L2 fate observed under lamina-wide Notch or Dl knockdowns: without Notch signalling to restrict relay competence, the system enters a state of runaway source expansion with more proximal cells initiating *hh* expression. This would progressively alter the shape and range of the Hh gradient such that cells in position-2, -3, -4 and -5, normally fated as L3, L1, L4 and apoptotic fate, now experience high Hh signalling and no Notch, and consequently shift their fates towards L2. Under these conditions position-6 cells still differentiate into L5 neurons, which is expected given that cells in position-6 commit to L5 fate in response to xg^O^-mediated ERK activity long before the local expansion of *hh* expression reaches the proximal lamina.

### Precise cell fate patterning requires distinct Notch thresholds

Our data reveal complex interactions between glial-dependent Dl-Notch activity, photoreceptor- and lamina-derived Hh sources and Notch-dependent restriction of the lamina Hh source. To investigate how these interactions impact pattern resolution and robustness, we developed a theoretical framework of ERK-driven Dl-Notch signalling together with Hh morphogen dynamics (see section S1 of Computational Methods in Supplementary Materials). We modelled Dl and Notch signalling dynamics (equations 1 and 2, Fig. 4A) dependent on Dl-driven trans-activation of Notch between neighbouring cells, and inhibition of Dl by Notch signalling (schematic, Fig. S3A)(*32, 33*). To reflect our experimental finding that glia induce *Dl* expression, we simulated temporally staggered Dl production driven by the progressive infiltration of wrapping glia and the spatially fixed xg^O^. We also modelled the secretion, diffusion and degradation dynamics of the Hh morphogen (equation 3, Fig. 4A) along each lamina column. Hh is produced by the photoreceptors, represented as a distal source, and additional local production within the lamina is regulated by Hh and Notch signalling, in line with our experimental findings (Fig. 3A,B).

**Fig. 4:**
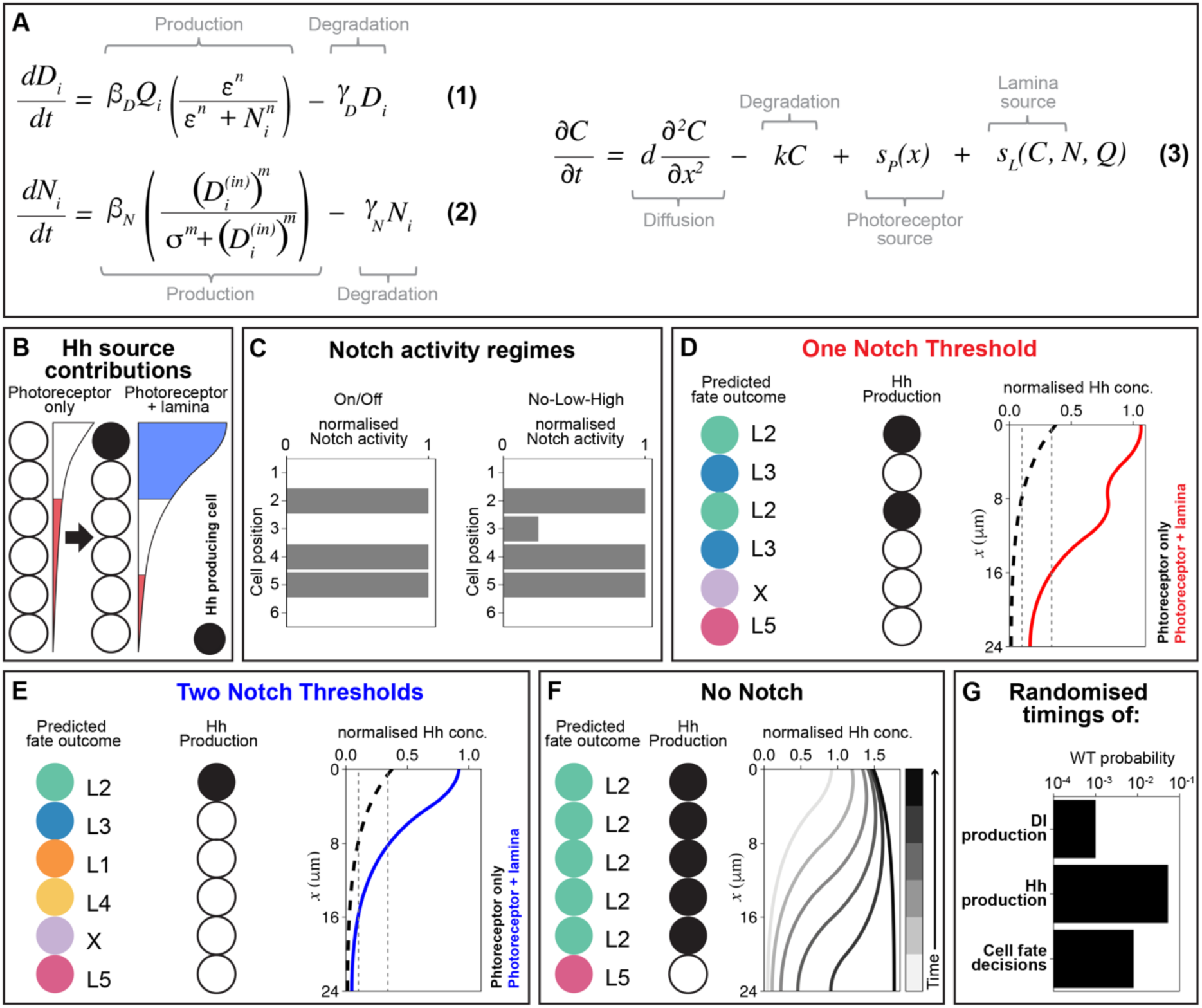
Two Notch thresholds are required for simulation results to recapitulate experimental findings. **(A)** Equations (1) and (2) describe the temporal dynamics of Delta (*D_i_*) concentration and Notch (*N_i_*) activity for the *i*^th^ cell, respectively. The maximal production of *D_i_* and *N_i_* are given by *β_D_* and *β_N_*, respectively, and their degradation rates are represented by *γ_D_* and *γ_N_*, respectively. *Q_i_* = 1 when ERK is activated in the *i*^th^ cell, and 0 otherwise. The constants *ɛ* and *σ* represent transcriptional thresholds for Notch-mediated inhibition of Delta and Delta-mediated activation of Notch, respectively, *n* and *m* are Hill coefficients, and *D*_i_^(in)^ is the total Delta concentration in cells adjacent to the *i*^th^ cell. Equation (3) describes the evolution of the Hh concentration (*C*) in time (*t*) and space (*x*). Hh molecules diffuse with diffusivity *d* and degrade with rate *k*. The source terms *s_P_*(*x*) and *s_L_*(*C*, *N*, *Q*) define Hh production by photoreceptors and lamina cells, respectively (see Computational Modelling Methods in Supplementary Materials for more details). Briefly, Hh is always produced at a constant rate *v_P_* within the photoreceptor source region of width *w_P_* at the edge of the lamina (*s_P_*(*x*) = *v_P_* when −*w_P_* < *x* < 0 and 0 otherwise). If the Notch level at a given position is below the low Notch threshold (*N_i_* < *N*^(*low*^), the average Hh level across the *i*^th^ cell is above the low Hh threshold (〈*C*〉*_i_* > *C*^(*low*)^), and ERK is active (*Q_i_* = 1) then local lamina Hh production is activated in the *i*^th^ cell at rate *v_L_*, and it is zero otherwise. **(B)** A schematic illustrating our assumptions about the relative contribution of the photoreceptor-only Hh gradient and the combined photoreceptor and lamina Hh gradient in a wild-type, and the responsive-domains they establish. **(C)** Notch activity levels in cells normalised to *β*_+_/*γ*_+_ at the time of fate specification fall into two qualitatively distinct regimes depending on parameter values: on/off and no-low-high (see Computational Modelling Methods in Supplementary Materials). In the latter, position-3 cells exhibit low but detectable levels of Notch activity. **(D)** For simulations with one Notch threshold (indicated with a red dashed line in Fig. S3B): schematics of the final cell fates and Hh production state, and the Hh concentration profile at steady state normalised to *v_P_*/*k*, with photoreceptor input alone (black dashed line) and when combined with the lamina sources (red line). **(E)** Same as (D) for two Notch thresholds (indicated with blue dashed lines in Fig. S3C). **(F)** For simulations with no Notch production: schematics of the final cell fates and Hh production states; and the Hh concentration profile normalised to *v_P_*/*k*, which shifts as wrapping glia sequentially activate ERK in cells located more proximally. **(G)** Bar chart representing the probabilities of achieving the wild-type (WT) cell fate pattern following the randomisation of the times of: Delta production, lamina Hh production, or cell fate decisions.

Our experimental data indicate that the three Hh-responsive domains within a lamina column – high, intermediate and low – span two cells each and depend on the combined photoreceptor and lamina sources of Hh. A baseline gradient derived from only photoreceptors, nonetheless, activates *hh* expression only in position-1 and not position-6, despite both experiencing similar Notch and ERK states. This indicates that the photoreceptor-only gradient is sufficient to create at least two concentration-responsive domains. Therefore, in our model we assumed that the photoreceptor-only Hh gradient generates exactly two concentration-responsive domains: intermediate Hh in positions-1 and - 2 and low Hh cells in position-3 through -6 (Fig. 4B; for more details on parameter choices see Computational Modelling Methods in Supplementary Materials). We return to and validate this assumption experimentally below. Accordingly, cells require at least intermediate Hh concentrations to induce *hh* expression. When local Hh production is restricted to position-1 cells, this increases Hh concentration locally and expands the gradient, thereby generating the high Hh domain necessary for the L2 and L3 fates, as well as intermediate, and low Hh-responsive domains across the column (Fig. 4B).

Finally, in line with our experimental observations, a cell is competent to commit to a neuronal fate only when ERK signalling is active in that cell. To capture glial-induced activation of ERK signalling in lamina columns, we represented ERK activity in cells by a binary on/off state that is switched on by the arrival of glia. We assume that the xg^O^, which are positioned below the lamina, only activate ERK signalling in the most proximal cells (position-6), and that this occurs at the same time as ERK activation in the most distal precursor (position-1).

When simulating Dl-Notch dynamics, we observed two qualitatively distinct Notch activity patterns at the time of cell specification, depending on parameter values. In one regime, Notch activity resembled a simple on/off pattern: position-1, -3, and -6 cells all exhibited similarly very low, effectively zero, activity; while position-2, -4 and -5 cells experienced high activity (Fig. 4C, left). In the other regime, Notch activity resolved into three distinct states – no, low and high – such that position-1 and position-6 cells had effectively zero (‘no’) activity; position-2, -4 and -5 cells had high activity; and position-3 cells had low but detectable Notch activity (Fig. 4C, right). Compared with parameter sets that generated the on/off-like Notch activity patterns, those that produced three distinct Notch activity states shared two key features (see Computational Modelling Methods in Supplementary Materials). Firstly, inhibition of Dl production by Notch signalling was relatively weak, permitting low levels of Dl production even in cells with relatively high Notch activity. Secondly, Dl-Notch trans-activation was responsive to small amounts of Dl, enabling position-3 cells to activate Notch signalling to detectable levels despite low Dl input. This regime of no, low, high Notch activity resembles our *in vivo* data showing graded, rather than purely on/off, Notch activity states across each column (Fig. 1B-E), and can be understood by considering the boundary and initial conditions in our system. Cells in positions-1 and -6 experience no Dl input from their neighbours during their specification window, and Dl production in position-1 cells following ERK activation by the wrapping glia generates high Notch activity in neighbouring cells (position-2 and position-5). Due to prior Notch activation, only low levels of Dl can be produced by cells in position-2 upon ERK activation, which generates low but non-zero Notch activity in position-3; this Notch expression is insufficient to repress Dl production in position-3 to the same extent as in position-2, resulting in high Notch levels in the adjacent position-4 cells (see Computational Modelling Methods in Supplementary Materials).

We next considered how cells might interpret Notch activity. One possibility is that graded Notch arises as a mere side-effect of glial-induced *Dl* and column boundary conditions, without bearing any functional significance for cell type specification. Under this assumption, cells would apply a single threshold to read Notch activity, effectively collapsing the low Notch state in position-3 cells and the ‘no’ Notch state in positions-1 and -6 cells into a single “off” state (Fig. S3B). In this case, the three-state regime would be functionally indistinguishable from the on/off-like Notch activity regime. Despite its simplicity, these conditions still allow for a cell fate code in which each of the six cell fates – five neuronal fates and cell death – are defined by a unique combination of Hh-responsive domain (high, intermediate or low) and binary Notch state (See cell fate code in Fig. S3B). Surprisingly, implementing this in our theoretical model failed to generate the wild-type cell fate pattern. Instead, we observed an alternating pattern of L2s and L3s in position-1 through -4 (Fig. 4D). When we examined local *hh* expression and the Hh gradient in these simulations we noted that cells in both position-1 <u>and</u> -3 expressed *hh* (Fig. 4D). This ultimately expands the high Hh domain to include the first 4 cells, thus explaining the patterning outcome (Fig. 4D). These results suggest that the binary Notch and Hh-responsive domain cell fate code (Fig. S3B) is insufficient to achieve the wild-type pattern, in particular, because it fails to restrict lamina Hh production to position-1 cells only.

Therefore, we reasoned that cells may interpret three distinct Notch activity levels at the time of specification – no, low and high – using two thresholds (Fig. S3C), and that spurious Hh production in position-3 could be mitigated if we assume that any minimal Notch activity inhibits Hh production. To test this, we revised the model to include a second Notch threshold and imposed a rule that lamina Hh production is permitted only in cells with <u>no</u> Notch activity (Fig. S3C). With these assumptions, Hh production was restricted to position-1 and the wild-type cell fate pattern was recapitulated (Fig. 4E). These findings suggest that the graded Notch activity we observed *in vivo* (Fig. 1B-D), resulting from column boundary conditions and the temporal dynamics of glial-induced ERK activity and subsequent *Dl* expression, plays a functional role in restricting *hh* expression to position-1 cells. This competence-gating mechanism restricts expansion of the Hh source, likely improving patterning precision while preventing runaway propagation of the source.

Our theoretical model incorporating two Notch thresholds also recapitulated the experimental outcome of lamina-wide Notch knockdown, predicting expansion of the local Hh source and sequential conversion of all but the most proximal cells, which differentiated into L5s, into L2 neurons (Fig. 4F). Therefore, our theoretical model and *in vivo* data together suggest a framework where cells are responsive to three Hh concentration domains and three Notch activity levels, which together determine the competence of cells to respond to Hh signalling with *hh* production, and to commit to one of six fates (L1-L5 and apoptosis; Fig. 4E,F).

### Glia synchronise local Hh production and cell fate decisions with Notch signalling

Key assumptions in our model include that glia induce Dl production, in line with our experimental findings (Fig. 1H), and that both the Hh-producing cell state and neuronal fates are locked-in only upon glial-dependent ERK activation. We asked whether glial-induced ERK activity is also required for lamina cells to express *hh in vivo*, by disrupting EGFR activity in glia. This led to a complete loss of *hh* expression in the lamina (Fig. S4A,B), demonstrating that in addition to requiring sufficient Hh signalling and the absence of Notch activity, *hh* expression in the lamina is further gated by glial-induced ERK activation.

Next, we sought to assess the importance of the temporal coupling between *Dl* expression, lamina *hh* expression and cell type specification using our theoretical model. To do so, we removed the requirement for glial-driven ERK activity and instead allowed cells to initiate Dl or Hh production, or to commit to a cell fate, at random times, independently of glial position and ERK activity. In these simulations, the wild-type pattern emerged only stochastically and at low frequency (Figs. 4G and S4C; see Computational Modelling Methods in Supplementary Materials). Patterning failures were common in simulations with randomised Dl production because local Hh production frequently began before Notch expression could prevent runaway propagation of the Hh source. In the other simulations, patterning failures commonly arose because cells locked-in their fates before local Hh production enhanced the gradient or before Dl-Notch activity had stabilised following activation in adjacent cells (see Section X of Computational Methods in Supplementary Materials). These results suggest that glia confer a crucial time-keeping mechanism that links the emergence of *hh*-expressing cells with the Dl-Notch signalling context that both limits further local *hh* expression and permits accurate cell fate commitment.

### Lamina-wide Notch hyperactivation reveals source-specific Hh contributions and the requirement for Notch thresholds in resolving cell fates

In our theoretical framework, we assume that the photoreceptor-only Hh gradient establishes <u>two</u> concentration domains: intermediate, across positions-1 and -2, and low, across position-3 to -6. We further assume that differential Notch activity within the intermediate Hh-responsive domain instructs the L1 versus L4 fate decision. To test these assumptions, we used the model to predict how cell fates would be altered upon global Notch hyperactivation, a perturbation we could test *in vivo*. First, we modelled this perturbation computationally by introducing baseline, Dl-independent, Notch signalling. Raising baseline Notch signalling above the low Notch threshold eliminated the local *hh*-producing cell state (Fig. 5A). Under these conditions, cell fate decisions were determined by the photoreceptor-only Hh gradient and glial-induced ERK activity patterns, which were unchanged, as well as Dl-Notch signalling consisting of a global elevation superimposed on the endogenous glial-dependent pattern. As a result, without local *hh* production, cells in positions-1 and -2 experienced only intermediate Hh concentrations, while all other cells experienced low Hh concentrations (Figs. 4B and 5A) and were predicted to die due to the combination of low Hh concentration and intermediate-high Notch activity (see Two Notch Threshold cell state/fate code in Fig. S3C; Fig. 5A).

**Fig. 5:**
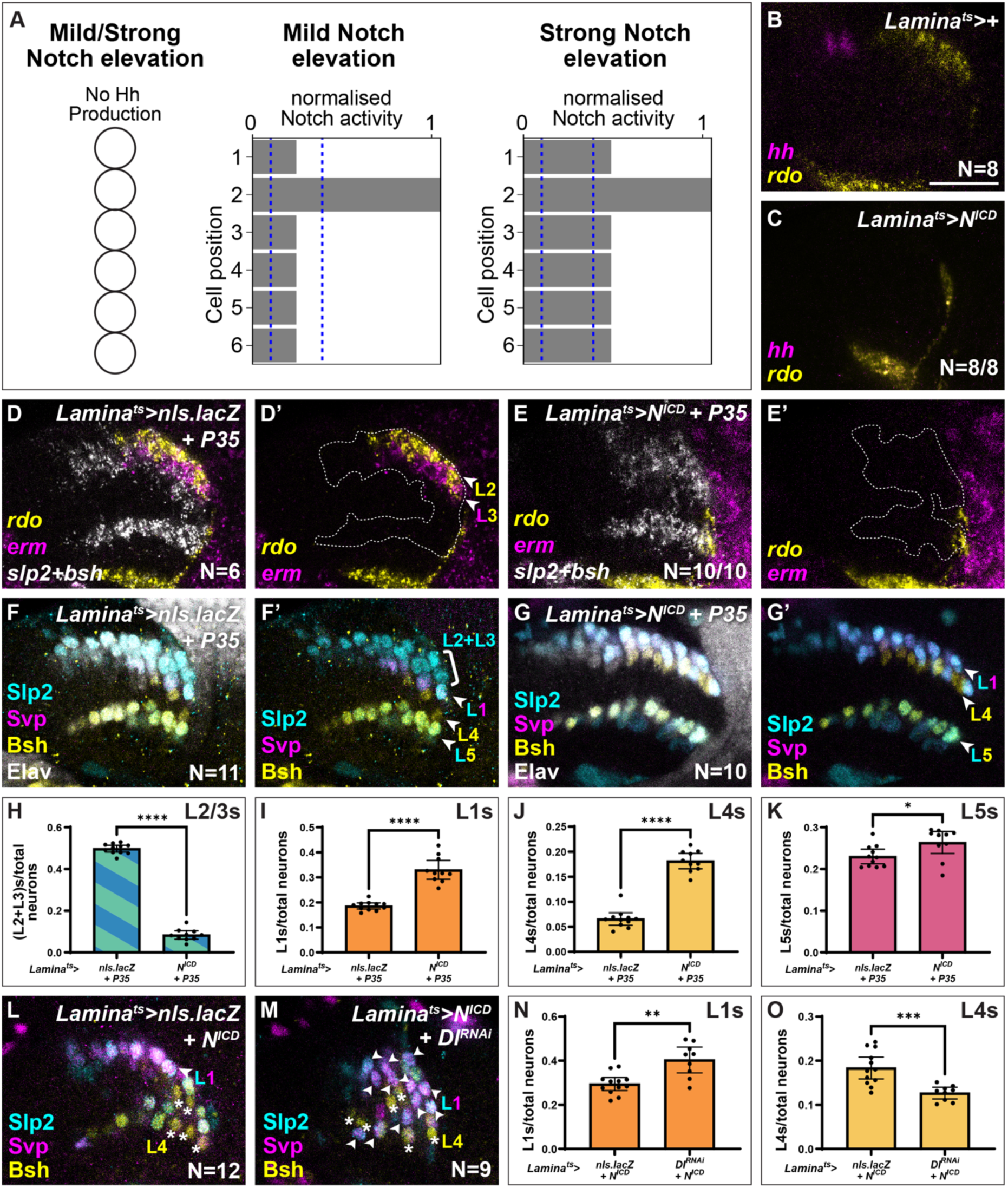
Lamina-wide Notch hyperactivation reveals threshold-dependent cell fate outcomes aligned with model predictions. **(A)** Schematic of the final Hh production states for each cell following mild or strong Notch elevation in simulations and the corresponding Notch signalling levels normalised to *β*_+_/*γ*_+_ in each cell at the time of fate specification. **(B, C)** Optic lobes labelled with *hh* (magenta) and *rdo* (yellow) in a **(B)** control and when **(C)** N^ICD^ was over-expressed in the lamina. **(D, E)** Optic lobes labelled with r*do* (yellow) and *erm* (magenta) to label L2 and L3 neurons, and *slp2* combined with *bsh* (*slp2+bsh*; white) to label all neurons in a **(D)** control and when **(E)** N^ICD^ and P35 were co-expressed in the lamina. **(F, G)** Elav (white) labelling all neurons, together with Slp2 (cyan), Svp (magenta) and Bsh (yellow) to distinguish L2/3s (Slp2+ only), L1s (Slp2+Svp+ only), L4s (Bsh+ only) and L5s (Slp2+Bsh+ only) in a **(F)** control and when **(G)** N^ICD^ and P35 were co-expressed in the lamina. **(H-K)** Quantifications of the proportion of each lamina neuron type out of total Elav-positive neurons per focal slice from (F) and (G): **(H)** L2 and/or L3s, **(I)** L1s, **(J)** L4s and **(K)** L5s. Error bars represent 95% confidence intervals. Mann-Whitney U-test, p*<0.05, p****<0.0001. **(L, M)** Optic lobes labelled with Slp2 (cyan), Svp (magenta) and Bsh (yellow) in a **(L)** control and when **(M)** N^ICD^ is co-expressed with Dl^RNAi^. White arrows indicate L1 neurons (Slp2+Svp+ only), white asterisks indicate L4 neurons (Bsh+ only). **(N, O)** Quantifications of the proportion of **(N)** L1 and **(O)** L4 neurons out of total Elav-positive neurons per optical slice. Error bars represent 95% confidence intervals. Mann-Whitney U-test, p**<0.01, p***<0.0005. Scale bar = 20 µm

The theoretical model further predicted that the specific fates of cells in positions-1 and -2 depend on the strength of baseline Notch signalling elevation. For mild increases in baseline Notch signalling, generating Notch levels above the low and below the high Notch thresholds, the endogenous Dl-Notch signalling maintained differential activity across these cells, specifying L1 and L4 fates in position-1 and position-2, respectively (Fig. 5A). In contrast, raising baseline Notch signalling such that Notch levels are consistently above the high Notch threshold overwhelmed endogenous Dl-Notch signalling, leading to uniform specification of L4 fate in both positions (Fig. 5A). We refer to these conditions as “mild” and “strong” baseline Notch elevation, respectively.

To test these predictions *in vivo*, we mis-expressed N^ICD^ in the lamina. Consistent with model predictions and the role of Notch in restricting the relay of Hh morphogen, overexpressing a single copy of N^ICD^ resulted in the loss of local *hh* expression within the lamina, indicating that the *hh*-producing cell state was not specified (Fig. 5B,C). We also observed a reduction in lamina size, consistent with increased apoptosis predicted by the model (Figs. S5A,B and 5A). To test this directly, we co-expressed the baculoviral caspase inhibitor P35 to block apoptosis alongside N^ICD^ overexpression (Fig. S5C,D). This restored normal lamina size and structure (Fig. S5C-E), but importantly, did not rescue *hh* expression in the lamina (Fig. S5F,G).

We next examined neuron type specification. Although we could not precisely determine whether our manipulations corresponded to the mild or the strong baseline Notch elevation in the model, we reasoned that both scenarios were experimentally distinguishable based on cell fate outcome. We assessed L2 and L3 fate specification and found that neither neuron type was present in laminas mis-expressing N^ICD^ alone or in combination with P35 (Figs. S5H,I and 5D-G), consistent with model predictions that L2 and L3 identities would not be specified (Fig. 5A). We then evaluated L1, L4, and L5 identities and found a small number of L5 neurons, alongside a mixture of predominantly L1 and L4 neurons, in overall smaller laminas with N^ICD^ misexpression alone (Fig. S5J,K). Blocking apoptosis with P35 restored normal lamina morphology and L5 numbers (Figs. 5F-K and S5J,K), confirming that cells experiencing low Hh and some Notch activity are fated for apoptosis. Moreover, we observed that cells in position-1 and -2 adopted L1 and L4 identities, respectively. These outcomes align with the theoretical prediction for mild baseline Notch elevation, where endogenous Dl-Notch feedback remains strong enough to preserve differential Notch signalling between position-1 and position-2 cells. Since intermediate Hh domain fates – L1 and L4 – were specified in positions 1 and 2, with the low Hh domain apoptotic fate in all other positions, these results validate our assumption that the photoreceptor-only Hh gradient sets up two Hh-responsive domains: intermediate Hh for positions-1 and -2, and low Hh for positions-3 to -6.

As a further test of our hypothesis and the theoretical assumption that differential Notch signalling between position-3 and position-4 instructs the L1 versus L4 fate decision, we reasoned that knocking down Dl expression in the N^ICD^ overexpression background should “break” the endogenous Dl-Notch signalling, causing both positions-1 and -2 to adopt the same fate. Our model predicted that this fate would be L1, as these cells would maintain low Notch activity but be deprived of the Dl-Notch trans-activation necessary for the cell in position-2 to activate the high levels of Notch signalling required for L4 specification (Fig. S5L). Consistent with these predictions, co-expressing N^ICD^ and Dl^RNAi^ in the lamina led to an increase in the proportion of L1 neurons at the expense of L4 neurons compared to N^ICD^-only controls (Figs. 5L-O and S5M,N).

Together, these results demonstrate that: (i) the photoreceptor-only Hh source establishes two Hh-responsive domains in lamina columns; and (ii) differential Notch activity in the intermediate Hh-responsive domain directs the decision between L1 and L4 neuronal fates. Thus, by combining experiments and theory, we identify the conditions that specify six cell fates within lamina columns.

## Discussion

Our work demonstrates that morphogen, juxtacrine, and non-morphogen secreted signals integrate dynamically in space and time to sculpt the morphogen gradient and couple cell fate decisions to appropriate competence windows, thereby yielding reproducible high-resolution patterns of cell fate.

The importance of visual processing for animal fitness places strong selective pressure on the visual system to achieve both precision and robustness in its underlying circuitry, demands that are typically viewed as competing and reflecting an underlying trade-off (*34–36*). In the lamina, we show that xg^O^ and infiltrating wrapping glia establish a deterministic spatiotemporal pattern of graded Dl-Notch activity, which restricts cellular competence to respond to and relay Hh, thereby shaping the Hh gradient in space and time. In parallel, glial-induced ERK activity acts as a developmental timer, coordinating the *hh-* expressing cell state and cell fate decisions with differential Notch signalling. Through this integration, Hh signalling, glial-induced ERK activity and graded Notch activity collectively instruct cell fate decisions with single-cell resolution, repeatedly and reproducibly across hundreds of columns.

Developmental patterns of cell identity can be remarkably fine-grained, down to the scale of single cells. For instance, in the vertebrate inner ear and Drosophila sensory organs, juxtacrine Delta-Notch signalling plays key roles in generating such patterns (reviewed in (*37, 38*)). By contrast, the fine-grained pattern of segment polarity genes in the Drosophila embryo, perhaps the best characterised example of such a pattern arising from morphogens, emerges through successive layers of transcriptional regulation that progressively subdivide the broad gene expression domains set by maternal morphogens, ultimately yielding cell identities resolved to single cells (reviewed in (*39, 40*)). Such multi-layered regulation has been proposed to render positional information more accessible to cells and to enhance the robustness of the resulting patterns (*41, 42*). Recent theoretical studies have begun exploring how global positional information from morphogens can combine with self-organising patterning mechanisms, such as reaction-diffusion or juxtacrine signalling, to inform cell fate decisions (*43, 44*). These interactions are proposed to endow developing tissues with diverse patterning strategies differing in timing, precision, robustness, and evolvability (*43, 44*). Indeed, our *in vivo* data in the lamina reveal that single-cell resolution in tissue patterning can emerge without iterative refinement, and can be achieved instead through the dynamic integration of morphogen and juxtacrine signals in space and time. Our findings highlight three general principles:

First, we show that patterning mechanisms such as lateral inhibition, which are often associated with stochastic outputs, can be constrained to act deterministically, thereby providing positional information to augment that provided by morphogens, as anticipated by theory (*45*). In the lamina, column boundary conditions together with temporally unfolding glial-induced *Dl* expression provide constraints that generate a deterministic, graded Notch activity pattern, which subdivides Hh-responsive domains to increase the resolution of patterning.

Second, we identify a morphogen-independent mechanism that spatially restricts morphogen relay, thereby preventing runaway source propagation. Morphogen relay is a known strategy for modulating gradient shape and range over time, but it carries the risk of runaway signal spread (*46–52*). In many contexts, such as for Nodal signalling in the zebrafish blastula and in human gastruloids, and Bone Morphogenetic Protein signalling in the mouse dorsal neural tube, relay is self-limiting, *i.e.,* morphogen signalling induces expression of a repressor that inhibits further relaying (*50, 52, 53*). Alternatively, in the vertebrate ventral neural tube, relaying is curtailed by external signals: Fibroblast Growth Factor from the caudal parts of the embryo and later retinoic acid from adjacent somites limit floor plate identity, thereby restricting Sonic hedgehog relay (*54–56*). Likewise in the lamina, graded Notch activity, independent of morphogen signalling, gates relay competence, helping to localise the Hh source, and shapes the gradient with high spatial and temporal precision.

Third, we find that accurate and robust pattern formation requires temporal coupling between cell state transitions and the dynamics of instructive signalling. In other words, appropriate fate choices depend on restricting competence windows in space and time, enabling the same signalling components to be redeployed to different ends in different contexts. In the lamina, this coordination is achieved through glial-induced ERK activity, which ensures fate commitment only once differential Notch activity is established and the Hh gradient has been shaped by both the photoreceptor and lamina sources.

Our findings provide a mechanistic interpretation of lamina patterning to reconcile previous observations. Prior work proposed that Notch signalling distinguishes L4 from L5 fate, based on shared expression of the transcription factor Bsh and the observation that lamina-wide Notch disruptions led to loss of L4s, which was interpreted as an L4-to-L5 fate conversion (*22*). Conversely, N^ICD^ overexpression was proposed to drive the inverse process (*22*). By examining all lamina cell identities and integrating these data with computational modelling, we demonstrate that these phenotypes are instead due to changes in the Hh gradient. Specifically, L2s expand at the expense of L3, L1 and L4 identities due to runaway Hh source expansion in the lamina in the absence of Notch-mediated restriction, and L5 loss under N^ICD^ overexpression is due to elevated apoptosis, not fate switching.

This study demonstrates that, rather than acting downstream of or parallel to morphogen input, signals such as Dl-Notch and ERK synergise with the Hh morphogen, unfolding alongside it in space and time to encode and enhance positional information. This interplay is orchestrated by glial morphogenesis, whose dynamics act as a timekeeper for the patterning process. Thus, precise and robust patterning outcomes emerge from dynamic interactions among distinct cell types, driving multiple signalling modes across spatial and temporal scales.

## Supporting information

HCR Probe sequences

## Acknowledgements

We are grateful to S. Ackerman, M. Amoyel, J. Briscoe, C. Desplan, J. Delas, N. Konstantinides, R. Mayor, T. Patel, J. Rister, C. Stern, N. Tapon and members of the Fernandes and Amoyel labs for critical feedback on the work and manuscript. AD was funded by UCL’s Research Opportunity Scholarship, VMF is funded by a Wellcome CDA (225986/Z/22/Z), EMBO Young Investigator Award and Lister Institute Prize. ZH and LM were supported by the Francis Crick Institute, which receives its core funding from Cancer Research UK, the UK Medical Research Council, and the Wellcome Trust.

## List of Supplementary Materials

**Fig. S1:**
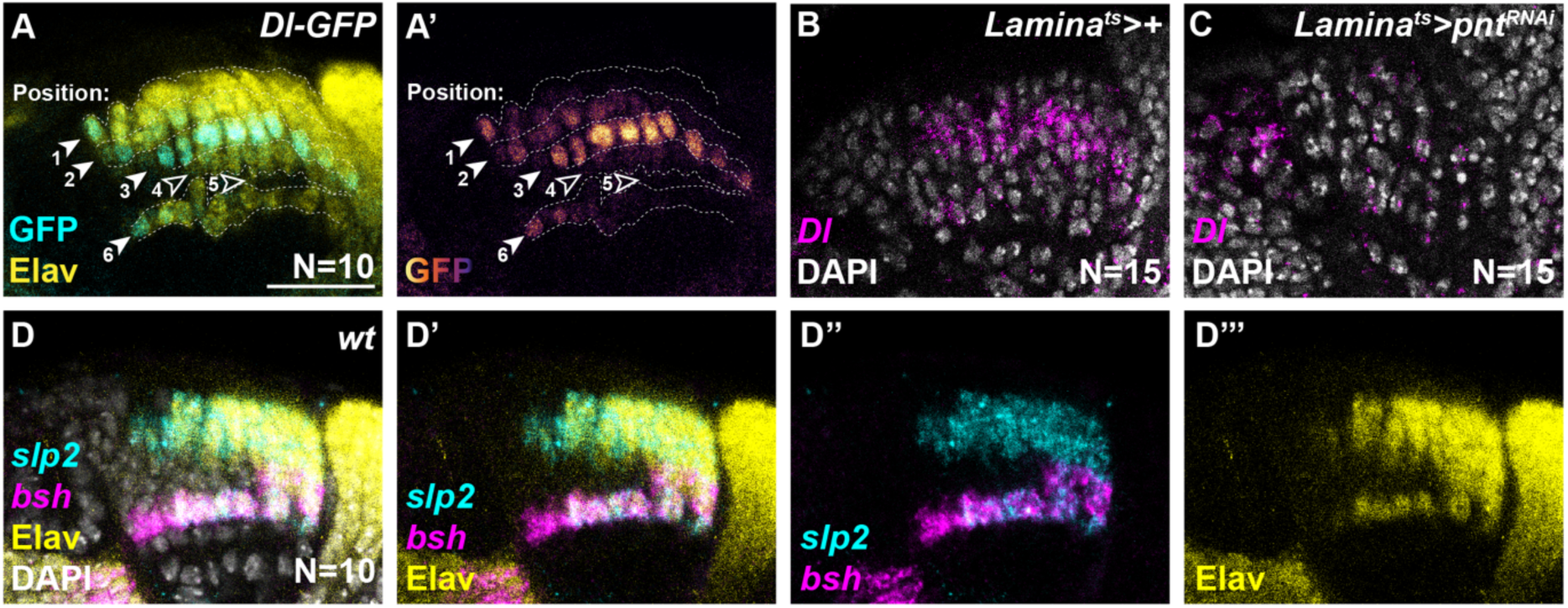
Pointed is required for *Delta* expression in the lamina. **(A)** Dl-GFP expression (GFP; cyan in (A) and pseudo-coloured in (A’)) with neurons marked by Elav (yellow). **(B, C)** *Dl* expression (magenta) with DAPI (white) in **(B)** a control lamina and when **(C)** the ERK transcriptional effector Pointed (Pnt) is knocked down in the lamina. **(D)** Wild-type expression of *sloppy paired 2 (slp2*; cyan) with *brain specific homeobox (bsh*; magenta) and Embryonic lethal abnormal vision (Elav; yellow), showing overlap between the combined domains of *slp2* and *bsh* expression with Elav expression. We use *slp2+bsh* to mark all neurons in subsequent figures. DAPI (white) marks nuclei. Scale bar = 20 µm

**Fig. S2:**
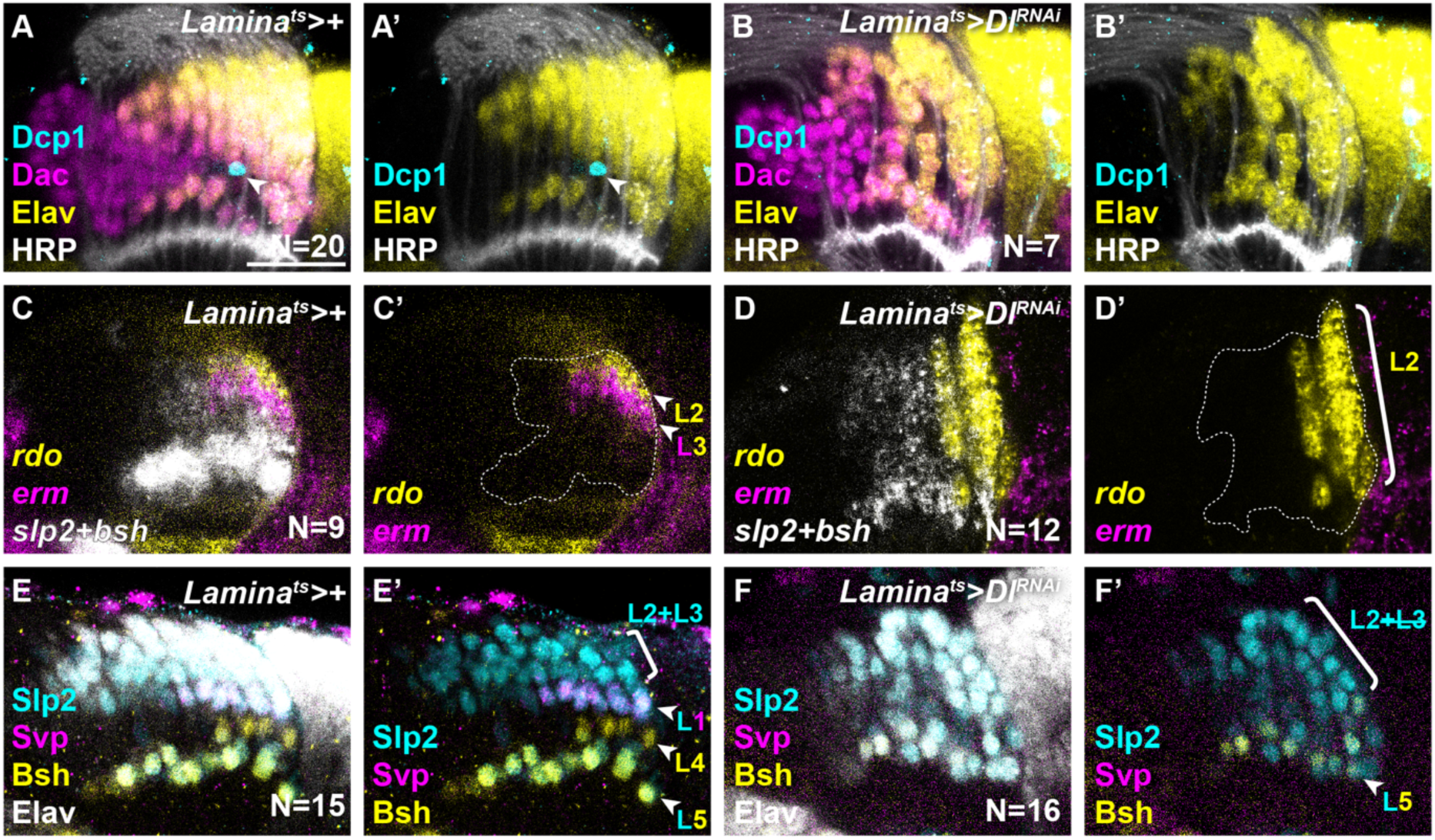
Lamina-wide knockdown of Dl skews cells towards L2 fate. **(A, B)** Optic lobes labelled with Dcp1 (cyan), Dac (magenta), Elav (yellow) and HRP (white) in **(A)** a control and **(B)** when Dl^RNAi^ is expressed throughout the lamina. **(C, D)** Optic lobes labelled with *rdo (*yellow) and *erm (*magenta) to distinguish between L2 (*rdo+erm-*) and L3 (*erm+*) neurons, and *slp2+bsh* (white) to label all neurons in **(C)** a control and **(D)** when Dl^RNAi^ is expressed throughout the lamina. **(E, F)** Elav (white) labelling all neurons, together with Slp2 (cyan), Svp (magenta) and Bsh (yellow) to distinguish L2/3s (Slp2+ only), L1s (Slp2+Svp+ only), L4s (Bsh+ only) and L5s (Slp2+Bsh+ only) in **(E)** a control and **(F)** when Dl^RNAi^ is expressed throughout the lamina. Scale bar = 20 µm

**Fig. S3:**
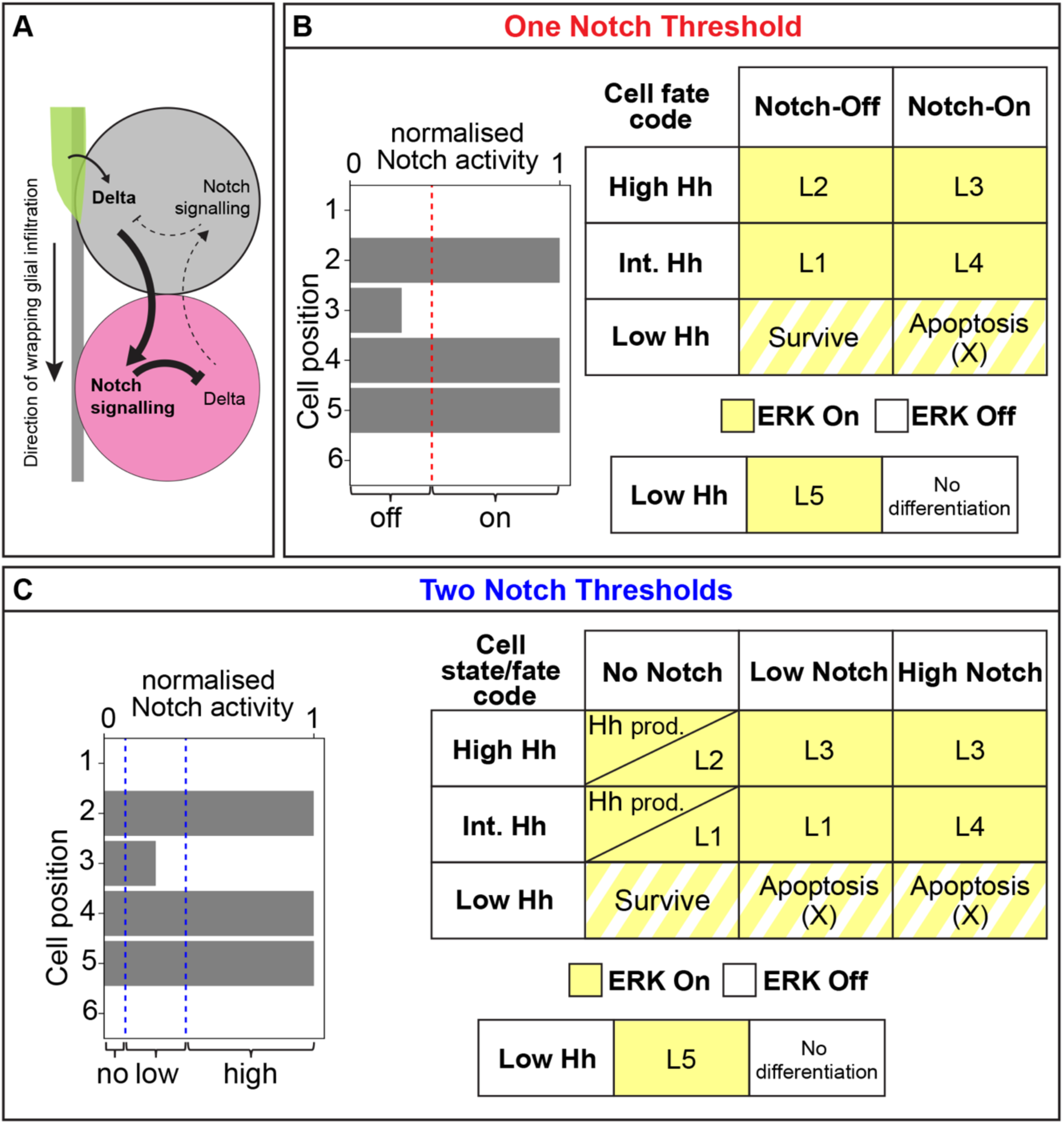
Cell fate/state codes when we implement one or two Notch activity thresholds in our theoretical model. **(A)** Schematic of the Dl-Notch interactions captured in equations (1) and (2) in Fig. 4A. **(B)** Applying one activity threshold (red dashed line) to Notch signalling levels normalised to *β*_+_/*γ*_+_ in cells at the time of specification, collapses the no-low-high Notch activity regime (Fig. 4C) to an on/off-like state. Cell fate code combining these Notch conditions (on/off) with the three Hh responsive domains (low, intermediate and high) uniquely defines all six lamina cell fates (L1-L5 and apoptosis). **(C)** The same Notch signalling levels normalised to *β*_+_/*γ*_+_ at the time of cell specification as in (A) but with two activity thresholds applied (blue dashed lines). Cell state/fate code combining these Notch conditions (no, low and high) with the three Hh responsive domains (low, intermediate and high) uniquely defines the Hh producing cell state as well as all six lamina cell fates (L1-L5 and death).

**Fig. S4:**
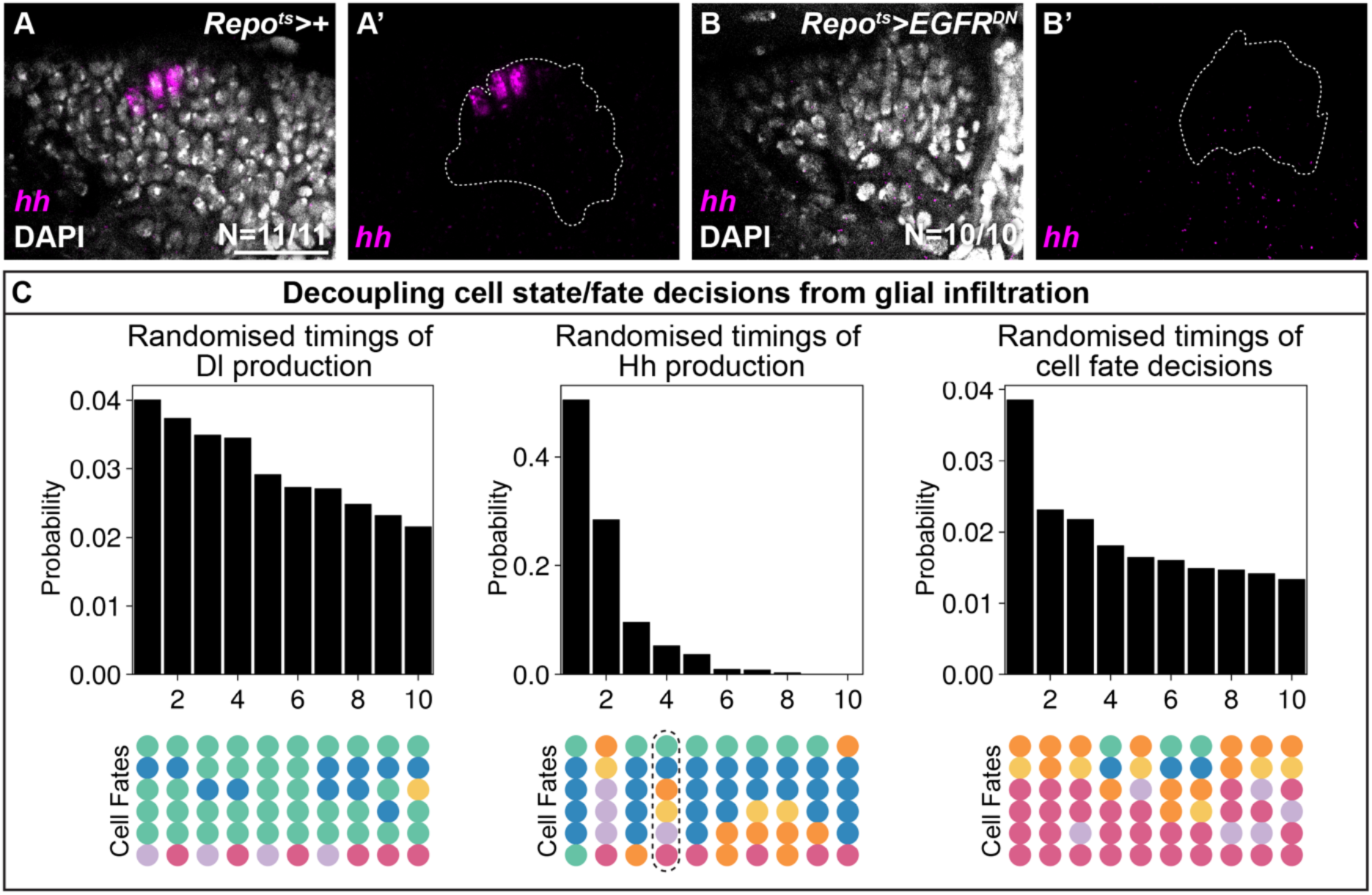
Timekeeping by glia through ERK activity is required for appropriate lamina patterning. **(A, B)** Optic lobes labelled with *hh* (magenta) and DAPI (white) in **(A)** a control and **(B)** when EGFR^DN^ is expressed in all glia. Scale bar = 20 µm. **(C)** Bar charts representing the probabilities of achieving specified cell fates following the randomisation of the times of: Delta production, lamina Hh production, or cell type specification. Only the 10 most abundant cell fates are shown. A black dashed line outlines the wildtype pattern.

**Fig. S5:**
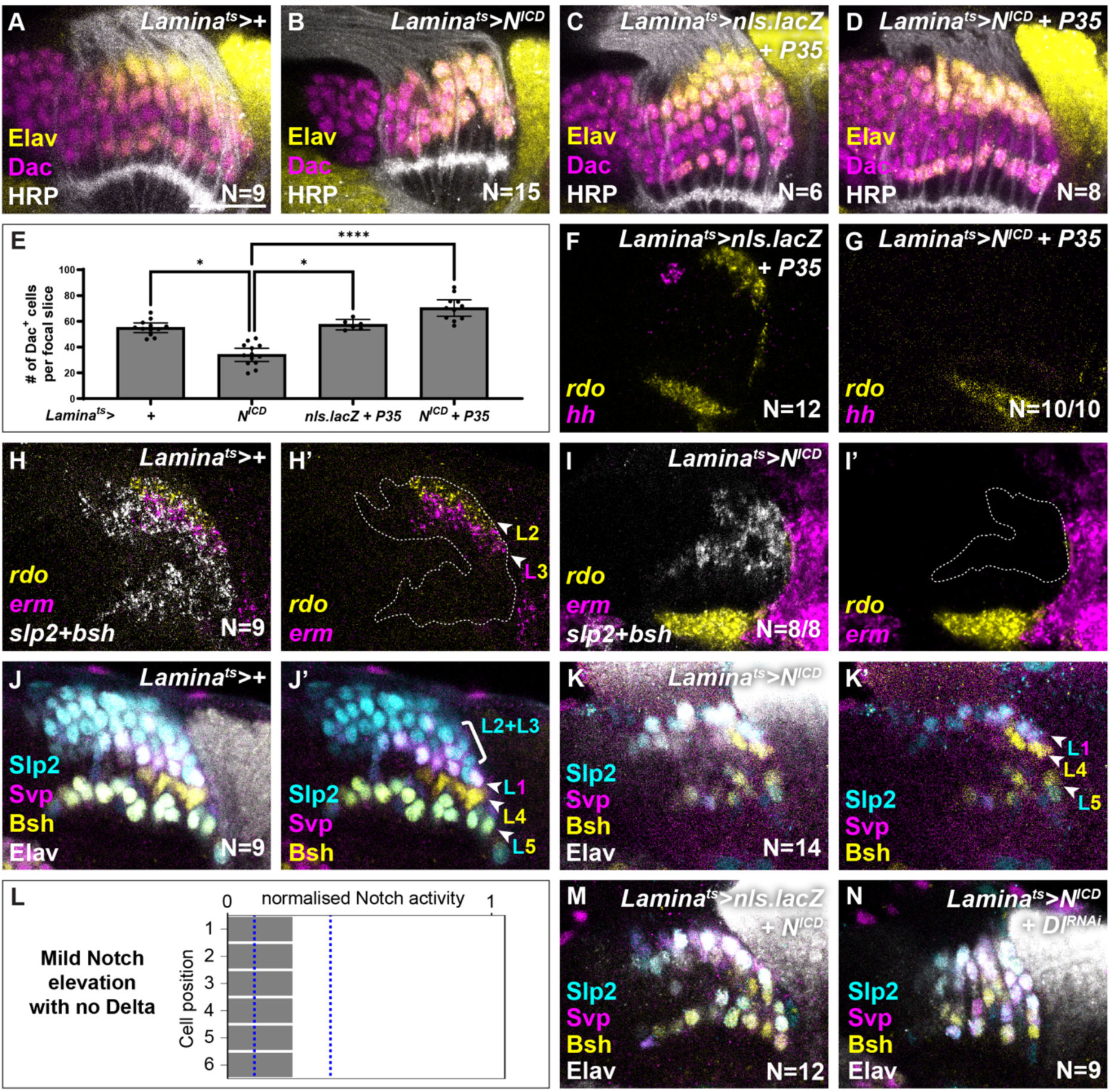
Lamina-wide N^ICD^ overexpression reduces lamina size and skews surviving neurons towards L1 and L4 fates. **(A-D)** Dac (magenta), Elav (yellow) and HRP (white) in **(A)** a control, **(B)** when N^ICD^ is overexpressed in the lamina, **(C)** when nls.lacZ and P35 are expressed in the lamina (titration control) and **(D)** when N^ICD^ and P35 are expressed in the lamina. **(E)** Quantification of the number of Dac+ cells per focal slice for (A-D). One-way ANOVA with Dunn’s multiple comparisons test, p*<0.05, p****<0.0001. **(F, G)** *hh* (magenta) and *rdo* (yellow) expression in optic lobes when **(F)** nls.lacZ and P35, and **(G)** N^ICD^ and P35 are expressed throughout the lamina. *hh* expression was absent in **(G)** in 10/10 lobes. **(H, I)** Optic lobes labelled for *rdo* (yellow) and *erm* (magenta) to mark L2 and L3 neurons, and *slp2+bsh* (white) to mark all neurons in **(H)** control and **(I)** lamina-wide N^ICD^ over-expression. *rdo* and *erm* were absent in **(I)** in 8/8 lobes. **(J, K)** Elav (white) labelling all neurons, together with Slp2 (cyan), Svp (magenta) and Bsh (yellow) to distinguish L2/3s (Slp2+ only), L1s (Slp2+Svp+ only), L4s (Bsh+ only) and L5s (Slp2+Bsh+ only) in **(J)** control and **(K)** lamina-wide N^ICD^ over-expression. **(L)** Notch signalling levels normalised to *β*_+_/*γ*_+_ in each cell at the time of fate specification from simulations of mild Notch elevation combined with no Delta, along with a schematic of the final cell fates. **(M, N)** Optic lobes labelled with Slp2 (cyan), Svp (magenta), Bsh (yellow) and Elav (white) in a **(M)** control and when **(N)** N^ICD^ is co-expressed with Dl^RNAi^. Scale bar = 20 µm

