## Supplementary Material for "Morphogen and juxtacrine signalling dynamically integrate to specify cell fates with single-cell resolution"

by

Alicia Donoghue[1§], Lewis S Mosby[2,3,4§], Inês Lago-Baldaia[1], Tamara Hodgetts[1, 2], Evelina Ursu[1], Zeynep Erten[1], Zena Hadjivasiliou[2,3,4]* and Vilaiwan M Fernandes[1]*

[1] Department of Cell and Developmental Biology, University College London, University College London, Gower Street, London, WC1E 6DE.
[2] Mathematical and Physical Biology Laboratory, The Francis Crick Institute, 1 Midland Road, London NW1 1AT, UK.
[3] Department of Physics and Astronomy, University College London, Gower Street, London WC1E 6BT, UK.
[4] London Centre for Nanotechnology, 19 Gordon Street, London WC1H 0AH, UK.

[§]These authors contributed equally to this work

**This document includes:**

- Figs. S1 to S5
- Materials and Methods
- References
- Tables S1
- Computational Modelling Methods

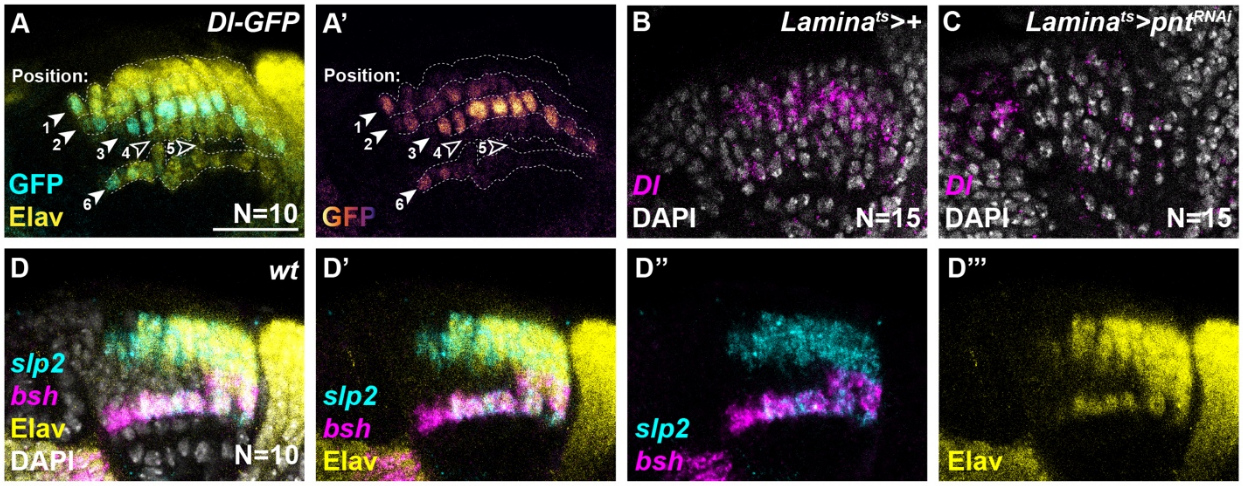

**Fig. S1: Pointed is required for *Delta* expression in the lamina**

**(A)** Dl-GFP expression (GFP; cyan in (A) and pseudo-coloured in (A')) with neurons marked by Elav (yellow).

**(B, C)** *Dl* expression (magenta) with DAPI (white) in **(B)** a control lamina and when **(C)** the ERK transcriptional effector Pointed (Pnt) is knocked down in the lamina.

**(D)** Wild-type expression of *sloppy paired 2 (slp2*; cyan*)* with *brain specific homeobox (bsh*; magenta*)* and Embryonic lethal abnormal vision (Elav; yellow), showing overlap between the combined domains of *slp2* and *bsh* expression with Elav expression. We use *slp2+bsh* to mark all neurons in subsequent figures. DAPI (white) marks nuclei.

Scale bar = 20 µm

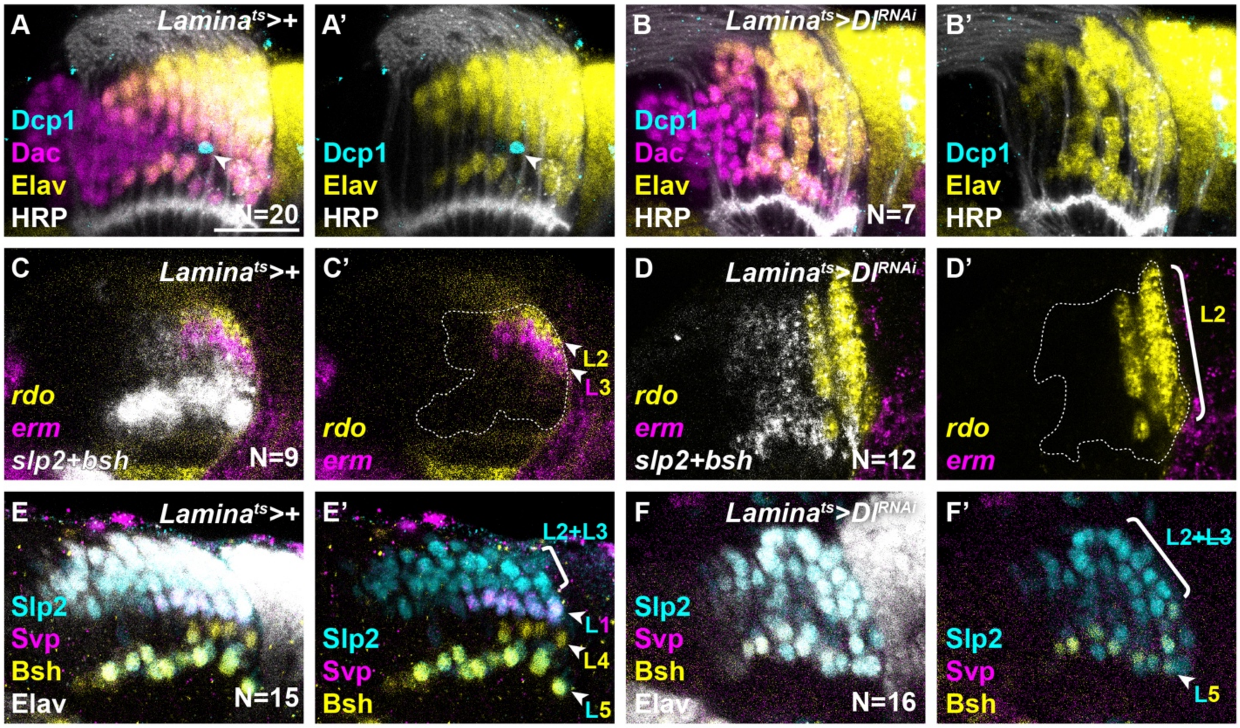

**Fig. S2: Lamina-wide knockdown of Dl skews cells towards L2 fate**

**(A, B)** Optic lobes labelled with Dcp1 (cyan), Dac (magenta), Elav (yellow) and HRP (white) in **(A)** a control and **(B)** when Dl$^{RNAi}$ is expressed throughout the lamina.

**(C, D)** Optic lobes labelled with *rdo (*yellow*)* and *erm (*magenta*)* to distinguish between L2 (*rdo*+*erm*-) and L3 (*erm*+) neurons, and *slp2+bsh* (white) to label all neurons in **(C)** a control and **(D)** when Dl$^{RNAi}$ is expressed throughout the lamina.

**(E, F)** Elav (white) labelling all neurons, together with Slp2 (cyan), Svp (magenta) and Bsh (yellow) to distinguish L2/3s (Slp2+ only), L1s (Slp2+Svp+ only), L4s (Bsh+ only) and L5s (Slp2+Bsh+ only) in **(E)** a control and **(F)** when Dl$^{RNAi}$ is expressed throughout the lamina.

Scale bar = 20 µm

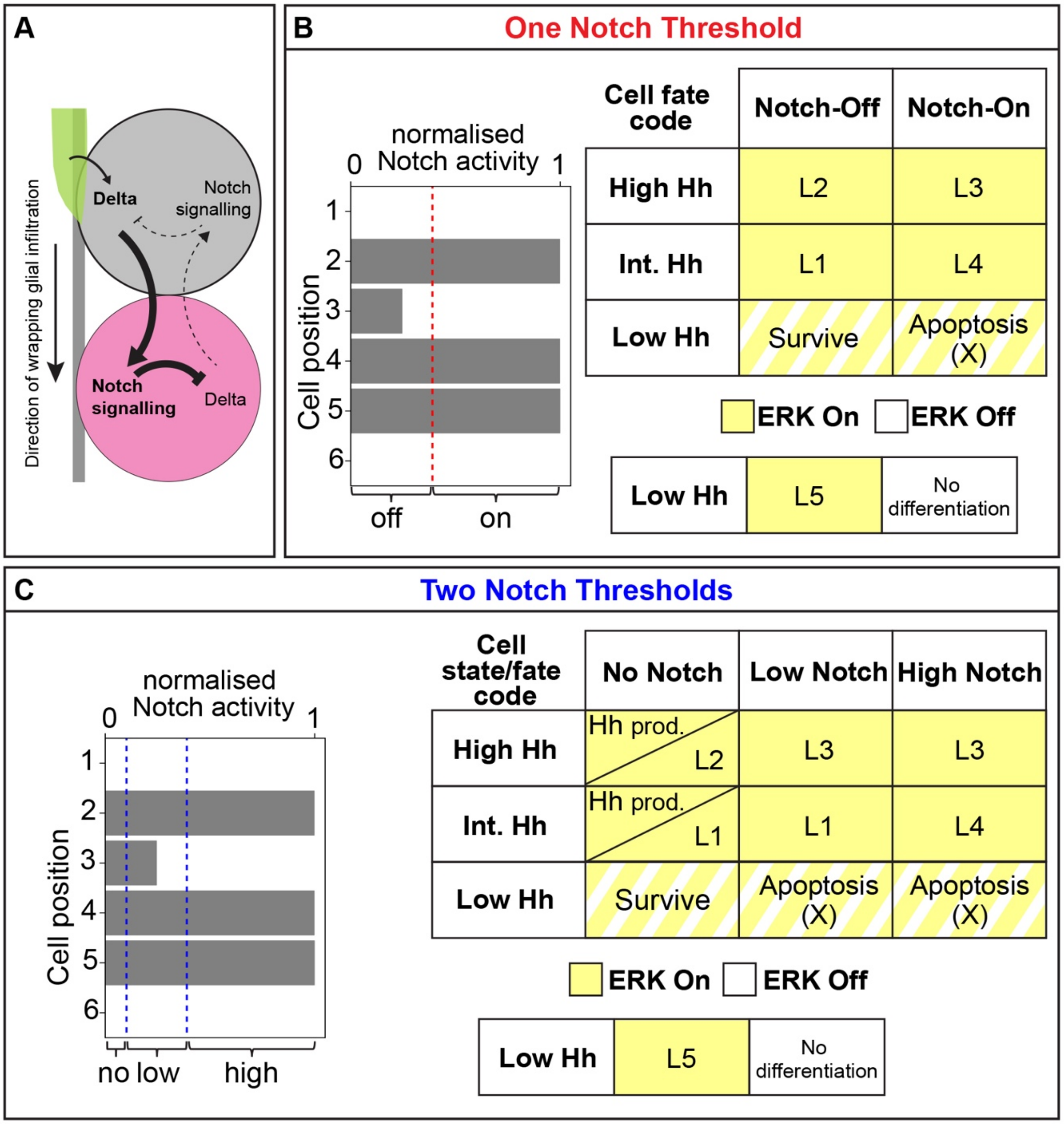

**Fig. S3: Cell fate/state codes when we implement one or two Notch activity thresholds in our theoretical model**

**(A)** Schematic of the Dl-Notch interactions captured in equations (1) and (2) in Fig. 4A.

**(B)** Applying one activity threshold (red dashed line) to Notch signalling levels normalised to $\beta_N/\gamma_N$ in cells at the time of specification, collapses the no-low-high Notch activity regime (Fig. 4C) to an on/off-like state. Cell fate code combining these Notch conditions (on/off) with the three Hh responsive domains (low, intermediate and high) uniquely defines all six lamina cell fates (L1-L5 and apoptosis).

**(C)** The same Notch signalling levels normalised to $\beta_N/\gamma_N$ at the time of cell specification as in (A) but with two activity thresholds applied (blue dashed lines). Cell state/fate code combining these Notch conditions (no, low and high) with the three Hh responsive domains (low, intermediate and high) uniquely defines the Hh producing cell state as well as all six lamina cell fates (L1-L5 and death).

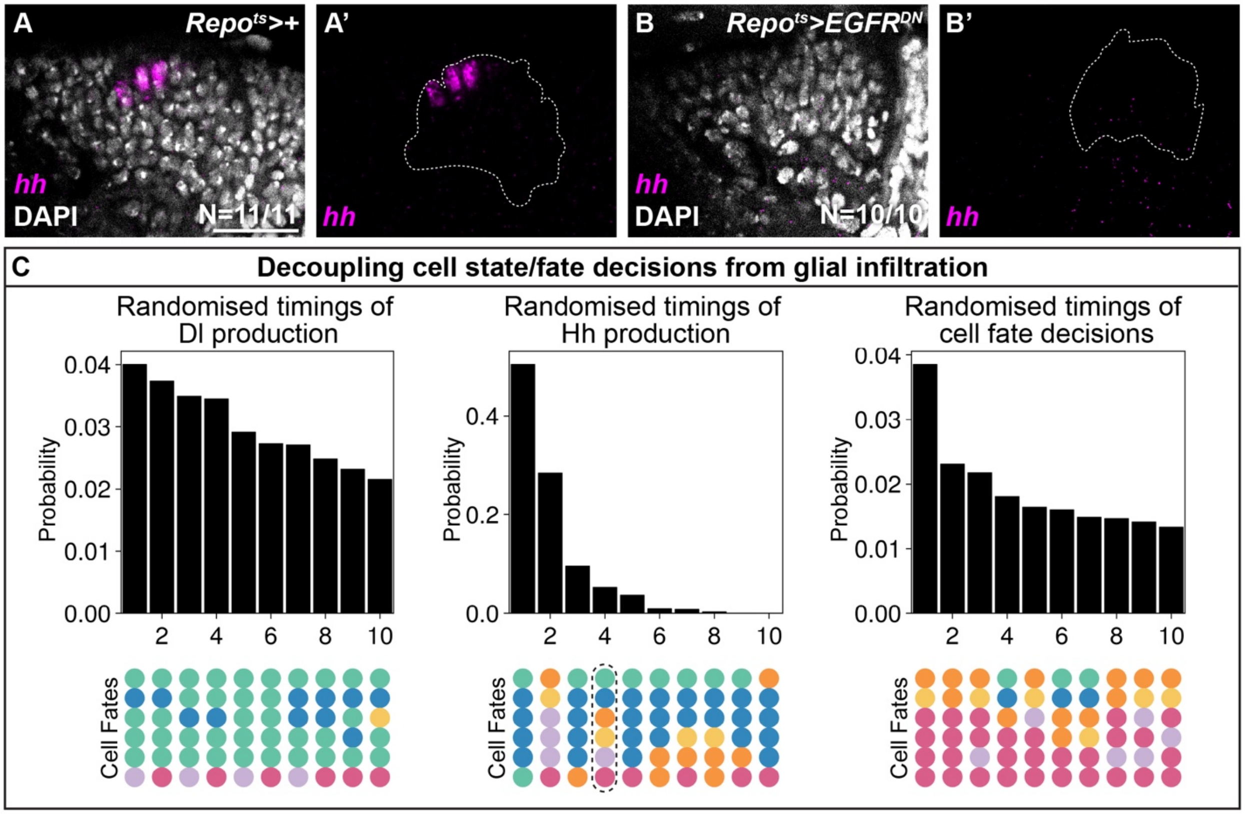

**Fig. S4: Timekeeping by glia through ERK activity is required for appropriate lamina patterning**

**(A, B)** Optic lobes labelled with *hh* (magenta) and DAPI (white) in **(A)** a control and **(B)** when $EGFR^{DN}$ is expressed in all glia. Scale bar = 20 µm.

**(C)** Bar charts representing the probabilities of achieving specified cell fates following the randomisation of the times of: Delta production, lamina Hh production, or cell type specification. Only the 10 most abundant cell fates are shown. A black dashed line outlines the wildtype pattern.

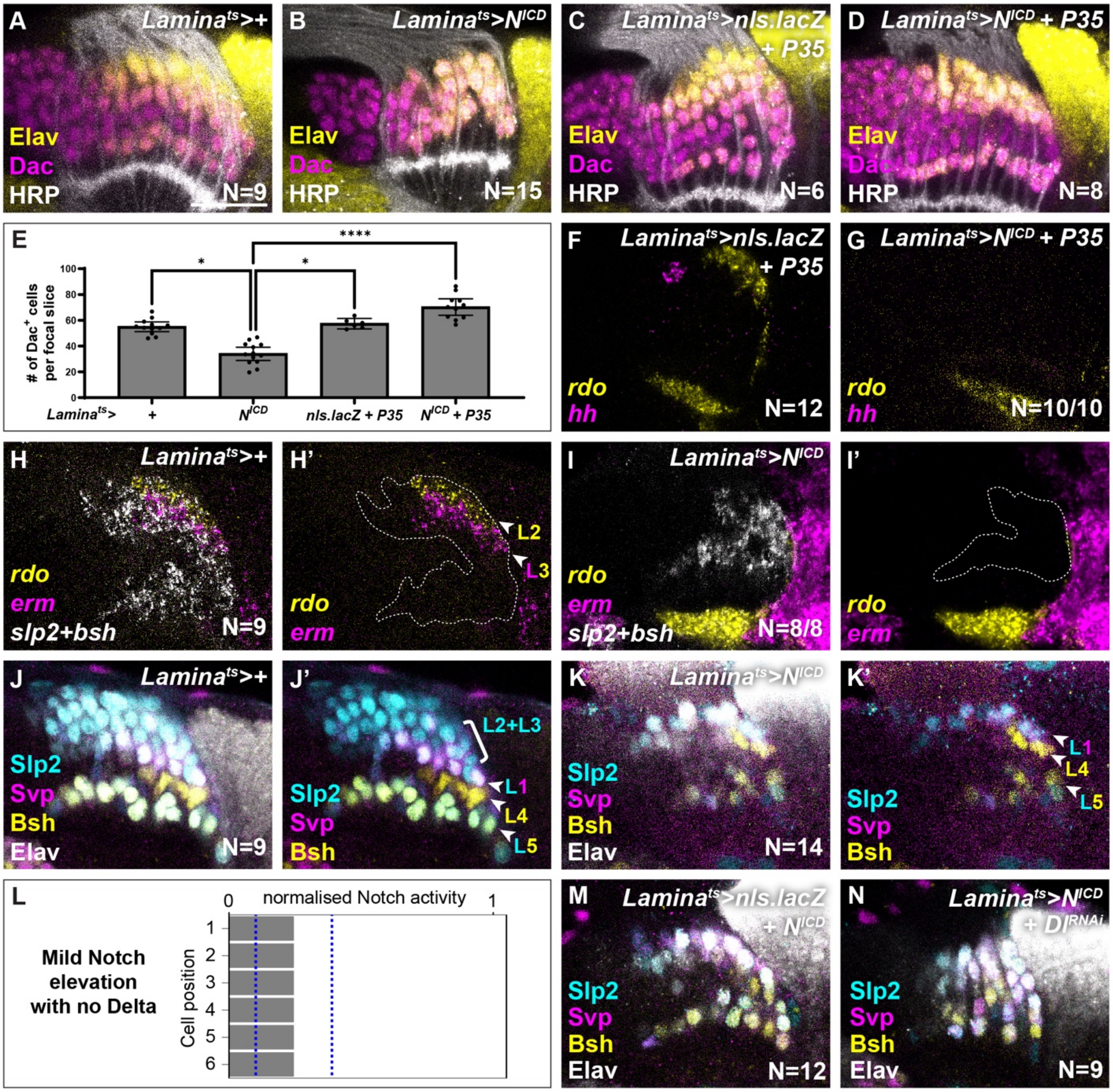

**Fig. S5: Lamina-wide $N^{ICD}$ overexpression reduces lamina size and skews surviving neurons towards L1 and L4 fates**

**(A-D)** Dac (magenta), Elav (yellow) and HRP (white) in **(A)** a control, **(B)** when $N^{ICD}$ is overexpressed in the lamina, **(C)** when nls.lacZ and P35 are expressed in the lamina (titration control) and **(D)** when $N^{ICD}$ and P35 are expressed in the lamina.

**(E)** Quantification of the number of Dac+ cells per focal slice for (A-D). One-way ANOVA with Dunn's multiple comparisons test, $p^{*}<0.05$, $p^{****}<0.0001$.

**(F, G)** *hh* (magenta) and *rdo* (yellow) expression in optic lobes when **(F)** nls.lacZ and P35, and **(G)** $N^{ICD}$ and P35 are expressed throughout the lamina. *hh* expression was absent in **(G)** in 10/10 lobes.

**(H, I)** Optic lobes labelled for *rdo* (yellow) and *erm* (magenta) to mark L2 and L3 neurons, and *slp2+bsh* (white) to mark all neurons in **(H)** control and **(I)** lamina-wide $N^{ICD}$ over-expression. *rdo* and *erm* were absent in **(I)** in 8/8 lobes.
**(J, K)** Elav (white) labelling all neurons, together with Slp2 (cyan), Svp (magenta) and Bsh (yellow) to distinguish L2/3s (Slp2+ only), L1s (Slp2+Svp+ only), L4s (Bsh+ only) and L5s (Slp2+Bsh+ only) in **(J)** control and **(K)** lamina-wide $N^{ICD}$ over-expression.
**(L)** Notch signalling levels normalised to $\beta_N/\gamma_N$ in each cell at the time of fate specification from simulations of mild Notch elevation combined with no Delta, along with a schematic of the final cell fates.
**(M, N)** Optic lobes labelled with Slp2 (cyan), Svp (magenta), Bsh (yellow) and Elav (white) in a **(M)** control and when **(N)** $N^{ICD}$ is co-expressed with $Dl^{RNAi}$.
Scale bar = 20 µm

#### Materials and Methods

##### *Drosophila* stocks and maintenance

*Drosophila melanogaster* strains and crosses were reared on standard cornmeal medium and raised at 25°C or shifted from 18°C to 29°C for genotypes with temperature-sensitive Gal80 (Gal80$^{ts}$). Table S1 in the supplementary materials includes details of the full experimental genotypes associated with each figure and supplementary figure panel and the specific temperatures used to rear them.

We used the following mutant and transgenic flies, including stocks obtained from the Bloomington Drosophila Stock Center (BDSC; NIH P40OD018537) and Vienna Drosophila Resource Center (VDRC), in combination or recombined in this study (see TableS1 for more details; {} enclose individual genotypes, separated by commas):
*{Dl$^{2.6}$-GFP}* (*1*), *{;E(spl)mγ-GFP;}* (a gift from S Bray), *Canton S*, {*;tub-Gal80$^{ts}$; repo-Gal4/TM6B*} (BDSC: 7415), {*;UAS-nls.LacZ*} (BDSC: 3955), *{;;UAS-nls.LacZ}* (BDSC: 3956), *{;UAS-EGFR$^{DN}$; UAS-EGFR$^{DN}$}* (BDSC: 5364), {*ey-Gal80; sp/CyO;*} (BDSC: 35822), {*;Gal80ts; TM2/TM6B*} (BDSC: 7108), {*$w^{1118}$;; R27G05-Gal4*} (lamina-Gal4; BDSC: 48073), *{$y^1$,sc$^*$,$v^1$,sev$^{21}$;;UAS-pnt-RNAi}* (BDSC: 35038), {*ywhsflp122, tub-Gal4, UAS-nlsGFP; FRT42D, Gal80/CyO*;} (MARCM42D, a gift from M. Amoyel), {*$y^1$; FRT42D, ptc$^{S2}$/CyO*} (BDSC: 6332), *{$w^{1118}$;;UAS-Notch-RNAi}* (Vienna Drosophila Resource Center; VDRC: 27229), *{UAS-NICD}* (a gift from S. Bray), *{y,w; R64B07-Gal4;}* (L5-Gal4; BDSC: 71106), *{ $y^1$ sc$^*$ $v^1$ sev$^{21}$;; UAS-Dl-RNAi}* (BDSC: 34322), *{$y^1$, w$^*$; UAS-NICD/SM6a}* (BDSC: 94074), *{$y^1$, w$^*$;; hh-Gal4/TM3, Sb$^1$, Ser$^1$}* (BSDB: 67493), *{UAS-His2Av::eGFP}* (BSDB: 93904), *{;UAS-p35;}* (BSDB: 5072), *{;UAS-Dl-RNAi;}* (VDRC: 109491).

##### MARCM clone induction

To induce *ptc$^{S2}$*, *ptc$^{S2}$; Notch-RNAi* and *ptc$^{S2}$; N$^{ICD}$* MARCM clones we heat-shocked larvae 3 days after egg laying (AEL) at 37°C for 20-30 minutes. All MARCM crosses were raised at 25°C until dissection at 0-5 hr After Puparium Formation (APF).

##### Immunohistochemistry, antibodies and microscopy

Eye-optic lobe complexes were dissected from female and male pupae (0–5 hr APF) in 1× phosphate-buffered saline (PBS). Samples were then fixed in 4% formaldehyde for 20 min and washed in 1× PBS with 0.5% TritonX (PBSTx) before incubating with primary antibodies diluted in PBSTx with 5% normal donkey serum (blocking solution) for two nights at 4°C. Samples were then washed in PBSTx, incubated in secondary antibodies diluted in blocking solution, washed in PBSTx and mounted in SlowFade (Life Technologies).

We used the following primary antibodies in this study: rat anti-Elav (1:100, Developmental Studies Hybridoma Bank; DHSB), rabbit anti-GFP (1:500, Thermofisher), guinea pig anti-Dpn (1:500, a gift from C. Desplan), rabbit anti-Dcp-1 (1:100, Cell Signaling), guinea pig anti-Slp2 (1:100, a gift from C. Desplan), mouse anti-Svp (1:20, DHSB), rabbit anti-Bsh (1:500, a gift from C. Desplan), mouse anti-Dac$^{2-3}$ (1:100, DHSB), AlexaFluor405-conjugated goat anti-HRP (1:50, Jackson Immunolabs) and AlexaFluor647-conjugated goat anti-HRP (1:200, Jackson Immunolabs).

Images were acquired using Zeiss 800 and 880 confocal microscopes with 40X objectives.

##### *In situ* hybridisation chain reaction (HCR)

To assess *Hey, Dl, svp, rdo, erm, slp2, bsh* and *hh* transcript expression *in vivo* we performed *in situ* hybridisation chain reaction (HCR). We used antisense probe pairs with the corresponding

initiator sequences for amplifiers B1, B3 and B5 (*2*), which we obtained as DNA oligos from Thermo Fisher (at 100µM in water and frozen). All probe sequences used in this study are included as a supplementary file (HCR Probe sequences.xls).
Eye-optic lobe complexes were dissected, fixed, and permeabilised as above. Samples were incubated in probe hybridisation buffer at 37°C for 30 min, followed by overnight incubation with probes (0.01µM) at 37°C. Samples were washed four times for 15min at 37°C with probe wash buffer, then once for 10min with 5×saline-sodium citrate with 0.001% Tween 20 solution (SSCT). A 20×SSCT solution was prepared in distilled $H_2O$, 58.44g/mol sodium chloride, 294.10g/mol sodium citrate, pH adjusted to 7 with 14M hydrochloric acid, with 0.004% Tween 20. Samples were incubated in an amplification buffer for 10 min before addition of snap-cooled hairpins. Hairpins H1 and H2 were snap-cooled (heated to 95°C for 90s and cooled to room temperature for 30min) separately to avoid hairpin oligomerisation. Approximately 12pmol of each hairpin was added to samples in amplification buffer and incubated in darkness overnight at room temperature. Following amplification, samples were washed in SSCT for ten minutes and then incubated in darkness at room temperature with 1:100 dilution of DAPI (Sigma: D9542) for 45 min. Samples were washed in 1× PBS for 15min and mounted as previously described.

**HCR with immunohistochemistry**
To visualise transcript and protein expression concurrently, we dissected, fixed and permeabilised eye-optic lobe complexes as above for HCR. Samples were incubated with probes and hairpins as above. Samples were then washed with 10 min in SSCT and re-fixed in 1× PBS with 2% formaldehyde for 20 min, washed in PBSTx for two hours, blocked in 5% normal donkey serum, and incubated in primary antibodies diluted to twice the normally used concentration in the blocking solution overnight at 4°C. Samples were then washed in PBSTx for one hour, incubated in double the normally used concentration of secondary antibodies diluted in the blocking solution, washed in PBSTx and mounted as above.

**Quantifications and statistical analysis**
All quantifications were performed blinded to experimental condition.

***Dl* mean fluorescence intensity measurements**
In Fiji-ImageJ (*3*) we used the free hand selection tool to draw a region of interest (ROI) around lamina cells (marked with DAPI; excluding the preassembly domain), and the lamina plexus. We then measured the mean fluorescence intensity (MFI) of *Dl* transcripts labelled by HCR in each ROI for 10 focal slices (step size = 1 µm) located centrally in the lamina and normalised the *Dl* MFI of the lamina to the MFI of the lamina plexus to account for differences in background signal. We then plotted the average normalised MFI for each optic lobe in GraphPad Prism.

**Dcp-1 quantifications**
We selected 30 focal slices of the central lamina in Fiji-ImageJ (step size = 1µm) and quantified the number of Dcp-1+ Dac+ cells within the lamina.

***hh* quantification**
For 20 central focal slices of the lamina (step size = 1µm) we quantified the number of nuclei marked by DAPI and positive for *hh* transcripts labelled by HCR in Fiji-ImageJ.

**Dac quantifications**
We selected the 5 central focal slices in Fiji-ImageJ (step size = 1µm) and quantified the number of Dac positive cells within the lamina (not including the preassembly domain).

#### Cell-type quantifications

***rdo, erm:*** In Fiji-ImageJ, we chose the 10 most centrally located focal slices (step size = 1µm) and measured the area of the lamina that was positive for both *slp2* and *bsh* in order to record total area occupied by neurons within these slices. Next, to establish the proportion of L2 and L3 neurons, we measured the area occupied by *rdo*+ *erm*- cells (L2) and *erm*+ *rdo*- cells (L3) relative to the total neuronal area. We then calculated the proportion of L2s and L3s relative to controls and plotted these in GraphPad Prism.
Note that to determine the identity of GFP+ MARCM clones, we chose a centrally located focal slice and quantified the number of *rdo*+ *erm*- cells (L2 identity) and *erm*+ *rdo*- cells (L3 identity) in older columns only (*i.e.,* from the $7^{th}$ column onwards). We then calculated the proportion of L2s and L3s relative to controls and plotted these in GraphPad Prism.

**Slp2, Svp, Bsh:** L2 and L3 neurons cannot be distinguished using this combination of markers, therefore, we quantified the number of L2 and/or L3s, L1s, L4s, and L5s in the 10 most centrally located focal slices (step size = 1µm). We calculated the proportion of each lamina neuron type relative to the total number of neurons (Elav positive cells) and plotted these in GraphPad Prism.

#### Statistical analysis

We used Fiji-ImageJ to process and quantify confocal images as described below. We used GraphPad Prism 9, and R (version 4.4.2) software to perform statistical tests. In all graphs, whiskers indicate the confidence interval. We show individual data points in all statistical plots; when individual points are not clearly visible (e.g., due to overlap), the corresponding N values are indicated on the graph. We used Adobe Photoshop and Adobe Illustrator software to prepare figures.

**Table S1: Full experimental genotypes and experimental conditions associated with each figure panel.** (Note that only female genotypes are listed though both sexes were included in our analyses)

| **Figure** | **Panel** | **Genotype** | **Conditions** |
|---|---|---|---|
| 1 | B, C, J, K | *Canton S* | Raised at 25°C. |
| 1 | D | *;E(spl)mγ-GFP;* | Raised at 25°C. |
| 1 | F | *y,w,hsflp$^{122}$/+; tub-Gal80$^{ts}$/UAS-nls.lacZ; repo-Gal4/UAS-nls.lacZ* | Raised at 18°C for 8 days and then shifted to 29°C for 48 hours. |
| 1 | G | *y,w,hsflp$^{122}$/+; tub-Gal80$^{ts}$/UAS-EGFR$^{DN}$; repo-Gal4/UAS-EGFR$^{DN}$* | Raised at 18°C for 8 days and then shifted to 29°C for 48 hours. |
| 1 | L, M | *y,w,hsflp$^{122}$/+, tub-Gal4, UAS-nls.GFP; FRT42D, tub-Gal80 /FRT42D, ptc$^{S2}$;* | Raised at 25°C before and after heat shocking. See MARCM clone induction for more detail. |
| 1 | N | *y,w,hsflp$^{122}$/+, tub-Gal4, UAS-nls.GFP; FRT42D, tub-Gal80 /FRT42D, ptc$^{S2}$; UAS-Notch$^{RNAi}$* | Raised at 25°C before and after heat shocking. See MARCM clone induction for more detail. |
| 1 | O | *y,w,hsflp$^{122}$/+, tub-Gal4, UAS-nls.GFP; FRT42D, tub-Gal80 /FRT42D, ptc$^{S2}$; UAS-N$^{ICD}$* | Raised at 25°C before and after heat shocking. See MARCM clone induction for more detail. |
| 1 | Q | *w$^{1118}$; R64B07-Gal4/UAS-nls.lacZ;* | Raised at 25°C. |
| 1 | R | *w$^{1118}$; R64B07-Gal4/UAS-N$^{ICD}$; UAS-N$^{ICD}$/+* | Raised at 25°C. |
| 2 | A, D, G | *ey-Gal80/+; Gal80$^{ts}$/+; R27G05-Gal4/+* | Raised at 18°C for 8 days and then shifted to 29°C for 48 hours. |
| 2 | B, E, H | *ey-Gal80/+; Gal80$^{ts}$/+; R27G05-Gal4/UAS-N$^{RNAi}$* | Raised at 18°C for 8 days and then shifted to 29°C for 48 hours. |
| 3 | A, B | *Canton S* | Raised at 25°C. |
| 3 | C | *;;Hh-Gal4/His2AveGFP* | Raised at 25°C. |
| 3 | D | *ey-Gal80/+; Gal80$^{ts}$/+; R27G05-Gal4/+* | Raised at 18°C for 8 days and then shifted to 29°C for 48 hours. |

| | | | |
|---|---|---|---|
| 3 | E | *ey-Gal80/+; Gal80$^{ts}$/+; R27G05-Gal4/UAS-Ci$^{76}$* | Raised at 18°C for 8 days and then shifted to 29°C for 48 hours. |
| 3 | G | *ey-Gal80/+; Gal80$^{ts}$/+; R27G05-Gal4/+* | Raised at 18°C for 8 days and then shifted to 29°C for 48 hours. |
| 3 | H | *ey-Gal80/+; Gal80$^{ts}$/+; R27G05-Gal4/N$^{RNAi}$* | Raised at 18°C for 8 days and then shifted to 29°C for 48 hours. |
| 5 | B | *ey-Gal80/+; Gal80$^{ts}$/+; R27G05-Gal4/+* | Raised at 18°C for 8 days and then shifted to 29°C for 48 hours. |
| 5 | C | *ey-Gal80/+; Gal80$^{ts}$/+; R27G05-Gal4/UAS-N$^{ICD}$* | Raised at 18°C for 8 days and then shifted to 29°C for 48 hours. |
| 5 | D, F | *ey-Gal80/+; Gal80$^{ts}$/UAS-p35; R27G05-Gal4/UAS-nls.lacZ* | Raised at 18°C for 8 days and then shifted to 29°C for 48 hours. |
| 5 | E, G | *ey-Gal80/+; Gal80$^{ts}$/UAS-p35; R27G05-Gal4/UAS-N$^{ICD}$* | Raised at 18°C for 8 days and then shifted to 29°C for 48 hours. |
| 5 | L | *ey-Gal80/+; Gal80$^{ts}$/UAS-nls.lacZ; R27G05-Gal4/UAS-N$^{ICD}$* | Raised at 18°C for 8 days and then shifted to 29°C for 48 hours. |
| 5 | M | *ey-Gal80/+; Gal80$^{ts}$/UAS-Dl$^{RNAi}$; R27G05-Gal4/UAS-N$^{ICD}$* | Raised at 18°C for 8 days and then shifted to 29°C for 48 hours. |
| S1 | A | *;Dl-GFP;* | Raised at 25°C. |
| S1 | B | *ey-Gal80/+; Gal80$^{ts}$/+; R27G05-Gal4/+* | Raised at 18°C for 8 days and then shifted to 29°C for 48 hours. |
| S1 | C | *ey-Gal80/+; Gal80$^{ts}$; R27G05-Gal4/UAS-pnt$^{RNAi}$* | Raised at 18°C for 8 days and then shifted to 29°C for 48 hours. |
| S1 | D | *Canton S* | Raised at 25°C. |
| S2 | A, C, E | *ey-Gal80/+; Gal80$^{ts}$/+; R27G05-Gal4/+* | Raised at 18°C for 8 days and then shifted to 29°C for 48 hours. |
| S2 | B, D, F | *ey-Gal80/+; Gal80$^{ts}$/+; R27G05-Gal4/UAS-Dl$^{RNAi}$* | Raised at 18°C for 8 days and then shifted to 29°C for 48 hours. |
| S4 | A | *y,w,hsflp$^{122}$/+; tub-Gal80$^{ts}$/hh-sfGFP; repo-Gal4/UAS-nls.lacZ* | Raised at 18°C for 8 days and then shifted to 29°C for 48 hours. |
| S4 | B | *y,w,hsflp$^{122}$/+; tub-Gal80$^{ts}$/hh-sfGFP;* | Raised at 18°C for 8 days and then shifted to 29°C for 48 hours. |

| | | | |
|---|---|---|---|
| | | *repo-Gal4/UAS-EGFR*$^{DN}$ | |
| S5 | A, H, J | *ey-Gal80/+; Gal80*$^{ts}$*/+; R27G05-Gal4/+* | Raised at 18°C for 8 days and then shifted to 29°C for 48 hours. |
| S5 | B, I, K | *ey-Gal80/+; Gal80*$^{ts}$*/+; R27G05-Gal4/UAS-N*$^{ICD}$ | Raised at 18°C for 8 days and then shifted to 29°C for 48 hours. |
| S5 | C, F | *ey-Gal80/+; Gal80*$^{ts}$*/UAS-p35; R27G05-Gal4/nls.lacZ* | Raised at 18°C for 8 days and then shifted to 29°C for 48 hours. |
| S5 | D, G | *ey-Gal80/+; Gal80*$^{ts}$*/UAS-p35; R27G05-Gal4/UAS-N*$^{ICD}$ | Raised at 18°C for 8 days and then shifted to 29°C for 48 hours. |
| S5 | M | *ey-Gal80/+; Gal80*$^{ts}$*/UAS-nls.lacZ; R27G05-Gal4/UAS-N*$^{ICD}$ | Raised at 18°C for 8 days and then shifted to 29°C for 48 hours. |
| S5 | N | *ey-Gal80/+; Gal80*$^{ts}$*/UAS-Dl*$^{RNAi}$*; R27G05-Gal4/UAS-N*$^{ICD}$ | Raised at 18°C for 8 days and then shifted to 29°C for 48 hours. |

### Computational Modelling Methods: Morphogen and juxtacrine signalling dynamically integrate to specify cell fates with single-cell resolution

Alicia Donoghue[1,], Lewis S Mosby[2,3,4,], Inês Lago-Baldaia[1], Tamara Hodgetts[1,2], Evelina Ursu[1], Zeynep Erten[1], Zena Hadjivasiliou[2,3,4,*] and Vilaiwan M Fernandes[1]

[1] *Department of Cell and Developmental Biology, University College London, Gower Street, London, WC1E 6DE.*

[2] *Mathematical and Physical Biology Laboratory, The Francis Crick Institute, 1 Midland Road, London NW1 1AT, UK.*

[3] *Department of Physics and Astronomy, University College London, Gower Street, London WC1E 6BT, UK.*

[4] *London Centre for Nanotechnology, 19 Gordon Street, London WC1H 0AH, UK.*

[§] *These authors contributed equally to this work*

**4 Numerical Methods** **16**

**5 Supplementary Tables** **18**

**6 Supplementary Figures** **21**

### 1 Theoretical Model

#### 1.1 Equations for Hh and Delta-Notch dynamics

We have developed a theoretical model that combines the spatial and temporal profiles of Delta, Notch and Hh concentrations with the dynamics of glial signalling, which locally activate ERK signalling as described by previous studies and the *in vivo* experiments in the main text (1, 2).

Wrapping glia likely form barriers that limit Hh diffusion and Delta-Notch interactions between cells in adjacent columns. We therefore assume that there are no interactions between Delta, Notch and Hh concentrations in adjacent lamina columns, so that the system can be fully described in 1D. Therefore, we model a single array of six cells spanning the positions $x \in [0, 6\,w_{\text{cell}}]$ for cells of width $w_{\text{cell}}$, and set the location of the photoreceptor source of Hh to the positions $x \in [-w_{\text{cell}}, 0]$ ($w_P = w_{\text{cell}}$ in Fig. 4A in the main text). This formalism captures the Hh gradient generated by the photoreceptors and any local lamina sources without needing to describe the trafficking details that underlie Hh transport from the photoreceptors to the lamina.

The dynamics of Delta, Notch and Hh on this domain are described by the equations in Fig. 4A in the main text, which build on previous work (3, 4). We include ligand-independent production only when probing the effects of Notch overexpression (5, 6). We assume that Delta production is gated by the ERK activation state of a cell, as seen in experiments, which we introduce mathematically using the term $Q_i$. We set the value of $Q_i$ to 1 when the glia has passed the centre point of the $i^{\text{th}}$ cell ($x_i = ((2\,i-1)/2)w_{\text{cell}}$) and to 0 otherwise. The position of the glia is defined as $x_{\text{glia}} = x_{\text{init}} + v_{\text{glia}}\,t$, where $v_{\text{glia}}$ is the speed at which the glia grow into the lamina and $t$ is the time variable. We initialise the system with zero Delta and Notch levels in all cells ($D_i(t=0) = N_i(t=0) = 0\ \forall i$), but set the initial Hh concentration to the steady-state profile generated by only the photoreceptor source. Finally, we assume zero diffusive flux boundary conditions for the Hh gradient at $x = -w_{\text{cell}}, 6w_{\text{cell}}$, such that $\partial_x C|_{x=-w_{\text{cell}},6w_{\text{cell}}} = 0$. We explain how parameters values were chosen for our analysis in section 2 and summarize the values in table M1.

#### 1.2 Rules for Cell Fate Specification

Delta, Notch and Hh dynamics are coupled through the rules that dictate Hh expression and cell fate specification, mirroring our experimental observations. For the $i^{\text{th}}$ cell, these decisions are based on the level of local Notch activity $N_i$ and the Hh concentration averaged over the cell width $\langle C \rangle_i = \int_i C\,dx/w_{\text{cell}}$, how they compare to Notch and Hh thresholds, respectively, and how the cell fate code combines these quantities. We define these rules following our experimental observations as follows (Fig. S3B,C):

1. If $\langle C \rangle_i > C^{(\text{low})}$, $N_i < N^{(\text{low})}$ and ERK is on, then the cell activates local Hh production as long as no other cell fate has been specified (this remains switched on following further cell fate specification),

2. If $\langle C \rangle_i > C^{(\text{high})}$, $N_i < N^{(\text{low})}$ and ERK is on, then the cell adopts the L2 fate,

3. If $\langle C\rangle_i > C^{(\mathrm{high})}$, $N > N^{(\mathrm{low})}$ and ERK is on, then the cell adopts the L3 fate,
4. If $C^{(\mathrm{high})} > \langle C\rangle_i > C^{(\mathrm{low})}$, $N_i < N^{(\mathrm{high})}$ and ERK is on, then the cell adopts the L1 fate,
5. If $C^{(\mathrm{high})} > \langle C\rangle_i > C^{(\mathrm{low})}$, $N > N^{(\mathrm{high})}$ and ERK is on, then the cell adopts the L4 fate,
6. If $\langle C\rangle_i < C^{(\mathrm{low})}$ and $N > N^{(\mathrm{low})}$ at any time (regardless of ERK state) then the cell undergoes apoptosis,
7. If $\langle C\rangle_i < C^{(\mathrm{low})}$, $N < N^{(\mathrm{low})}$ and ERK is on, then the cell adopts the L5 fate.

For a system where we define only a single Notch threshold ($N^{(\mathrm{high})}$ only), these rules still apply after setting $N^{(\mathrm{low})} \equiv N^{(\mathrm{high})}$. Following a cell fate decision (other than local Hh production) a cell can no longer change its fate or initiate Hh production.

##### 1.3 Timescales of Cell Fate Commitment and Delta-Notch Inactivation

We have experimentally noted delays between glial-mediated ERK activation, cell fate decision making, and the inactivation of local Delta production (1, 2) (Fig. 1A-K in the main text). We therefore introduce relevant timescales in our theoretical model to reflect these observations.

A delay has previously been observed between the timings of ERK activation and Elav expression in lamina cells (1, 2). We have further found that Elav expression is observed before the markers for the L2 and L3 fates (Fig. 1K). We capture these delays in our theoretical model by introducing the timescale $\tau_\mathrm{c}$ that defines the delay between the time at which the wrapping glia reach the centre point of the $i^{\mathrm{th}}$ cell ($x_{\mathrm{glia}} = x_i = ((2\,i-1)/2)w_{\mathrm{cell}}$), or the time at which the outer chiasm giant glia ($\mathrm{xg}^{\mathrm{O}}$) induces ERK activation in position-6 cells, and the time at which the cell can differentiate. This delay has also been introduced for cells that undergo apoptosis, such that if cells only transiently exhibit the necessary Hh concentration and Notch activity for a duration less than $\tau_\mathrm{c}$, then apoptosis will not be triggered until the conditions are met again for a duration $\tau_\mathrm{c}$.

We also note that Delta production *in vivo* is inactivated following a time delay after glial-mediated ERK activation (Figs. 1F and S1A in the main text). For this reason, we introduce a timescale $\tau_{\mathrm{DN}}$ to our simulations after which $Q_i$ switches back from $1 \to 0$, and after which Notch can no longer be activated in a cell corresponding to inactivation of both Delta and Notch receptors.

##### 1.4 Non-Dimensional Delta-Notch and Hh Equations

We non-dimensionalised the equations describing Delta, Notch and Hh dynamics (Fig. 4A in the main text), which reduces the number of free parameters in subsequent phase-space explorations. We firstly defined a common non-dimensional timescale $\tau = k\,t$, where $1/k$ is the characteristic timescale of Hh degradation. We then non-dimensionalised the Delta and Notch levels with respect to the maximal values $\beta_D/\gamma_D$ and $\beta_N/\gamma_N$ (respectively) to obtain,

$$\frac{d\tilde{D}_i}{d\tau} = \tilde{\gamma}_D \left[ Q_i \left( \frac{\tilde{\epsilon}^n}{\tilde{\epsilon}^n + \tilde{N}_i^n} \right) - \tilde{D}_i \right], \tag{M1a}$$

$$\frac{d\tilde{N}_i}{d\tau} = \tilde{\gamma}_N \left[ \left( \frac{\left(\tilde{D}_i^{(\mathrm{in})}\right)^m}{\tilde{\sigma}^m + \left(\tilde{D}_i^{(\mathrm{in})}\right)^m} \right) - \tilde{N}_i \right], \tag{M1b}$$

where $\tilde{D}_i = D_i/(\beta_D/\gamma_D)$, $\tilde{N}_i = N_i/(\beta_N/\gamma_N)$, $\tilde{\gamma}_D = \gamma_D/k$, $\tilde{\gamma}_N = \gamma_N/k$, $\tilde{\epsilon} = \epsilon/(\beta_N/\gamma_N)$, $\tilde{\sigma} = \sigma/(\beta_D/\gamma_D)$, and $\tilde{D}_i^{(\mathrm{in})}$ is the total incoming (non-dimensionalised) Delta for cell $i$ summed over its neighbours. In this case, the steady-state Delta and Notch levels are the solutions of the implicit equations,

$$\tilde{D}_i = Q_i \left( \frac{\tilde{\epsilon}^n}{\tilde{\epsilon}^n + \tilde{N}_i^n} \right), \qquad \tilde{N}_i = \frac{\left(\tilde{D}_i^{(\mathrm{in})}\right)^m}{\tilde{\sigma}^m + \left(\tilde{D}_i^{(\mathrm{in})}\right)^m}, \tag{M2}$$

and the non-dimensionalised rates $\tilde{\gamma}_D$ and $\tilde{\gamma}_N$ purely govern the time it takes for Delta and Notch to reach steady-state.

We non-dimensionalised the Hh concentration with respect to the value $\nu_P/k$, corresponding to the maximum amplitude of a system where Hh is only produced by photoreceptor cells, and also the time $t$ with respect to the Hh degradation rate ($\tau = k\,t$). Together, these transformations generate the equation,

$$\frac{\partial \tilde{C}}{\partial \tau} = \tilde{d}\, \frac{\partial^2 \tilde{C}}{\partial x^2} - \tilde{C} + \tilde{s}_P(x) + \tilde{s}_L(\tilde{C}, \tilde{N}, Q), \tag{M3a}$$

$$\tilde{s}_P(x) = \begin{cases} 1 & \text{if } -w_{\mathrm{cell}} \le x < 0 \\ 0 & \text{otherwise,} \end{cases} \tag{M3b}$$

$$\tilde{s}_L(\tilde{C}, \tilde{N}, Q) = \begin{cases} \tilde{\nu}_L & \text{if } \tilde{N} < \tilde{N}^{(\mathrm{low})},\ \tilde{C} > \tilde{C}^{(\mathrm{low})},\ \text{\& ERK On} \\ 0 & \text{otherwise,} \end{cases} \tag{M3c}$$

where $\tilde{C} = C/(\nu_P/k)$, $\tau = k\,t$, $\tilde{d} = d/k$, $\tilde{\nu}_L = \nu_L/\nu_P$, $\tilde{N}^{(\mathrm{low})} = N^{(\mathrm{low})}/(\beta_N/\gamma_N)$, and $\tilde{C}^{(\mathrm{low})} = C^{(\mathrm{low})}/(\nu_P/k)$. Together, eqs. (M1a,M1b) describing Delta and Notch dynamics, eqs. (M3a-M3c) describing Hh dynamics, and the non-dimensionalised equation describing glia motion $x_{\mathrm{glia}} = x_{\mathrm{init}} + \tilde{v}_{\mathrm{glia}}\,\tau$ (where $\tilde{v}_{\mathrm{glia}} = v_{\mathrm{glia}}/k$) fully prescribe the state of our system at time $\tau$.

### 2 Choice of Parameters Values

In order to simulate a version of our model that can be compared to experimental data, we sought to identify suitable input parameters describing Delta-Notch and Hh dynamics and the various characteristic timescales associated with ERK activation, cell fate commitment and Delta-Notch inactivation. In this section we summarize how parameters were inferred from experimental data and how parameters we did not have access to experimentally were chosen. For parameter values that are not directly accessible from the experimental data, we performed an exhaustive phase-space exploration to find when the wild-type pattern is observed. We show that there is a wide range of parameters that yield the wild-type pattern and so our conclusions are not sensitive to our choice of parameters. The final set of parameters we used for the simulations presented in the main text is summarised in table M1.

#### 2.1 Parameters for Hh and Delta-Notch Dynamics

We next sought physiologically-relevant parameters describing Hh and Delta-Notch dynamics that: (i) recapitulate the wild-type cell fate pattern observed experimentally; and (ii) generate Hh and Notch concentrations for individual cells that are 'well-separated' from their nearest thresholds at the time of differentiation. The first of these conditions allows us to assess which ranges of parameters can reproduce our experimental results, while the latter allows us to choose a final parameter set that generates Hh and Notch concentration profiles that are not highly susceptible to the effects of noise. We define the concentration $A_i$ (corresponding to either $N_i$ or $\langle C \rangle_i = \int_i C \, dx / w_{\text{cell}}$) in cell $i$ as being 'well-separated' from its nearest threshold if it exhibits a suitably high value of the quantity,

$$R_{A_i} = \min\left(\frac{|A_i - A^{(\text{low})}|}{A_i}, \frac{|A_i - A^{(\text{high})}|}{A_i}\right). \tag{M4}$$

Qualitatively, this equation extracts the minimum relative change between the Hh or Notch level in each cell and both of the corresponding thresholds, such that a larger value corresponds to Hh or Notch levels that are further (more 'well-separated') from their nearest threshold. We then compare between systems by defining $R_A = \min_i(R_{A_i})$, which parameterises a system by the separation of its least well-separated cell. Finally, we sought systems that maximise this quantity, since the cell fate patterns of these systems should be the least susceptible to fluctuations in the Hh or Notch levels.

We identify suitable parameter sets by carrying out a sweep over the parameter ranges presented in table M3 using the methods outlined in section 4. Importantly, we find that the parameters in our simulations can be varied, sometimes over orders of magnitude, without changing the output wild-type cell fate pattern (Figs. (M1-M3)). Therefore, the results of our simulations presented in the main text do not depend strongly on the parameter set we selected using the following analysis.

##### 2.1.1 Defining Hh Parameters

In order to select suitable parameters describing Hh dynamics, we first solved the equations describing non-dimensionalised Hh dynamics (eqs. (M3a-M3c)) analytically for different combinations of the photoreceptor and lamina source terms. In the case of no lamina production ($\tilde{s}_L(\tilde{C}, \tilde{N}, Q) = 0 \; \forall x$ in eq. (M3c)), where Hh is only produced by the photoreceptors, the Hh gradient takes the form (7),

$$\tilde{C} = \begin{cases} 1 - \left(\frac{\sinh\left(6\,w_{\text{cell}}/\tilde{\lambda}\right)}{\sinh\left(7\,w_{\text{cell}}/\tilde{\lambda}\right)}\right) \cosh\left(\frac{x+w_{\text{cell}}}{\tilde{\lambda}}\right) & \text{if } -w_{\text{cell}} \le x \le 0 \\ \left(\frac{\sinh\left(w_{\text{cell}}/\tilde{\lambda}\right)}{\sinh\left(7\,w_{\text{cell}}/\tilde{\lambda}\right)}\right) \cosh\left(\frac{6\,w_{\text{cell}}-x}{\tilde{\lambda}}\right) & \text{if } 0 < x \le 6\,w_{\text{cell}}, \end{cases} \tag{M5}$$

where $\tilde{\lambda} = \sqrt{\tilde{d}}$, the photoreceptor source occupies the positions $x \in [-w_{\text{cell}}, 0]$ ($w_P = w_{\text{cell}}$ in Fig. 4A in the main text), and the lamina occupies the positions $x \in [0, 6\,w_{\text{cell}}]$. Alternatively, in the case where Hh is produced locally in the lamina only in position-1 cell (the only cell observed to produce Hh in the wild-type case, Fig. 3 in the main text), then the Hh gradient instead takes the form,

$$\tilde{C} = \begin{cases} a \cosh\left(\frac{x+w_{\text{cell}}}{\tilde{\lambda}}\right) + 1 & \text{if } -w_{\text{cell}} \le x \le 0 \\ b_1\, e^{-\left(\frac{x+w_{\text{cell}}}{\tilde{\lambda}}\right)} + b_2\, e^{\left(\frac{x+w_{\text{cell}}}{\tilde{\lambda}}\right)} + \tilde{\nu_H} & \text{if } 0 < x \le w_{\text{cell}} \\ c \cosh\left(\frac{6\,w_{\text{cell}}-x}{\tilde{\lambda}}\right) & \text{if } w_{\text{cell}} < x \le 6\,w_{\text{cell}}, \end{cases} \tag{M6}$$

where the coefficients $a, b_{1,2}, c$ are given by,

$$\begin{aligned} a &= \frac{2\,b_1\,e^{-w_{\text{cell}}/\tilde{\lambda}} + (\tilde{\nu}_L - 1)}{\cosh\left(w_{\text{cell}}/\tilde{\lambda}\right) - \sinh\left(w_{\text{cell}}/\tilde{\lambda}\right)}, \\ b_1 &= \frac{\tilde{\nu}_L \tanh\left(5\,w_{\text{cell}}/\tilde{\lambda}\right) - \xi\,(\tilde{\nu}_L - 1)\tanh\left(w_{\text{cell}}/\tilde{\lambda}\right) e^{w_{\text{cell}}/\tilde{\lambda}}}{\left(1 - \tanh\left(5\,w_{\text{cell}}/\tilde{\lambda}\right)\right) e^{-2\,w_{\text{cell}}/\tilde{\lambda}} + \xi\left(1 + \tanh\left(w_{\text{cell}}/\tilde{\lambda}\right)\right)}, \\ b_2 &= \frac{b_1\left(1 + \tanh\left(w_{\text{cell}}/\tilde{\lambda}\right)\right) e^{-2\,w_{\text{cell}}/\tilde{\lambda}} + (\tilde{\nu}_L - 1)\tanh\left(w_{\text{cell}}/\tilde{\lambda}\right) e^{-w_{\text{cell}}/\tilde{\lambda}}}{1 - \tanh\left(w_{\text{cell}}/\tilde{\lambda}\right)}, \\ c &= e^{L/\tilde{\lambda}} \left(\frac{b_1\,e^{-9\,w_{\text{cell}}/\tilde{\lambda}} + b_2\,e^{-5\,w_{\text{cell}}/\tilde{\lambda}} + \tilde{\nu}_L\,e^{-7\,w_{\text{cell}}/\tilde{\lambda}}}{\cosh\left(5\,w_{\text{cell}}/\tilde{\lambda}\right)}\right), \\ \xi &= \frac{\tanh\left(5\,w_{\text{cell}}/\tilde{\lambda}\right) + 1}{\tanh\left(w_{\text{cell}}/\tilde{\lambda}\right) - 1}. \end{aligned} \tag{M7}$$

We can now sweep over the values of the non-dimensionalised diffusivity $\tilde{d}$, the source term ratio $\tilde{\nu}_L$, and the non-dimensionalised thresholds $\tilde{C}^{(\text{low,high})}$ to identify which parameters generate a wild-type cell fate pattern. We define the wild-type Hh concentration profile and thresholds as follows: Hh concentration is at 'intermediate' levels in positions-1 and -2, and low in positions-3 through -6 when there is no lamina source; and high in positions-1 and -2, intermediate in positions-3 and -4, and low in positions-5 and -6 when Hh is also produced by the position-1 cell.

Although a large range of parameters recapitulate the wild-type pattern, we observe strong negative correlation between the parameters $\tilde{d}$ and $\tilde{\nu}_L$ that generate the correct wild-type cell fate pattern (Fig. M1a). Qualitatively, increasing $\tilde{d}$ increases the effective decay length of the Hh gradient, and larger values of $\tilde{\nu}_L$

correspond to a higher contribution of the lamina Hh source relative to the photoreceptors. Our data therefore indicate that larger decay lengths require smaller values of $\tilde{\nu}_L$ to generate the wild-type cell fate pattern, and vice versa. This is the result of position-1 and -2 cells being in the intermediate Hh domain when Hh is only being produced by the photoreceptor cells, but needing to be shifted to the high Hh domain when Hh is also produced by the position-1 cell in the lamina. For small decay lengths (small $\tilde{d}$ and fast decaying Hh concentration gradients) a large enough local lamina Hh contribution (large $\tilde{\nu}_L$) is needed to shift both position-1 and position-2 cells above the intermediate Hh threshold. Conversely, for large decay lengths (large $\tilde{d}$ and slowly decaying Hh concentration gradients) the local lamina Hh contribution should not be so large that is also shifts position-3 cells into the high Hh domain ($\tilde{\nu}_L$ should not be too high). This reasoning explains the negative correlation between $\tilde{d}$ and $\tilde{\nu}_L$ needed to yield the wild-type pattern.

These constraints also explain the observed correlations between the dynamical parameters and the Hh thresholds: flatter Hh gradients (larger $\tilde{d}$, smaller $\tilde{\nu}_L$) require lower values of $\tilde{C}^{(\mathrm{high})}$ and higher values of $\tilde{C}^{(\mathrm{low})}$ to ensure that the Hh gradient can define the three regions of high, intermediate and low Hh levels following production of Hh by position-1 cells in the lamina. The reverse is true for sharper Hh gradients (smaller $\tilde{d}$, larger $\tilde{\nu}_L$) that require higher values of $\tilde{C}^{(\mathrm{high})}$ and lower values of $\tilde{C}^{(\mathrm{low})}$, leading to more separated thresholds.

The Hh parameters that generated the wild-type cell fate pattern and exhibited Hh levels that were the best separated from their nearest thresholds are indicated by a cross in Fig. M1a and are presented table M3. The resulting Hh gradient exhibits Hh levels in position-1 to -3 cells that are approximately equidistant from their nearest thresholds in log-space when there is only a photoreceptor Hh source, and Hh levels in position-2 to -5 cells that are also approximately equidistant from their nearest thresholds in log-space when there is local lamina production in only position-1 cells (Fig. M1b,c). This follows from the approximately exponential forms of eqs. (M5,M6). Additionally, none of the Hh levels of individual cells overlap with their nearest thresholds within the standard deviation of the Hh levels across the span of the cell. Therefore, the Hh parameter sets we have used for our analysis in the main text were selected so that: (i) the wild-type Hh domains with and without the lamina source are recreated; and (ii) our results are not sensitive to low-level noise in the local Hh concentration.

##### 2.1.2 Defining Delta and Notch Parameters

To determine parameters for Delta-Notch dynamics, we simulated the equations describing the non-dimensionalised Delta and Notch levels (eqs. (M1a,M1b)) for different values of the fractional Delta signalling threshold $\tilde{\sigma}$, the fractional Notch signalling threshold $\tilde{\epsilon}$, and the non-dimensionalised thresholds $\tilde{N}^{(\mathrm{low,high})}$ that define cell fate. Importantly, Delta and Notch levels are independent of the dynamics of the Hh gradient, and eq. (M2) shows that the steady-state Delta and Notch levels are independent of the non-dimensionalised degradation rates $\tilde{\gamma}_N$ and $\tilde{\gamma}_D$, which instead prescribe system timing. In this case, we define a wild-type Notch pattern as: (i) zero in position-1 cells, high in position-2 cells, low in position-3 cells, high in position-4 cells, high in position-5 cells and zero in position-6 cells at the time of cell fate specification; and (ii) non-zero (low or high) in cells at positions-2 through -5 at all times following the differentiation of cells at positions-1 and -6. This latter condition ensures that position-2 through -5 cells exhibit non-zero levels of Notch activity(Fig. 1 in the main text) and so do not produce Hh, as observed in wild-type experiments (Fig. 3 in the main text). The Notch levels in cells at positions-1 and -6 are always zero at the time they differentiate, as glial-induced Delta production has not yet been activated in any other cell, and both cells at positions-2 and -5 exhibit high Notch levels in the wild-type case as a result of trans-activation by Delta produced by cells at positions-1 and -6. Together, this means that only the Notch levels of positions-2 to -4 cells at the time of differentiation need to be tested against conditions (i) and (ii) defined above.

The capacity to generate graded Notch activity in cells across each column was central to obtaining the wild-type pattern in our simulations. As we note above and in the main text, this is because zero Notch activity combined with intermediate Hh signalling promote Hh production in the lamina, such that cells in positions-2 through -6 are capable of producing Hh in the absence of Notch. A way to mitigate this is for cells to exhibit either high or low (but non-zero) levels of Notch, which in turn inhibits Hh production in the lamina as seen in our experiments (Fig. 3 in the main text). Theoretically, cells that produce high levels of Delta can still exhibit non-zero Notch levels when low Notch is not sufficient to inhibit Delta expression. In dynamical systems terms, this represents a stable state for $(\tilde{D}_i, \tilde{N}_i)$ where both $\tilde{N}_i$ and $\tilde{D}_i$ can be non-zero.

For instance, according to eq. (M1b), non-zero Notch activity in position-3 requires a value of $\tilde{\sigma}$ small enough to ensure that the low Delta levels in position-2 are translated into intermediate Notch levels in position-3, but not so small that the low Delta levels in position-2 are instead translated into high Notch levels in position-3 (Fig. M4a,b). Similarly, eq. (M1a) indicates that non-zero Notch activity in position-3 also requires a value of $\tilde{\epsilon}$ small enough to limit the amount of Delta produced in position-2 and therefore the amount of Notch activity in position-3. In contrast, the value of $\tilde{\epsilon}$ cannot be so small that not enough Delta is produced in position-2 and the Notch activity in position-3 falls below the intermediate threshold (Fig. M4c,d). In the latter case, position-3 cells will be in an effectively zero ($\tilde{N}_i < \tilde{N}^{(\text{low})}$) Notch and intermediate Hh state, which results in the incorrect activation of local Hh production, ultimately leading to the L2, L3, L2, L3, X, L5 cell fate pattern (Fig. M4c,d). Although these explanations are focused on the effects on the Notch level at position-3, Fig. M4b shows that varying these parameters can also affect the Notch levels in other cells as well.

Consistent with the reasoning above, our simulations indicate that although a large range of values of the fractional signalling thresholds $\tilde{\sigma}$ and $\tilde{\epsilon}$ can generate the wild-type cell fate pattern, there is a strong correlation between them. In contrast, the non-dimensionalised thresholds $\tilde{N}^{(\text{low,high})}$ can take a comparatively wide range of values independently of one another. Outside the ranges shown in Fig. M4, Notch is either 0 or 'high' also explaining the two activity regimes discussed and shown Fig. 4C in the main text.

The Delta and Notch parameters that generated the correct wild-type cell fate pattern and exhibited Notch levels that were the best separated from their nearest thresholds are indicated by a cross in Fig. M2a and are presented table M3. In this case, only the separation between the Notch levels in positions-2 through -4 and their nearest neighbours are quantified at the time of cell fate specification ($R_N = \min_{i=[2,3,4]}(R_{N_i})$). The resulting Notch profile exhibits $\tilde{N}_{2,4} \simeq 1$, which always ensures maximal separation from the nearest Notch threshold $\tilde{N}^{(\mathrm{high})}$ (Fig. M2b). The Notch level in position-3 is instead approximately equidistant from the thresholds $\tilde{N}^{(\mathrm{low,high})}$. This corresponds to a system that is unlikely to generate an incorrect wild-type cell fate pattern when there is relatively low-level noise in local Notch levels.

###### 2.1.3 Notch Overexpression

To investigate the impact of experimentally overexpressing $\mathrm{N}^{\mathrm{ICD}}$ in the theoretical model, we added a baseline Delta-independent Notch production rate, $\beta_{N_0}$, to the equation describing Notch dynamics in Fig. 4A in the main text. We then used our analytical model to assess how different degrees of Notch overexpression affect cell fate patterning, as shown in Fig. 5A in the main text.

Following the non-dimensionalisation procedure defined in section 1.4 earlier in this document, the equation describing Notch dynamics with baseline production can be written,

$$\frac{d\tilde{N}_i}{d\tau} = \tilde{\gamma}_N \left[ \tilde{\beta}_{N_0} + \left( \frac{\left(\tilde{D}_i^{(\mathrm{in})}\right)^m}{\tilde{\sigma}^m + \left(\tilde{D}_i^{(\mathrm{in})}\right)^m} \right) - \tilde{N}_i \right], \tag{M8}$$

where $\tilde{\beta}_{N_0} = \beta_{N_0}/\beta_N$.

To derive suitable values for $\tilde{\beta}_{N_0}$ equivalent to 'mild' and 'strong' Notch overexpression (Fig. 5A in the main text), we first consider the limit where $D_i^{(\mathrm{in})} = 0 \; \forall i$, i.e. in the absence of incoming Delta. This corresponds to the minimum observable Notch levels for a system with a baseline, cell-autonomous Notch production and no Delta-driven Notch activation. In this limit, the Notch concentration is uniform throughout the lamina, reflecting the absence of lateral inhibition, with magnitude $\tilde{N}_i = \tilde{\beta}_{N_0}$. It follows that the Notch activity of cells 1 and 6, which is usually zero, is sufficient to inhibit local Hh production when $\tilde{\beta}_{N_0} \geq \tilde{N}^{(\mathrm{low})}$, or in dimensional terms when $\beta_{N_0}/\gamma_N \geq N^{(\mathrm{low})}$. Therefore, we can simulate mild Notch overexpansion by setting $\beta_{N_0} \geq \gamma_N N^{(\mathrm{low})}$ and removing Delta production in all cells at all times. Similarly, for 'strong' Notch overexpression we need $\tilde{\beta}_{N_0} \geq \tilde{N}^{(\mathrm{high})}$ and therefore need to set $\beta_{N_0} \geq \gamma_N N^{(\mathrm{high})}$ while removing Delta production in all cells at all times.

Maintaining Delta production has the effect of pushing the Notch levels of the position-2 cell into the high Notch activity state independently of the value of $\beta_{N_0}$, since production of Delta in the position-1 cell still drives Notch activity in the position-2 cell. This explains why position-1 and -2 cells take on the L1 and L4 fates under mild Notch overexpression in our simulations when Delta production and Delta-driven activation of Notch in neighbouring cells is maintained in addition to the Notch overexpression (Fig. 5A in the main text).

#### 2.2 Timescales of Cell Fate commitment and Delta-Notch inactivation

In Section 1.3 of this document, we introduced a timescale $\tau_c$ that defines the delay between the time at which glia reache the centre point of the $i^{\text{th}}$ cell ($x_{\text{glia}} = x_i = ((2\,i-1)/2)w_{\text{cell}}$) and the time at which that cell can differentiate. In order to probe the values of $\tau_c$ that could lead to the observed wild-type cell fate pattern, we simulated Hh, Delta and Notch dynamics when $\tau_c = f\,w_{\text{cell}}/v_{\text{glia}}$ for various values of $f$ (Fig. M5). We found that the cell fate of the position-1 cell is disrupted when $\tau_c > w_{\text{cell}}/v_{\text{glia}}$; in this case Delta production is initiated in the position-2 cell before the position-1 cell has adopted its fate, increasing the Notch level in the position-1 cell past the $N^{(low)}$ threshold and resulting in the L3 fate. Similarly, we find that the position-1 cell incorrectly adopts the L1 fate when $f$ is below 0.25. In this case, the position-1 cell adopts its fate before the Hh concentration reaches the $C^{(high)}$ threshold following the initiation of local Hh production, resulting in the position-1 cell adopting a fate associated with intermediate Hh. The cell fates of the remaining positions appear robust to changes in $\tau_c$ (Fig. M5). We therefore use a value of $\tau_c = w_{\text{cell}}/v_{\text{glia}}$, equivalent to setting $f = 1$, but any value of $f$ that allows sufficient time for Hh build-up following local lamina production and that satisfies $f \leq 1$ will yield a wild-type cell fate pattern. These findings highlight the timekeeping role of wrapping glia during lamina patterning, discussed in the main text.

We have also introduced a timescale $\tau_{\text{DN}}$ that defines the delay between the time at which wrapping glia reach the centre point of the $i^{\text{th}}$ cell ($x_{\text{glia}} = x_i = ((2\,i-1)/2)w_{\text{cell}}$), or the time at which the outer chiasm giant glia ($\text{xg}^{\text{O}}$) induces ERK activation in position-6 cells, and the time at which Delta production is inactivated in that cell. To estimate this timescale experimentally, we noted, based on the rate of column assembly, the delay between Delta production being switched on and switched back off again (Figs. 1F and S1A) in a cell is approximately 3.6 – 4.6 hours (8). We also estimate wrapping glial velocity to be 2.1 – 3.0 µm/hour (we used the average value of 2.55 µm/hour in our simulations, see table M1) (9). In this case, the glia moves approximately 7.5 – 13.8 µm during the delay, equivalent to roughly 2 – 3 cell widths. This implies that $2\,w_{\text{cell}}/v_{\text{glia}} \lesssim \tau_{\text{DN}} \lesssim 3\,w_{\text{cell}}/v_{\text{glia}}$, we have therefore employed $\tau_{\text{DN}} = 2\,w_{\text{cell}}/v_{\text{glia}}$ in our analysis.

We finally assume that the Hh, Delta and Notch profiles relax to their steady-states faster than the time it takes for glia to travel between cells ($w_{\text{cell}}/v_{\text{glia}}$). We have used values of the Hh degradation rates of $\sim 10(v_{\text{glia}}/w_{\text{cell}})$, which are also of the same order of magnitude as previously calculated values for Hh degradation (10). We assume similar degradation rates for Delta and Notch. All timings in simulations were rescaled by the characteristic glia timescale $w_{\text{cell}}/v_{\text{glia}}$, such that only the relative values of the relaxation timescales are important (see the section 4 later in this document). We also explore how slower Delta-Notch dynamics impact our conclusions in section 3.2 later in this document.

### 3 Additional simulation results

In this section, we elaborate and expand on key results of the theoretical model presented in the main text. We focus on the roles of the relative timescales of cell fate specification, Delta-Notch inactivation, and coupling of glial signalling dynamics to different cell behaviours.

##### 3.1 Variation in Delay Timescales

We explored the effects of varying the time delay between glial-mediated ERK activation and cell fate specification, $\tau_c$, on cell fate patterning in section 2.2 earlier in this document in order to guide our parameterisation, as this timescale was not directly available to us experimentally. In order to investigate the effects of varying the time delay between glial-mediated ERK activation and Delta and Notch inactivation, $\tau_{DN}$ , on cell fate specification we set $\tau_{DN} = f\, w_{cell}/v_{glia}$, and simulated Hh, Delta and Notch dynamics for various values of $f$ (Fig. M6). When $f = 0$, each cell immediately inactivates Delta production following its activation, leading to the trivial case of no Delta production. This leads to all cells exhibiting no Notch activity at the time they are reached by the glia, resulting in activation of local Hh production in positions-1 through -5 cells, and each of these cells adopting the L2 fate (Fig. M6b). This cascade of Hh production is similar to that observed in Notch knockdown experiments (Fig. 3H in the main text). Setting $f = 1$ generates largely the same cell fate pattern, since Delta production is still not switched on in adjacent cells at the time when any cell differentiates, resulting in them exhibiting zero Notch activity. Note however, that in this case the position-5 cell undergoes apoptosis since it exhibits low Hh and high Notch levels for a time $\tau_c$ (Fig. M6b). When $f \geq 2$ the cascade of Hh production no longer occurs, since the high Delta level in position-1 drives a high Notch activity level in position-2 at the time when it differentiates, preventing local Hh production (Fig. M6b). Therefore, the wild-type cell fate pattern requires $f = 2$ (Fig. M6a). When $f > 2$, position-4 cells incorrectly adopt the L1 fate because both positions-2 and -4 cells produce Delta simultaneously, ultimately generating a high Notch level in position-3 that drives the Notch level in position-4 to remain below $N^{(\mathrm{high})}$ (Fig. M6c,d). Since this result occurs for any $f > 2$, only $f = 2$ can generate the wild-type cell fate pattern, which agrees with our experimentally-derived value (see section 2.2 earlier in this document).

##### 3.2 Variation in Delta-Notch degradation rates

Since our methods for obtaining Delta and Notch parameters in section 2.1.2 did not constrain the possible values of the Delta and Notch degradation rates, we explored the effects of varying the Delta and Notch degradation rates in simulations (Figs. M3 & M7). We found that the system parameters we used throughout the analysis shown in the main text (see table M1) allow for significant slowing of Delta-Notch dynamics without any changes in the observed cell fate pattern. However, for slow Delta-Notch degradation rates ($\gamma_D \leq 10^{-2}\,\gamma_D^{(0)}$ and $\gamma_N \leq 10^{-2}\,\gamma_N^{(0)}$, where $\gamma_D^{(0)}$ and $\gamma_N^{(0)}$ are the values presented in table M1), the Notch levels no longer reach steady state by the time of cell fate specification, which can disturb cell fate specification.

Dynamics even slower than this lead to a cascade of local Hh production akin to that observed in Notch knockdown experiments (Fig. M7a). In this case, for position-2 to -4 cells, Delta production in adjacent cells causes the local Notch activity level to increase, but this is too slow to cross the low Notch threshold ($\tilde{N}^{(\mathrm{low})}$)

before that cell differentiates, which occurs after the delay $\tau_c$ (Fig. M7b,c). These behaviours only start to appear when the degradation rates of Delta and Notch are 100-fold smaller than the values we used for the simulations in the main text, corresponding to the order of 10 hours, which is higher than the typical timescales of Delta-Notch dynamics (6). We therefore conclude that our findings are not sensitive to variation in the degradation and dynamics of Delta-Notch signalling at physiological levels.

In order to test whether any set of system parameters could generate the correct wild-type cell fate pattern when the Delta and Notch dynamics have been significantly slowed, we repeated our phase-space analysis for Delta and Notch degradation rates of $\gamma_D = \gamma_N = 2 \times 10^{-5}$ s$^{-1}$ (Fig. M3). This corresponds to values of $\gamma_D = 10^{-2}\,\gamma_D^{(0)}$ and $\gamma_N = 10^{-2}\,\gamma_N^{(0)}$, which we have shown perturb wild-type cell fate patterning for the parameters presented in table M1 (Fig. M7). We find that slowing the rates of Delta and Notch equilibration effectively constrains, but does not move, the regions of parameter space in which the wild-type cell fate pattern can be achieved (Fig. M7a). We have compared the time series of the Delta and Notch levels for the parameter sets that generate the correct wild-type cell fate pattern and exhibited Notch levels that were the best separated from their nearest thresholds for both the 'fast' and 'slow' phase-space analyses in Fig. M3b, and have presented the corresponding parameters in table M3. Although both systems can generating the correct wild-type cell fate pattern, it is clear that the Notch levels in the position-2 and -4 cells of the system with slower Delta-Notch dynamics are less well-separated from their nearest thresholds, since they do not reach the maximum values of $\tilde{N}_{2,4} \simeq 1$ exhibited in the faster system (Fig. M7b). Similarly, the systems with smaller degradation rates are more sensitive to values of $\tau_c \leq w_{\text{cell}}/v_{\text{glia}}$ of the delay timescale for cell fate specification following the arrival of the glia at a cell. This is because the Notch build up in any cell becomes slow relative to the time of cell fate specification. As a result, cells do not reach the necessary Notch activity levels to cross their endogenous thresholds before their cell fate is sealed, resulting in ectopic Notch activity levels at the time of cell fate specification and patterning errors.

#### 3.3 Decoupling Glia from Delta Production, Hh Production and Cell Fate Specification

A central conclusion in our work is that glial-mediated ERK activation is coupled to the timings of Delta production, local Hh production and cell fate decision making. We have shown that if this coupling is impaired, the wild-type cell fate pattern only emerges stochastically with low probability (Fig. 4G in the main text). Therefore, glial-induced ERK activity appears to be central to ensuring robust spatial patterning. In this section, we elaborate on how spatial patterning is impaired when we sequentially decouple glial-induced ERK activation from: (i) Delta production; (ii) Hh production; and (iii) cell fate specification. In each case, the distributions of cell fate patterns presented in Fig. S4C have been extended in Fig. M8, and the corresponding Hh production states have been added to aid understanding. The computational methods for these simulations are described in section 4 later in this document.

##### 3.3.1 Randomising the Timing of Delta Production

When we randomised the timing of activating Delta production in each cell, we noticed a misregulation of Hh production in the lamina compared to the wild-type (Fig. M8a). The most frequently observed cell fate patterns included cells adopting L2 and L3 fates throughout positions-1 to -5, with position-6 cells adopting either the L5 or the apoptotic fate (Figs. S4 & M8a). These were accompanied by an increase in the number of cells locally producing Hh (Figs. S4 & M8a), which is consistent with the increased frequency of L2 and L3 fates that correspond to the high Hh domain and necessitate a Hh source in the lamina.

The increase in the number of Hh producing cells under these conditions can be understood as follows. When Delta production is randomised, position-1 and position-2 cells can both produce Hh as long as they have no neighbour producing Delta when the glia reaches them and activates ERK, which is necessary for Hh local production (see section 1.2 earlier in this document). Production of Hh in position-1 cells can drive position-3 and -4 cells to the intermediate Hh domain, and Hh production in position-2 cells can drive positions-3 through -5 cells into the intermediate Hh domain. Following a similar logic as for cells in position-1 and -2, any other cells driven to the intermediate Hh domain are now also able to produce Hh as long as they have no neighbour producing Delta by the time the glia reaches them. These events occur with a relatively high probability when Delta production is randomised, and generate a cascade of Hh activation in position-1 through -5 cells in a similar way as seen in the Notch knockdown experiments and corresponding simulations presented in the main text (Figs. 2 & 4 in the main text). In contrast, position-6 cells always differentiate before any Hh production cascade shifts them into a high or intermediate Hh state, but they adopt the wild-type L5 fate only when the position-5 cell has not activated Delta production by the time they differentiate, and they undergo apoptosis when it has (Fig. M8a). Other cell fates are also possible with randomised timings of Delta production. For instance, Delta can be activated early in position-2 cells compared to the wild-type case, and activated at times in position-1 cells that are comparable or earlier than the wild-type case, which drive Notch above the low threshold in both cells by the time each of them experience glial-mediated ERK activation. This therefore inhibits Hh production in the lamina altogether and leads position-1 and -2 cells to adopt the L1 fate (Fig. M8a). These results suggest that the early or late activation of Delta production can prevent cells from adopting their wild-type cell fates by either causing a cascade of local Hh production within the lamina or inhibiting lamina production of Hh altogether.

##### 3.3.2 Randomising the Timing of Local Hh Production

When we randomised the timing of activating local Hh production in each cell, the distribution of possible cell fate patterns was much more constrained when compared to randomising Delta production (Figs. S4 & M8b). Note that although we assume that the timing of Hh production is glia-independent in these simulations, we still assume that intermediate-high Hh levels and zero Notch are necessary conditions for cells to produce Hh.

Under these conditions, the most frequently observed cell fate pattern involved all cells adopting either the

L2 or L3 fate across the column (Figs. S4 & M8b). This occurs when the random Hh production times in multiple cells coincides with them being in an intermediate-high Hh domain and a zero Notch level, corresponding to times before Delta production has been initiated in adjacent cells. In these cases, cells at all positions produce Hh with the same relatively high likelihood, resulting in several cells producing Hh each time we perform a single simulation. These multiple local sources are expected to elevate the Hh concentration above the high Hh threshold across the entire column, thereby leading all cells to adopt L2 or L3 fates depending on their Notch state.

The second most frequently observed cell fate pattern arises when no cells produce Hh (Figs. S4 & M8b). In this case, Hh production is not initiated in position-1 or -2 cells prior to the position-1 cell adopting its fate or the position-1 cell activating Delta production, respectively. In this scenario, position-1 and -2 cells adopt L1 and L4 fates, positions-3 through -5 cells undergo apoptosis, and position-6 cells adopt the L5 fate, as randomising the timing of local Hh production does not otherwise influence the timings of Delta production or cell fate specification. Together, these results suggest that both the early and late activation of local Hh production can prevent cells from adopting their wild-type cell fates.

##### 3.3.3 Randomising the Timing of Cell Fate Specification

When the timing of cell fate specification in each cell is randomised, the most frequently observed cell fate patterns generally either involve: (i) position-1 and -2 cells adopting the L1 or L4 fates; or (ii) position-1 cells initiating local Hh production such that position-1 and -2 cells exhibit L2 or L3 fates (Figs. S4 & M8b). This mirrors our results from section 1.3, where we demonstrated that varying the time delay for cell fate specification following glial-mediated ERK activation, $\tau_c$, governs whether position-1 cells adopt the L1 or L2 fates (Fig. (M5)). However, when cell fate specification occurs in position-1 cells before they initiate local Hh production, the Hh states and cell fates of position-2 through -5 cells are disrupted.

For example, if position-1 cells differentiate prematurely and do not initiate local Hh production, then none of the position-2 through -6 cells can ever locally produce Hh, as they either express high Notch levels or are in the low Hh domain. In this case, position-1 and -2 cells are in the intermediate Hh domain, and cells in position-3 through -6 are in the low Hh domain, as reflected by their cell fate decisions (see section 2.1.1). Alternatively, if position-1 cells produce Hh, then the wild-type cell fate pattern is possible as long as other cells do not differentiate prematurely. Premature differentiation for position-2 through -6 cells corresponds to cell fate specification prior to the Delta production being initiated in adjacent cells, which leads to ectopic Notch activity levels in these cells, and ultimately their cell fates.

Together, these results suggest that premature cell fate determination can prevent cells from adopting their wild-type cell fates. In particular, the premature differentiation of position-1 cells prevents the necessary expansion of the Hh source into the lamina to generate the three regions of distinct Hh activity, whereas premature differentiation in other cells can drive them to take on a cell fate decision before their Notch activity levels are set.

### 4 Numerical Methods

We solved key equations numerically and performed simulations using a forwards Euler scheme in the Julia programming language. Delta and Notch dynamics were quantified on a grid of 6 cells, such that position-1 and -6 cells have only one neighbour while positions-2 through -5 cells each have two neighbours. Hh dynamics were instead quantified on a spatial grid of 281 grid points spanning a length $L = 28$ µm. Before each simulation, the initial Hh gradient was first obtained by simulating Hh dynamics using the Rodas4P solver with only photoreceptor production according to the equation defined in Fig. 4A in the main text. The initial Delta and Notch concentrations were set to zero for all cells. The simulation was terminated when both the glia had reached the end of the lamina ($x_{\text{glia}} \geq x_6 = (11/2)w_{\text{cell}}$) and changes in the Delta, Notch and Hh concentrations were smaller than specified relative ($10^{-6}$) and absolute ($10^{-8}$) tolerances.

During simulations, timings were rescaled by the characteristic glia timescale $w_{\text{cell}}/v_{\text{glia}}$, such that $x_{\text{glia}} = x_i$ when $t = ((2\,i{+}1)/2)$, and after this non-dimensionalisation the simulation timestep was set to $\Delta t = 2.5{\times}10^{-6}$. Following each timestep the glia position is updated, and if the position of the glia satisfies $x_{\text{glia}} \geq x_i$ then $Q_i \to 1$ and Delta production in the $i^{\text{th}}$ cell is initiated. Cell fate decisions were made for the $i^{\text{th}}$ cell according to the rules defined in Fig. 4D,E in the main text and section 1.2 earlier in this document following a delay $\tau_{\text{c}}$ after the glia reaches it (at time $t = ((x_i + x_{\text{init}})/v_{\text{glia}}) + \tau_{\text{c}}$), and Delta and Notch production are also switched off following a delay $\tau_{\text{DN}}$ (at time $t = ((x_i + x_{\text{init}})/v_{\text{glia}}) + \tau_{\text{DN}}$).

The parameters used for the simulations shown in Figs. 4C-F & 5A in the main text are presented in tables M1 & M2. For the phase-space exploration detailed in section 2.1, parameters were sampled uniformly in log-space within the ranges specified in table M3. The parameter range chosen for Hh parameters was selected to cover diverse parameter sets, and adjusted to encapsulate the maxima of the robustness observed in Fig. M1. The parameter range chosen for Delta and Notch parameters was selected to encapsulate the parameters used for similar models in previous work ($\times 10^{\pm 2}$) (4). A total of $2 \times 10^6$ random parameter sets were sampled independently for the Hh and Notch analyses. The Hh profiles were calculated directly from eqs. (M5,M6), whereas the Delta and Notch profiles were obtained using simulations following the methods described earlier in this section. In this case, simulations were terminated when all cells had adopted a cell fate. Finally, we assumed there exists a minimum activity level below which cells cannot discern changes between a low and intermediate Hh level or a zero and low Notch level due to the presence of noise. We assume that this minimum activity level is 10% of the maximum activity level, such that the non-dimensionalised Hh and Notch cell fate decision thresholds are limited to the ranges $A^{(\text{low,high})} \in [0.1, 1.0]$. The parameters presented in table M3 represent the systems with Hh and Notch levels that are the most 'well-separated' from their nearest thresholds, as defined in eq. (M4).

Timings were randomised for each cell in the simulations shown in Fig. 4G in the main text by sampling from a Poisson distribution, using inverse transform sampling, with mean rate equal to the average rate of the relevant process (Delta production, Hh production or cell fate specification) in the wild-type case. A total of

9 999 random sets of timescales were sampled independently for each process. In each case, Delta production was still switched off following a delay $\tau_c = 2\,w_{\text{cell}}/v_{\text{glia}}$. Cell fate specification was also assumed to still occur following a delay $\tau_{\text{DN}} = w_{\text{cell}}/v_{\text{glia}}$ after the glia reaches the cell, except for in the case where the timing of cell fate specification was randomised, when no delay was enforced. Importantly, the randomised timings of cell fate specification and local Hh production refer only to the time after which cells *can* change fate provided they satisfy the conditions detailed in section 1.2 earlier in this document, and not necessarily the time at which cell fate changes occur.

The code used to generate all computational results in this work can be found at: `https://www.github.com/LMosby/MorphogenJuxtacrineIntegration`.

### 5 Supplementary Tables

| **Parameter** | **Unit** | **Description** | **Value** |
|---|---|---|---|
| $d$ | $\mu m^2\ s^{-1}$ | Hh diffusivity | **0.0711** |
| $k$ | $s^{-1}$ | Hh degradation rate | **0.0020** |
| $\nu_P$ | mol $\mu m^{-2}\ s^{-1}$ | Photoreceptor Hh production rate | **0.1000** |
| $\nu_L$ | mol $\mu m^{-2}\ s^{-1}$ | Cell Hh production rate | **0.1780** |
| $C^{(\text{low})}$ | mol $\mu m^{-2}$ | Low Hh threshold | **5.0841** |
| $C^{(\text{high})}$ | mol $\mu m^{-2}$ | High Hh threshold | **17.0238** |
| $\beta_N$ | molecules $s^{-1}$ | Delta-dependent Notch production rate | 0.1000 |
| $\gamma_N$ | $s^{-1}$ | Notch degradation rate | 0.0020 |
| $\sigma$ | molecules | Delta signalling threshold | **0.0785** |
| $m$ | – | Hill constant governing Delta signalling | 2 |
| $\beta_D$ | molecules $s^{-1}$ | Delta production rate | 0.1000 |
| $\gamma_D$ | $s^{-1}$ | Delta degradation rate | 0.0020 |
| $\epsilon$ | molecules | Notch signalling threshold | **4.8197** |
| $n$ | – | Hill constant governing Notch signalling | 3 |
| $N^{(\text{low})}$ | molecules | Low Notch threshold | **5.0650** |
| $N^{(\text{high})}$ | molecules | High Notch threshold | **19.4855** |
| $w_{\text{cell}}$ | µm | Cell width | 4.0000 |
| $v_{\text{glia}}$ | $\mu m\ s^{-1}$ | Glia velocity | $1.7708 \times 10^{-4}$ |
| $x_{\text{init}}$ | µm | Cell width | -4.0000 |

Table M1: **Simulation parameter values.** Parameter values used to generate the results of Figs. (4B-G,5A) in the main text by solving the equations defined in Fig. 4A in the main text. Bold values satisfy the conditions for optimising pattern robustness presented in table M3.

| Figure | Parameter | Unit | Value |
|---|---|---|---|
| 4C (left) | $\epsilon$ | molecules | 0.0563 |
| | $\sigma$ | molecules | 0.1515 |
| 4D | $N^{(\mathrm{low})}$ | molecules | 19.4855 |
| 5A (left) | $\beta_{N_0}$ | molecules $s^{-1}$ | 0.0246 |
| 5A (right) | $\beta_{N_0}$ | molecules $s^{-1}$ | 0.0490 |
| S3 (left) | $N^{(\mathrm{low})}$ | molecules | 19.4855 |
| S5L | $\beta_{N_0}$ | molecules $s^{-1}$ | 0.0246 |

Table M2: **Variations in parameter values.** Differenecs between the parameter values presented in table M1 and those used in perturbation simulations.

| **Non-Dimensionalised Parameter** | **Formula** | **Description** | **Value Range** | **Optimised Value** | **Optimisiation Method** |
|---|---|---|---|---|---|
| $\tilde{d}$ | $d/k$ | Re-scaled Hh diffusivity | $2.5\times10^0 - 2.5\times10^4$ | 35.5518 | Hh sweep |
| $\tilde{\nu}_L$ | $\nu_L/\nu_P$ | Fractional local Hh production | $10^{-1} - 10^3$ | 1.7796 | Hh Sweep |
| $\tilde{C}^{(\mathrm{low})}$ | $C^{(\mathrm{low})}/(\nu_P/k)$ | Fractional low Hh threshold | $0.1 - 1.0$ | 0.1017 | Hh Sweep |
| $\tilde{C}^{(\mathrm{high})}$ | $C^{(\mathrm{high})}/(\nu_P/k)$ | Fractional high Hh threshold | $0.1 - 1.0$ | 0.3405 | Hh Sweep |
| $\tilde{\sigma}$ | $\sigma/(\beta_D/\gamma_D)$ | Fractional Delta signalling threshold | $10^{-3} - 10^1$ | 0.0016 | Notch sweep (fast) |
| $\tilde{\epsilon}$ | $\epsilon/(\beta_N/\gamma_N)$ | Fractional Notch signalling threshold | $2.2\times10^{-3} - 2.2\times10^1$ | 0.0964 | Notch sweep (fast) |
| $\tilde{N}^{(\mathrm{low})}$ | $N^{(\mathrm{low})}/(\beta_N/\gamma_N)$ | Fractional low Notch threshold | $0.1 - 1.0$ | 0.1013 | Notch sweep (fast) |
| $\tilde{N}^{(\mathrm{high})}$ | $N^{(\mathrm{high})}/(\beta_N/\gamma_N)$ | Fractional high Notch threshold | $0.1 - 1.0$ | 0.3897 | Notch sweep (fast) |
| $\tilde{\sigma}$ | $\sigma/(\beta_D/\gamma_D)$ | Fractional Delta signalling threshold | $10^{-3} - 10^1$ | 0.0321 | Notch sweep (slow) |
| $\tilde{\epsilon}$ | $\epsilon/(\beta_N/\gamma_N)$ | Fractional Notch signalling threshold | $2.2\times10^{-3} - 2.2\times10^1$ | 0.1973 | Notch sweep (slow) |
| $\tilde{N}^{(\mathrm{low})}$ | $N^{(\mathrm{low})}/(\beta_N/\gamma_N)$ | Fractional low Notch threshold | $0.1 - 1.0$ | 0.1038 | Notch sweep (slow) |
| $\tilde{N}^{(\mathrm{high})}$ | $N^{(\mathrm{high})}/(\beta_N/\gamma_N)$ | Fractional high Notch threshold | $0.1 - 1.0$ | 0.3261 | Notch sweep (slow) |

Table M3: **Optimal parameter values obtained from parameter sweeps.** The ranges of parameter values tested during parameter sweep simulations over the non-dimensionalised Delta, Notch and Hh equations defined in eqs. (M1a-M3c), and the parameter values that optimise the robustness defined in eq. (M4) for each of these populations. Fast and slow Notch sweeps correspond to Delta and Notch degradation rates of $\gamma_D = \gamma_N = 2\times10^{-3}$ and $\gamma_D = \gamma_N = 2\times10^{-5}$, respectively.

### 6 Supplementary Figures

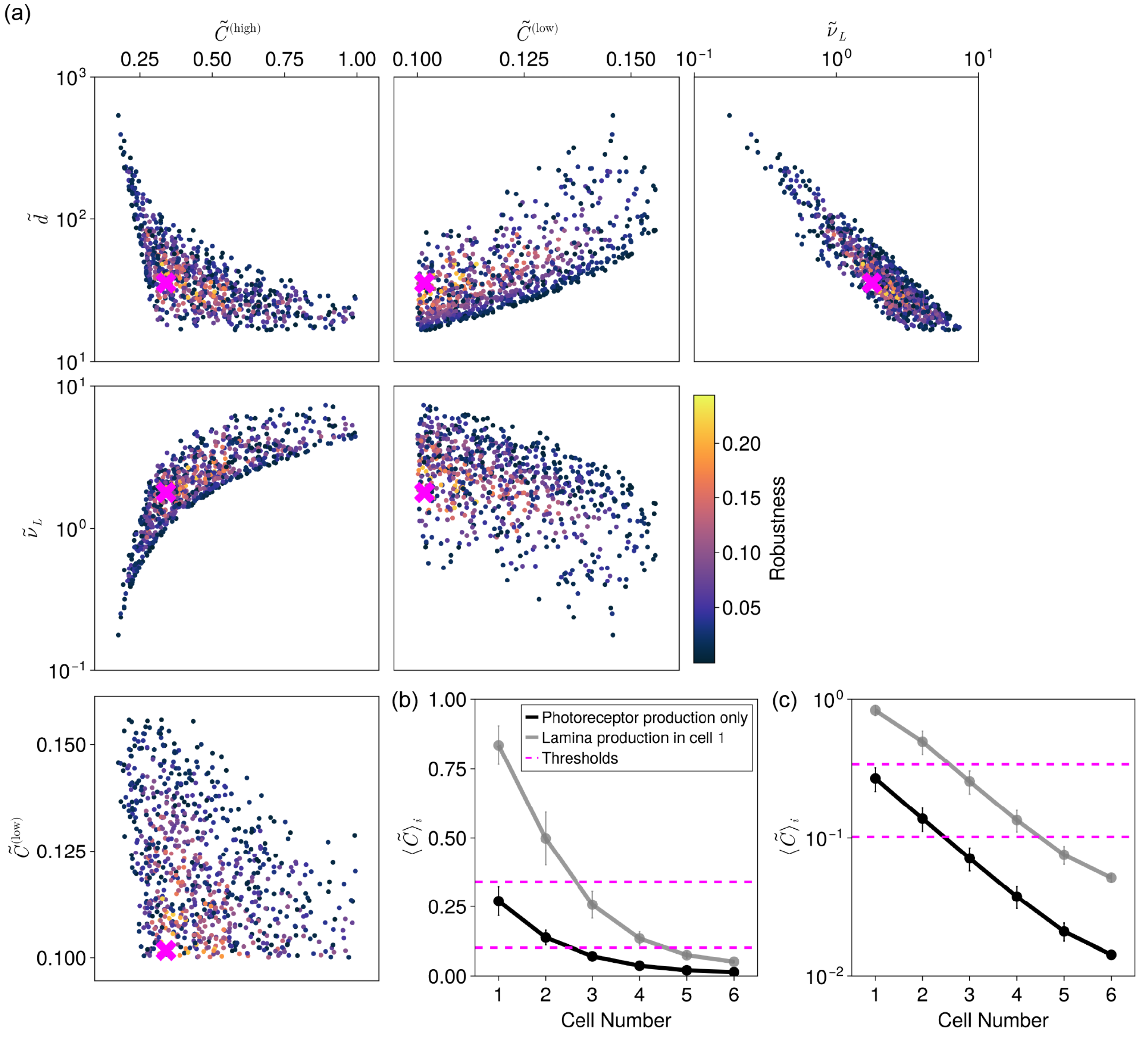

Figure M1: **Sweeping over Hh dynamical parameters to optimise robustness.** (a) Pair-wise plots of the dynamical parameters describing the Hh effective diffusivity $\tilde{d}$, the fractional local Hh production $\tilde{\nu}_L$, and the fractional Hh thresholds $\tilde{C}^{(\mathrm{low})}$ and $\tilde{C}^{(\mathrm{high})}$ that reproduce wild-type cell fate patterns. Colour indicates how 'well-separated' Notch levels are from their nearest thresholds, as quantified in eq. (M4). The magenta crosses indicate the parameters for the system with Hh levels that are the most well-separated from their nearest thresholds (plotted in (b)). All plots share the same colourbar. (b,c) Gradients of the average Hh concentration across each cell for the system with Hh levels that are the most well-separated from their nearest thresholds, for different values of the Hh source width relevant in wild-type experiments, plotted using a (b) linear or (c) logged y-axis. Error bars represent the standard deviations of the Hh concentration across each cell width.

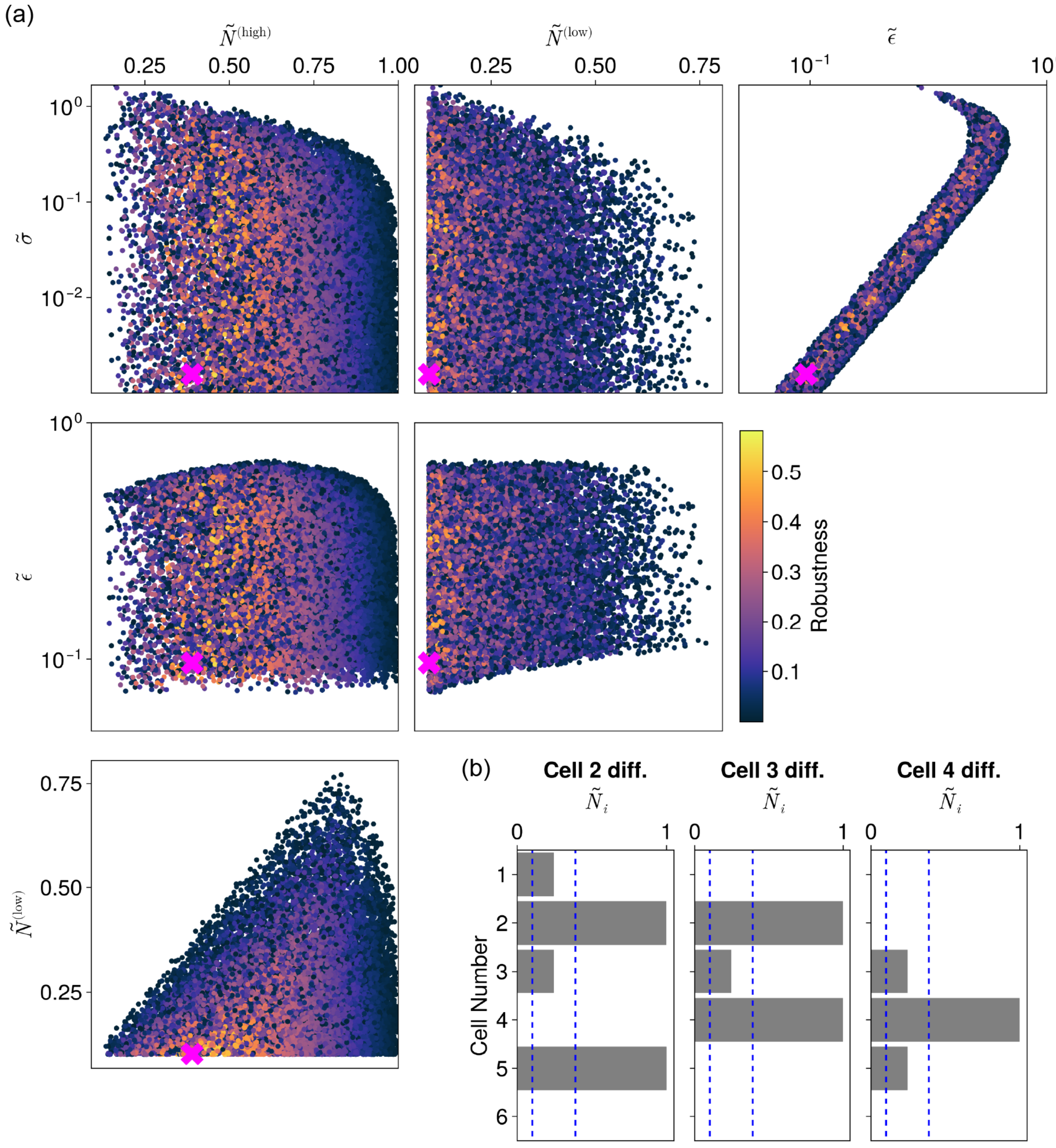

Figure M2: **Sweeping over Delta-Notch dynamical parameters to optimise Notch separation from thresholds.** (a) Pair-wise plots of the dynamical parameters describing the Delta and Notch signalling thresholds $\tilde{\sigma}$ and $\tilde{\epsilon}$, and the fractional Notch thresholds $\tilde{N}^{(\text{low})}$ and $\tilde{N}^{(\text{high})}$ that reproduce wild-type cell fate patterns. Colour indicates how 'well-separated' Notch levels are from their nearest thresholds, as quantified in eq. (M4). The magenta crosses indicate the parameters for the system with Notch levels that are the most well-separated from their nearest thresholds (plotted in (b)). All plots share the same colourbar. (b) Steady-state Notch profiles corresponding to the times at which cells in position $i$ differentiate. The blue dashed lines indicate the locations of the Notch thresholds $\tilde{N}^{(\text{low,high})}$.

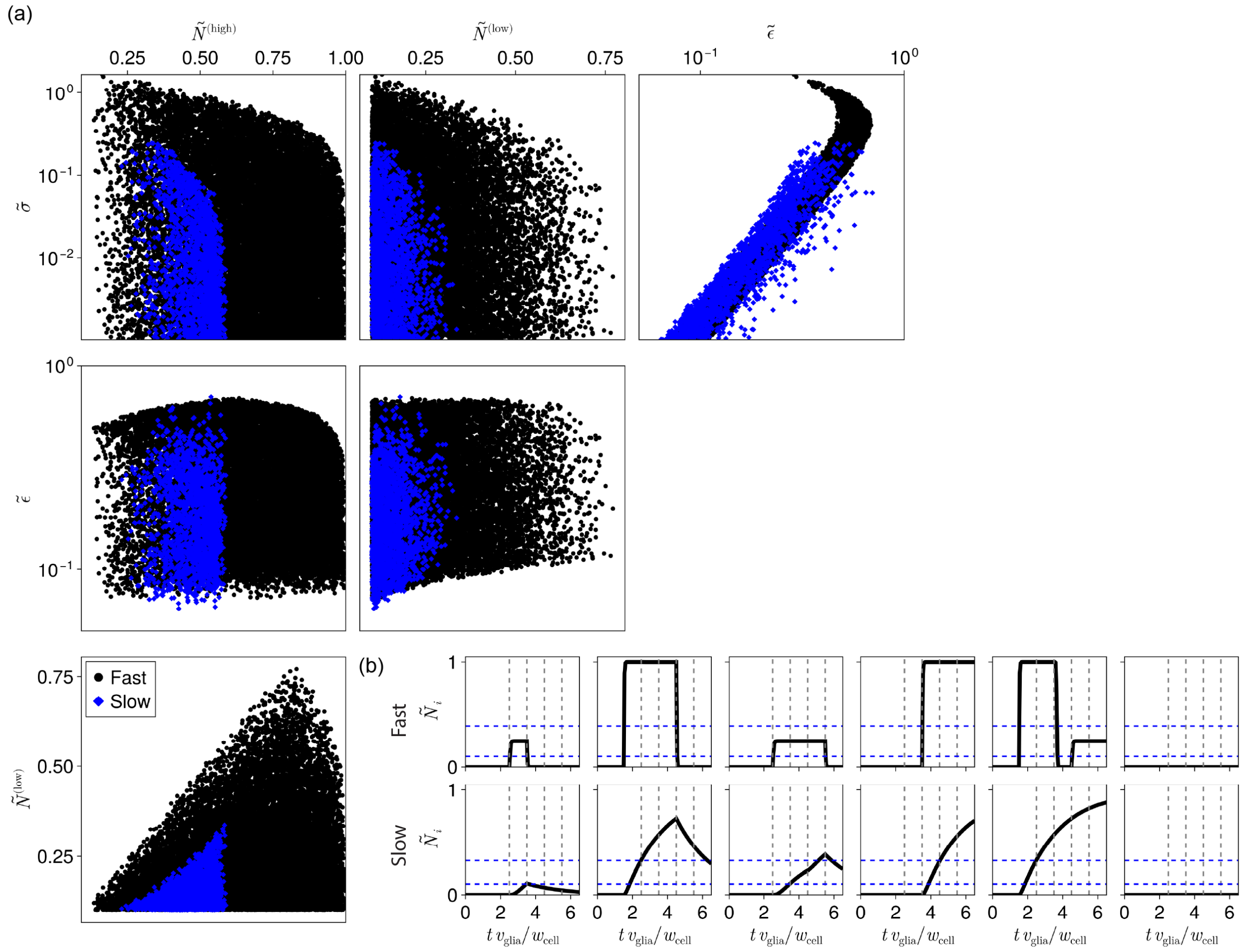

Figure M3: **Variation in Delta and Notch degradation rates constrain parameter space.** (a) Pairwise plots of the dynamical parameters describing the Delta and Notch signalling thresholds $\tilde{\sigma}$ and $\tilde{\epsilon}$, and the fractional Notch thresholds $\tilde{N}^{(\text{low})}$ and $\tilde{N}^{(\text{high})}$ that reproduce wild-type cell fate patterns for systems with fast ($\gamma_D = \gamma_N = 2 \times 10^{-3}$ s$^{-1}$) or slow ($\gamma_D = \gamma_N = 2 \times 10^{-5}$ s$^{-1}$) Delta and Notch degradation rates. (b) Comparison between the corresponding time series of the Notch concentrations in position-1 through -4 cells (grey vertical dashed lines indicate the times at which position-1 through -4 cells differentiate, in order; blue horizontal dashed lines indicate the locations of the Notch thresholds $\tilde{N}^{(\text{low},\text{high})}$).

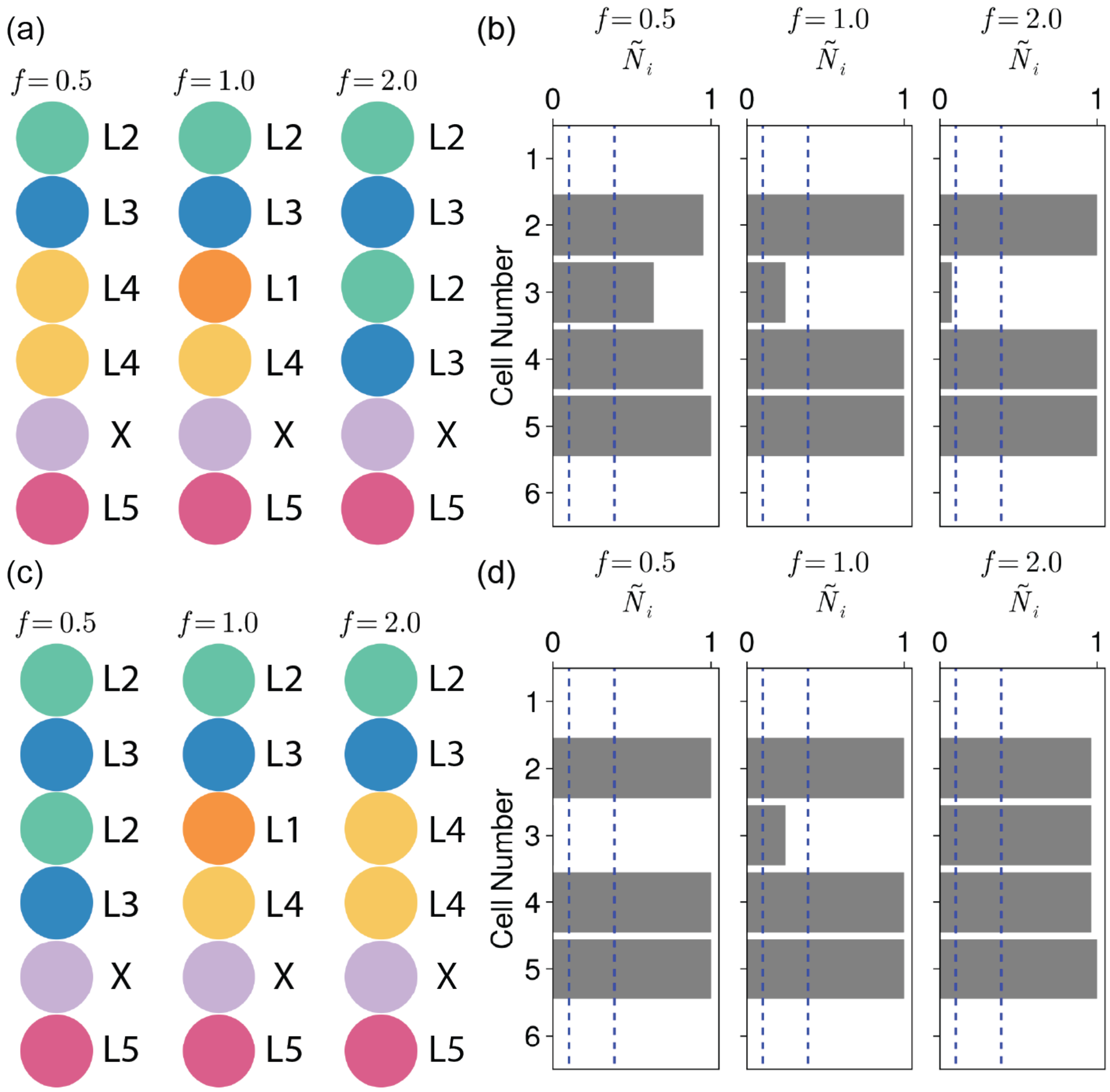

Figure M4: **The values of $\tilde{\sigma}$ and $\tilde{\epsilon}$ control graded Notch activity.** (a) The different phenotypes observed for the fractional Delta signalling threshold $\tilde{\sigma} = f\,\tilde{\sigma}^{(0)}$ for varying values of $f$, where $\tilde{\sigma}^{(0)}$ is the base value shown in table M1. (b) The corresponding Notch signalling levels in each cell at the time of cell fate specification (blue dashed lines indicate the locations of the Notch thresholds $\tilde{N}^{(\text{low,high})}$). (c) The different phenotypes observed for the fractional Notch signalling threshold $\tilde{\epsilon} = f\,\tilde{\epsilon}^{(0)}$ for varying values of $f$, where $\tilde{\epsilon}^{(0)}$ is the base value shown in table M1. (d) The corresponding Notch signalling levels in each cell at the time of cell fate specification.

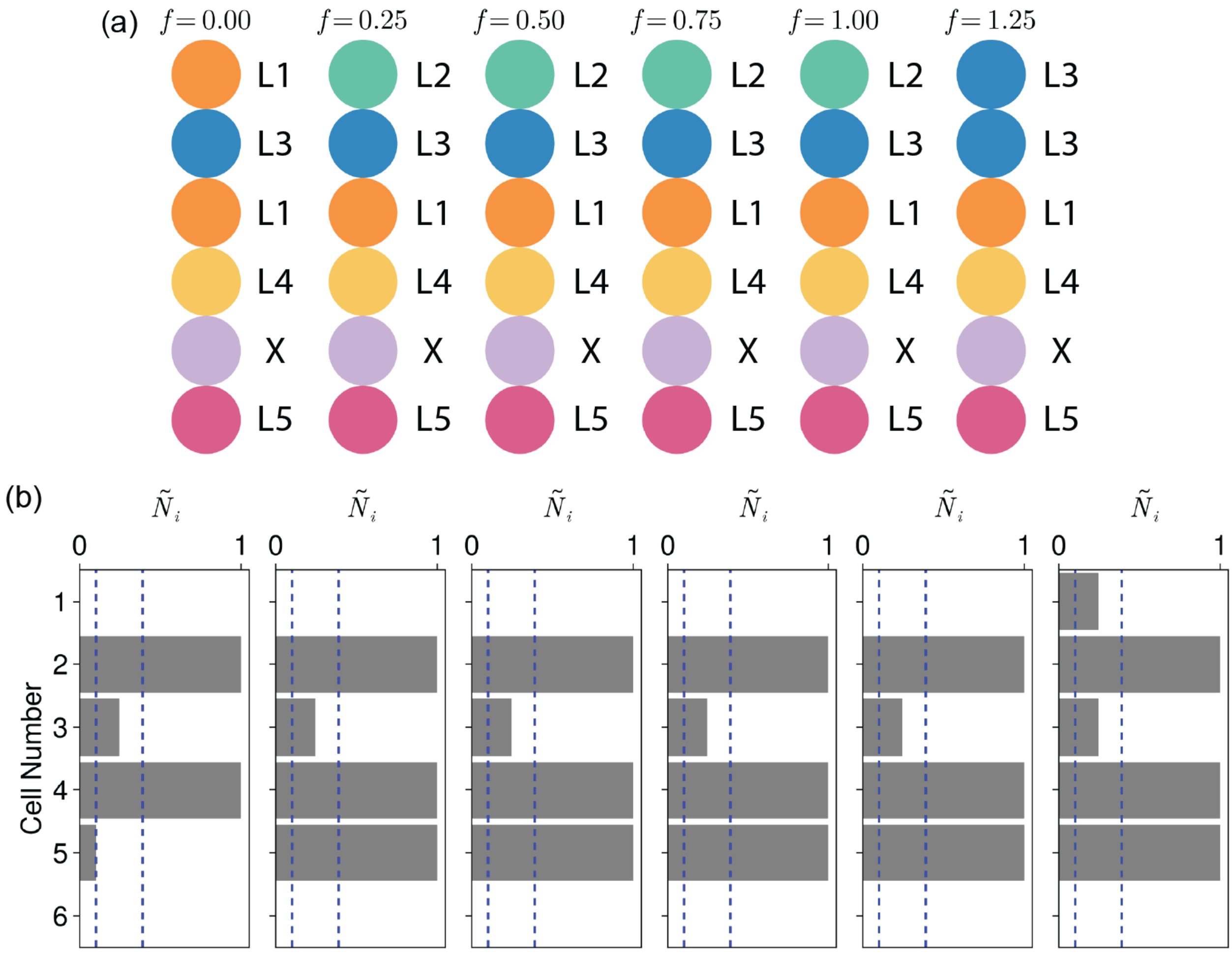

Figure M5: **Delays in making cell fate decisions affect phenotype.** (a) The different phenotypes observed when cell fate decisions are made after times $\tau_c = f\, w_{\text{cell}}/v_{\text{glia}}$ for varying values of $f$. The wild-type phenotype is only observed when $0.25 \leq \tau_{\text{delay}}^{(\text{fate})} \leq w_{\text{cell}}/v_{\text{glia}}$. (b) The corresponding Notch signalling levels in each cell at the time of cell fate specification (blue dashed lines indicate the locations of the Notch thresholds $\tilde{N}^{(\text{low,high})}$).

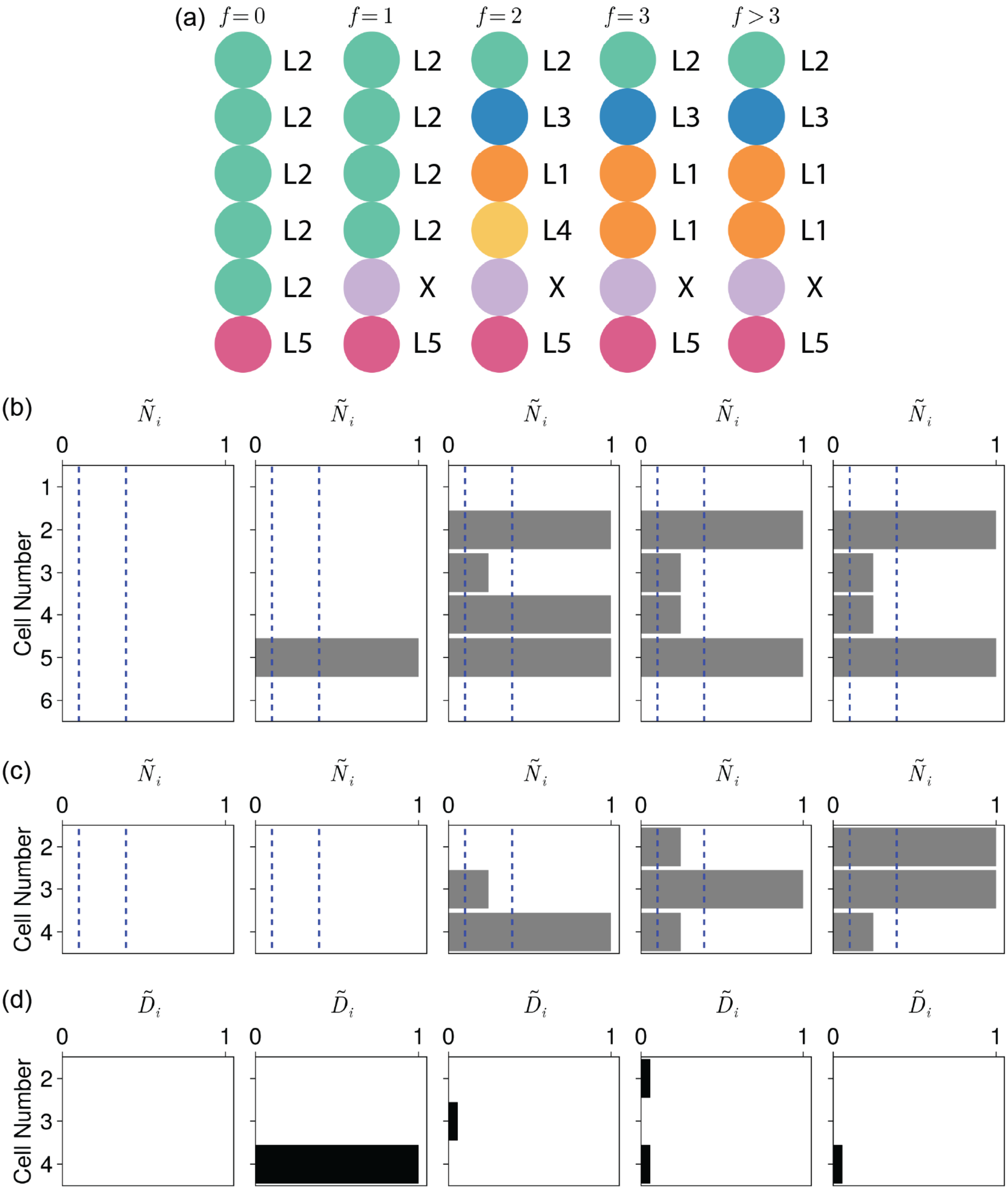

Figure M6: **Delays in switching off Delta production affect phenotype.** (a) The different phenotypes observed when Delta production is switched off after times $\tau_{\mathrm{DN}} = f\, w_{\mathrm{cell}}/v_{\mathrm{glia}}$ for varying values of $f$. The wild-type phenotype is only observed when $f = 2$. (b) The corresponding Notch signalling levels in each cell at the time of cell fate specification (blue dashed lines indicate the locations of the Notch thresholds $\tilde{N}^{(\mathrm{low,high})}$). (c,d) The (c) Notch (grey) and (d) Delta (black) levels in position-2 through -4 cells at the time when the position-4 cell differentiates.

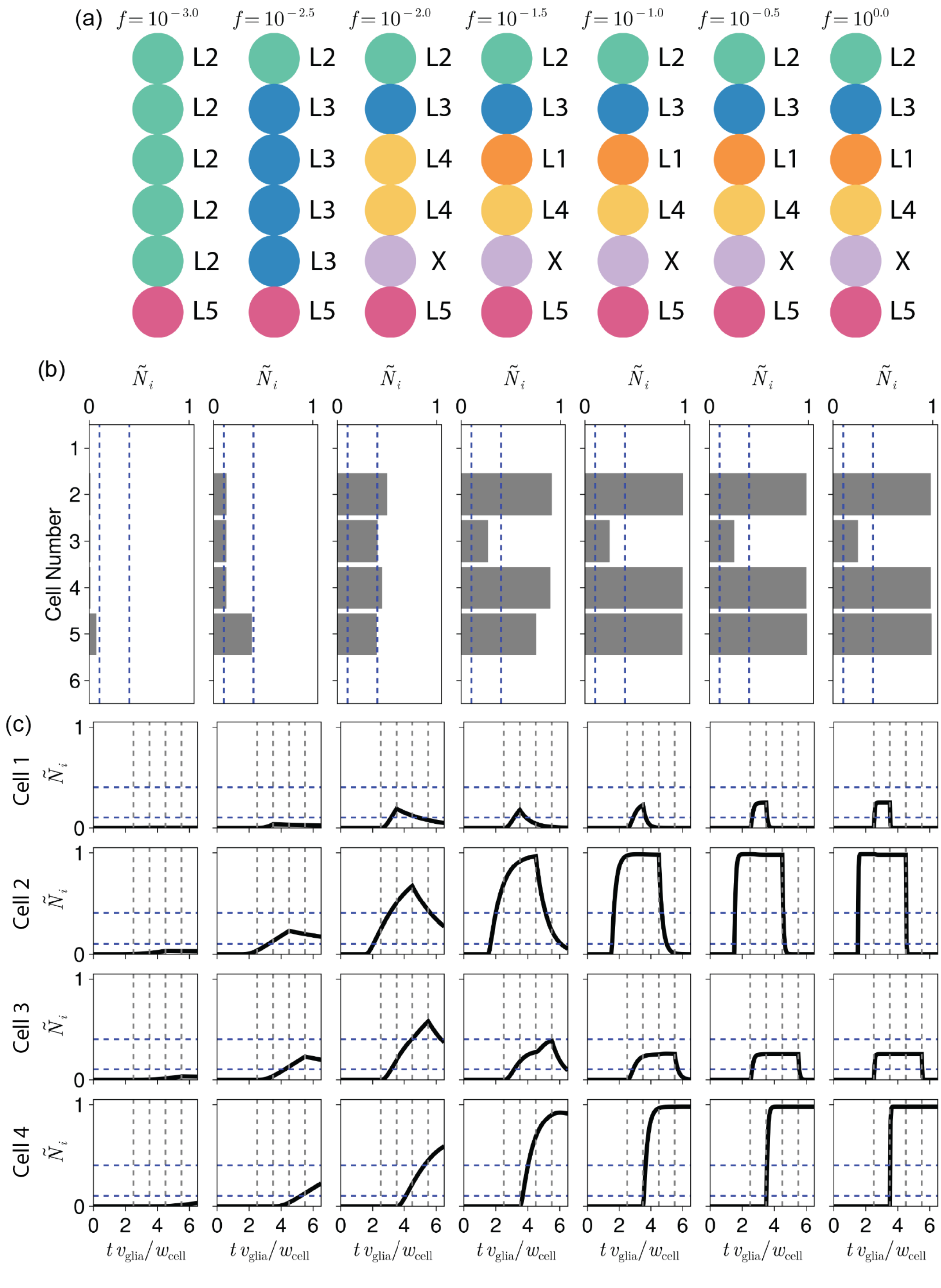

Figure M7: **Variation in Delta and Notch degradation rates affect phenotype.** (a) The different phenotypes observed for the Delta and Notch degradation rates $\gamma_D = f\,\gamma_D^{(0)}$ and $\gamma_N = f\,\gamma_N^{(0)}$ for varying values of $f$, where $\gamma_D^{(0)}$ and $\gamma_N^{(0)}$ are the base values shown in table M1. (b) The corresponding Notch signalling levels in each cell at the time of cell fate specification (blue dashed lines indicate the locations of the Notch thresholds $\tilde{N}^{(\text{low,high})}$). (c) The corresponding time series of the Notch concentration in position-1 through -4 cells (grey vertical dashed lines indicate the times at which position-1 through -4 cells differentiate, in order; blue horizontal dashed lines indicate the locations of the Notch thresholds $\tilde{N}^{(\text{low,high})}$).

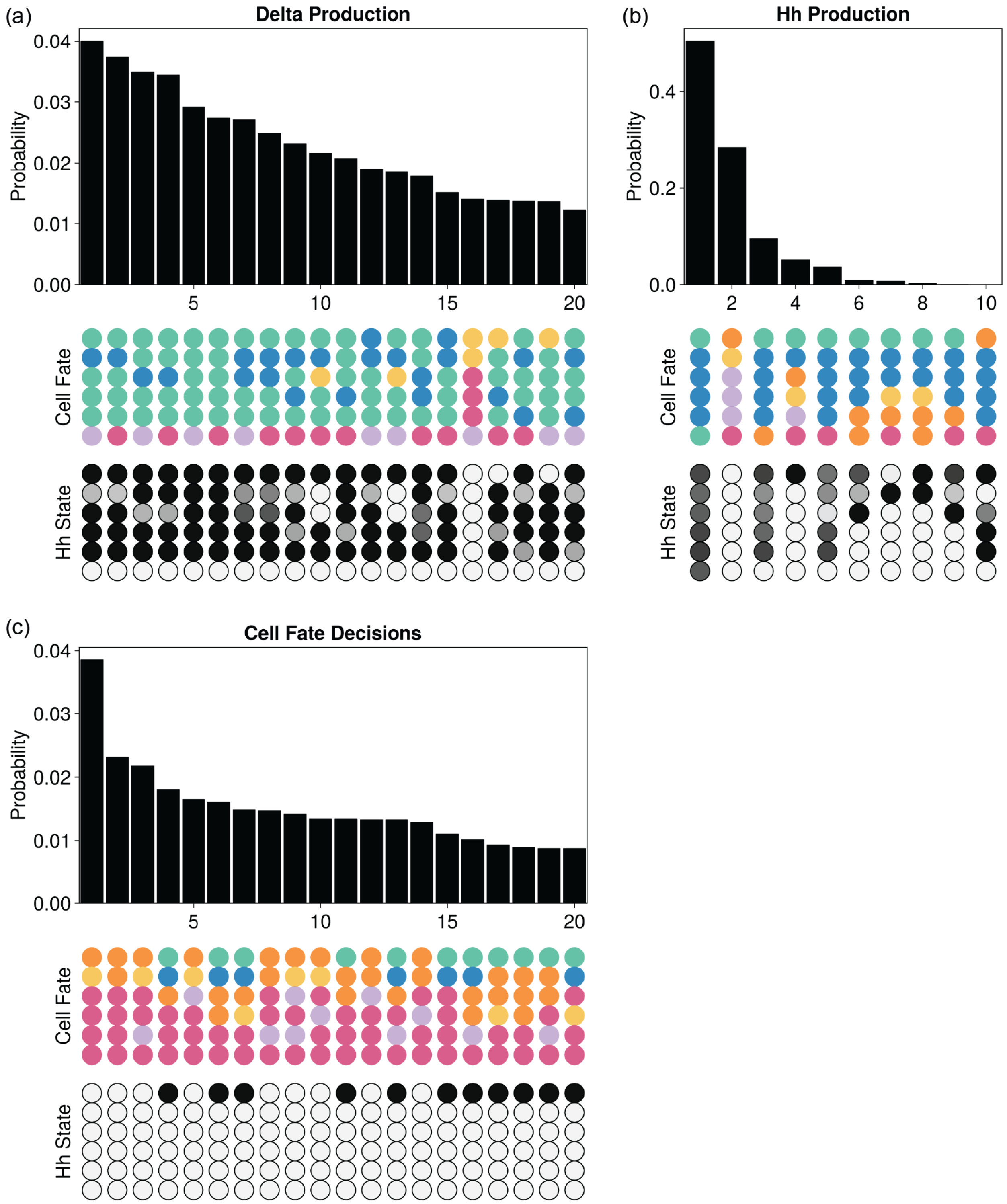

Figure M8: **Extended distributions of the cell fate phenotypes obtained by randomising system timescales.** Bar charts representing the probabilities of achieving specified cell fates following the randomisation of the times of: (a) Delta production, (b) lamina Hh production, and (c) cell fate specification. The most abundant 20 cell fates are shown for the Delta production and cell fate decisions plots, but only 10 cell fates were obtained when varying the timing of Hh production. Below each distribution are the corresponding cell fates (cell fate colours are the same as in Fig. 4E in the main text) and average final Hh production state (black is on, white is off, and grey represents an average that lies between those extremes).
